# Natural Selection is Unlikely to Explain Why Species Get a Thin Slice of *π*

**DOI:** 10.1101/2021.02.03.429633

**Authors:** Vince Buffalo

## Abstract

Neutral theory predicts that genetic diversity increases with population size, yet observed levels of diversity across metazoans vary only two orders of magnitude while population sizes vary over several. This unexpectedly narrow range of diversity is known as Lewontin’s Paradox of Variation (1974). While some have suggested selection constrains diversity, tests of this hypothesis seem to fall short. Here, I revisit Lewontin’s Paradox to assess whether current models of linked selection are capable of reducing diversity to this extent. To quantify the discrepancy between pairwise diversity and census population sizes across species, I combine previously-published estimates of pairwise diversity from 172 metazoan taxa with estimates of census sizes. Using phylogenetic comparative methods, I show this relationship is significant accounting for phylogeny, but with high phylogenetic signal and evidence that some lineages experience shifts in the evolutionary rate of diversity deep in the past. Additionally, I find a negative relationship between recombination map length and census size, suggesting abundant species have less recombination and experience greater reductions in diversity due to linked selection. However, I show that even using strong selection parameter estimates, models of linked selection are unlikely to explain the observed relationship between diversity and census sizes across species.

---

A longstanding mystery in evolutionary genetics is that the observed levels of genetic variation across sexual species are confined to an unexpectedly narrow range. Under neutral theory, the average number of nucleotide differences between lineages (pairwise diversity, *π*) is determined by the balance of new mutations and their loss by genetic drift (Kimura and Crow 1964; Malécot 1948; Wright 1931). In particular, the expected diversity at neutral sites in a panmictic population of *N*_*c*_ diploids is expected to be *π* ≈ 4*N*_*c*_*µ*, where *µ* is the per basepair per generation mutation rate. Given that metazoan germline mutation rates only differ 10-fold (10^−8^–10^−9^, Kondrashov and Kondrashov 2010; Lynch 2010), and census sizes vary over several orders of magnitude, one would expected under neutral theory that heterozygosity should also vary over several orders of magnitude. However, early allozyme surveys revealed that heterozygosity levels across a wide range of species varied just an order of magnitude (Lewontin 1974, p. 208); this is known as Lewontin’s “Paradox of Variation”. With modern sequencing-based estimates of *π* across taxa ranging over only three orders of magnitude (0.01–10%, Leffler et al. 2012), Lewontin’s paradox has persisted unresolved through the genomics era.

From the beginning, explanations for Lewontin’s Paradox have been framed in terms of the neutralist–selectionist controversy (Gillespie 1991, 2001; Kimura 1984; Lewontin 1974). The neutralist view is that beneficial alleles are sufficiently rare and deleterious alleles removed sufficiently quickly, that levels of genetic diversity are shaped predominantly by genetic drift and mutation (Kimura 1984). Specifically, *non-selective* processes decouple the effective population size implied by observed levels of diversity 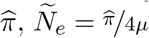, from the census size, *N*_*c*_. By contrast, the selectionist view is that the direct and indirect effects of linked selection suppress diversity levels across taxa, specifically because the impact of linked selection is greater in large populations. Undoubtedly, these opposing views represent a false dichotomy, as population genomic studies have uncovered both complex demographic histories that impact diversity within a species (e.g. Palkopoulou et al. 2015; Zhao et al. 2013) and evidence that selection depresses genome-wide diversity (e.g. Aguade et al. 1989; Begun and Aquadro 1992; Elyashiv et al. 2016; McVicker et al. 2009).

## Possible Explanations of Lewontin’s Paradox

A resolution of Lewontin’s Paradox would involve a mechanistic description and quantification of the evolutionary processes that prevent diversity from scaling with census sizes across species. This would necessarily connect to the broader literature on the empirical relationship between diversity and population size (Frankham 1996; Leroy et al. 2021; Nei and Graur 1984; Soulé 1976), and the ecological and life history correlates of genetic diversity (Nevo 1978; Nevo et al. 1984; Powell 1975). Three categories of processes stand out as potentially capable of decoupling census sizes from diversity: non-equilibrium demography, variance and skew in reproductive success, and selective processes.

It has long been appreciated that effective population sizes are typically less than census population sizes, tracing back to early debates between R.A. Fisher and Sewall Wright (Fisher and Ford 1947; Wright 1948). Possible causes of this divergence between effective and census population sizes include demographic history (e.g. population bottlenecks), extinction and recolonization dynamics, or the breeding structure of populations (e.g. the variance in reproductive success and population substructure). Early explanations for Lewontin’s Paradox suggested bottlenecks during the last glacial maximum severely reduced population sizes (Kimura 1984; Nei and Graur 1984; Ohta and Kimura 1973), and emphasized that large populations recover to equilibrium diversity levels more slowly (Nei and Graur 1984, Kimura 1984 p. 203-204). Another explanation is that cosmopolitan species repeatedly endure extinction and recolonization events, which reduces effective population size (Maruyama and Kimura 1980; Slatkin 1977).

While chance demographic events like bottlenecks and recent expansions have long-term impacts on diversity (since mutation-drift equilibrium is reached on the order of size of the population), characteristics of the breeding structure such as high variance (*V*_*w*_) or skew in reproductive success continuously suppress diversity below the levels predicted by the census size (Wright 1938). For example, in many marine animals, females are highly fecund, and dispersing larvae face extremely low survivorship, leading to high variance in reproductive success (Hauser and Carvalho 2008; Hedgecock and Pudovkin 2011; Waples et al. 2018, 2013). Such “sweepstakes” reproductive systems can lead to remarkably small ratios of effective population size to census population size (e.g. ^*N*^_*e*_/^*N*^_*c*_ can range from 10^−6^–10^−2^), since ^*N*^_*e*_/^*N*^ ≈ ^1^/^*V*^_*w*_ (Hedgecock 1994; Nunney 1993, 1996; Wright 1938), and require multiple-merger coalescent processes to describe their genealogies (Eldon and Wakeley 2006). Overall, these reproductive systems diminish the diversity in many species, but seem unlikely to explain Lewontin’s Paradox broadly across metazoans.

Alternatively, selective processes, and in particular the indirect effects of selection on linked neutral variation, could explain the observed narrow range of diversity. The earliest mathematical model of hitchhiking was proffered as a explanation of Lewontin’s Paradox (Maynard Smith and Haigh 1974). Since, empirical observations have demonstrated that linked selection shapes patterns of genome-wide diversity, as evidenced by the correlation between recombination and diversity in a variety of species (Aguade et al. 1989; Begun and Aquadro 1992; Cai et al. 2009; Cutter and Payseur 2003; Stephan and Langley 1998). Theoretic work to explain this pattern has considered diversity under a steady influx of new beneficial mutations (recurrent hitchhiking; Stephan 1995; Stephan et al. 1992), and purifying selection against new deleterious mutations (background selection, BGS; Charlesworth et al. 1993; Hudson and Kaplan 1995; Hudson and Kaplan 1994; Nordborg et al. 1996). Indeed, empirical work indicates background selection diminishes diversity around genic regions in a variety of species (Charlesworth 1996; Hernandez et al. 2011; McVicker et al. 2009), and now efforts have shifted towards teasing apart the effects of positive and negative selection on genomic diversity (Elyashiv et al. 2016).

An important class of theoretic selection models pertaining to Lewontin’s Paradox are recurrent hitchhiking models that decouple diversity from the census population size. These models predict diversity when strongly selected beneficial mutations regularly enter and sweep through the population, trapping lineages and forcing them to coalesce (Gillespie 2000; Kaplan et al. 1989). In general, decoupling occurs under these hitchhiking models when the rate of coalescence due to selection is much greater than the rate of neutral coalescence (e.g. Coop and Ralph 2012, equation 22). Under other linked selection models, the resulting effective population size is proportional to population size, and thus these models cannot decouple diversity, all else equal. For example, models of background selection and polygenic fitness variation predict diversity is proportional to population size, mediated by the total recombination map length and the deleterious mutation rate or fitness variation (Charlesworth et al. 1993; Nicolaisen and Desai 2012; Nordborg et al. 1996; Robertson 1961; Santiago and Caballero 1995, 1998).

## Recent Approaches Towards Solving Lewontin’s Paradox

Recently, Corbett-Detig et al. (2015) used population genomic data to estimate the reduction in diversity due to background selection and hitchhiking across 40 species, and showed the impact of selection increases with two proxies of census population size, species range and with body size. They argued this is evidence that selection could explain Lewontin’s Paradox; however, in a reanalysis, Coop (2016) demonstrated that the observed scale of these reductions is insufficient to explain the orders-of-magnitude shortfall between observed and expected levels of diversity across species. Other recent work has found that certain life history characteristics related to parental investment, such as propagule size, are good predictors diversity in animals (Chen et al. 2017; Romiguier et al. 2014). Nevertheless, while these diversity correlates are important clues, they do not propose a mechanism by which these traits act to constrain diversity within a few orders of magnitude.

Here, I revisit Lewontin’s Paradox by integrating a variety of data sets and assessing the predicted reductions in diversity under different selection models. Prior surveys of genetic diversity either lacked census population size estimates, used allozyme-based measures of heterozygosity, or included fewer species. To address these shortcomings, I first estimate census sizes by combining predictions of population density based on body size with ranges estimated from geographic occurrence data. Using these estimates, I quantify the relationship between census size and previously-published genomic diversity estimates across 172 metazoan taxa within nine phyla, to provide a sense of the scale of the divergence between *π* and *N*_*c*_ that leads to Lewontin’s Paradox.

Past work looking at the relationship between *π* and *N*_*c*_ has been unable to fully account for phylogenetic non-independence across taxa (Felsenstein 1985). To address this shortcoming, I use phylogenetic comparative methods (PCMs) with a synthetic time-calibrated phylogeny to account for shared phylogenetic history. Moreover, it is disputed whether considering phylogenetic non-independence is necessary in population genetics, since coalescent times are much less than divergence times (Lynch 2011; Whitney and Garland 2010). Using PCMs, I address this by estimating the degree of phylogenetic signal in the diversity census size relationship, and investigating how these traits evolve along the phylogeny.

Finally, I explore whether the predicted reductions of diversity under background selection and recurrent hitchhiking are sufficiently strong to resolve Lewontin’s Paradox. These predicted reductions in diversity across species are estimated using strong selection parameters from *Drosophila melanogaster*, a species known to be strongly affected by linked selection. Given the effects of linked selection are mediated by recombination map length, I investigate how recombination map lengths vary with census population size using data from a previously-published survey (Stapley et al. 2017). I find map lengths are typically shorter in large–census-size species, increasing the effects of linked selection in these species, which might further decouple diversity from census size. Still, I find the combined impact of these selection models with available parameter estimates falls short in explaining Lewontin’s Paradox, and discuss future avenues through which the Paradox of Variation could be fully resolved.

## Results

### Estimates of Census Population Size

A major impediment in quantifying Lewontin’s Paradox has been estimating census population sizes across many taxa, especially for extremely abundant, cosmopolitan species that define the upper limit of ranges. Previous work has surveyed the literature for census size estimates (Frankham 1996; Nei and Graur 1984; Soulé 1976), or used range, body size, or qualitative categories as proxies for census size (Corbett-Detig et al. 2015; Leffler et al. 2012). To quantify the relationship between genomic estimates of diversity and census population sizes, I first approximate census population sizes for 172 metazoan taxa (Figure 1). My approach predicts population densities from body sizes using a previously-observed linear relationship that holds across metazoans (see Supplementary Figure A4; Damuth 1981, 1987). Then, from geographic occurrence data, I estimate range sizes. Finally, I estimate population size as the product of these predicted densities and range estimates (see Methods: *Macroecological Estimates of Population Size*). Note that the relationship between population density and body size is driven by energy budgets, and thus reflects macroecological equilibria (Damuth 1987). Consequently, population sizes are underestimated for taxa like humans and their domesticated species, and overestimated for species with anthropogenically reduced densities or fragmented ranges. For example, the population size of *Lynx lynx* is likely around 50,000 (IUCN 2020) which is around two orders of magnitude smaller than my estimate. Additionally, the range size estimates do not consider whether an area has unsuitable habitat, and thus may be overestimated for species with particular niches or patchy habitats. While these methods to estimate census size are crude and approximate, they can be efficiently calculated for numerous taxa and are sufficient to estimate the scale of Lewontin’s Paradox (see Supplementary Information *Population Size Validation* for more on validation based on biomass and other approaches).

**Figure 1:**
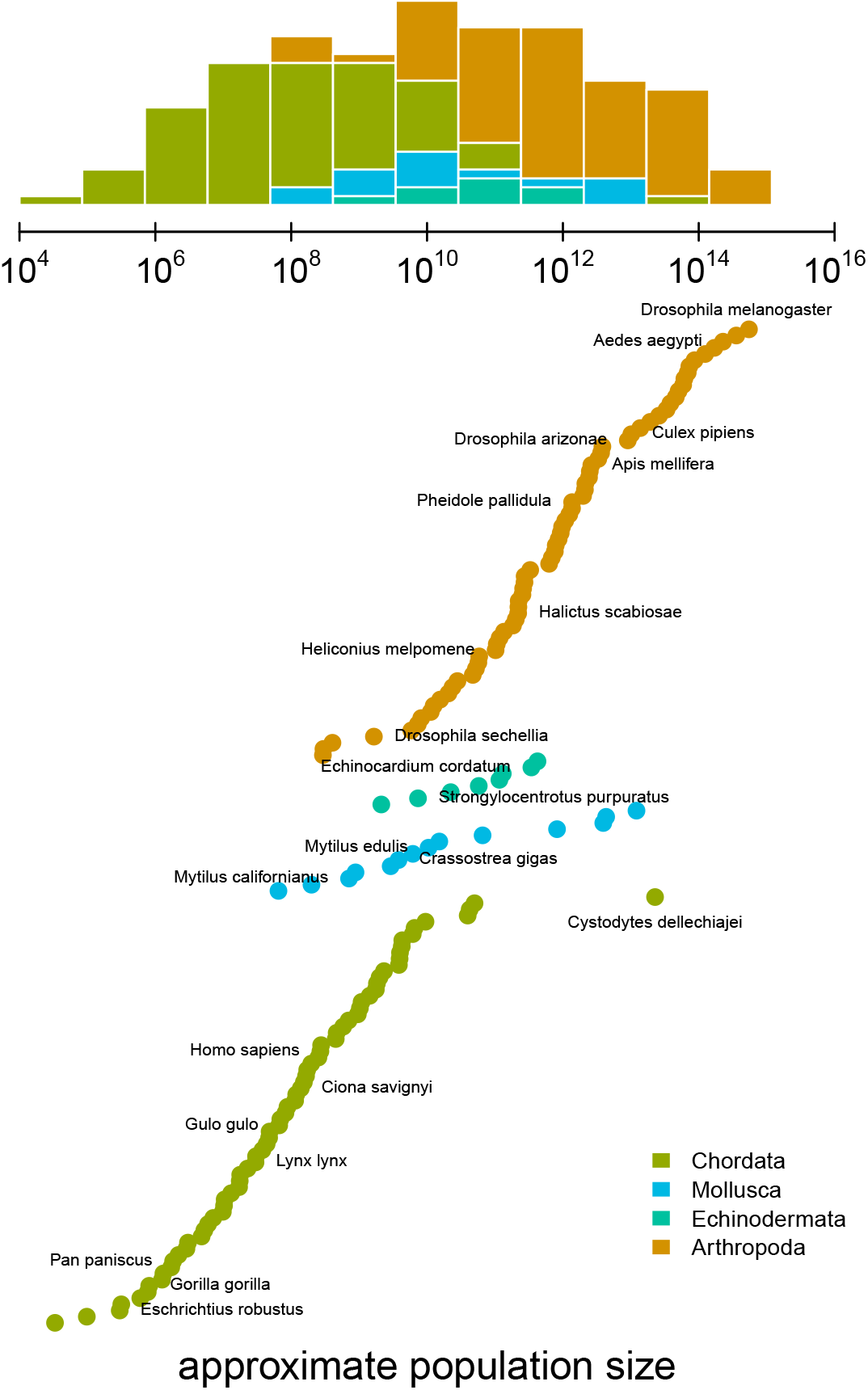
The distribution of approximate census population sizes estimated by this study. Some phyla containing few species were excluded for clarity.

### Quantifying Lewontin’s Paradox

To determine which ecological or evolutionary processes could decouple diversity from census population size, we first need to quantify this relationship across a wide variety of taxa. Previous work has found there is a significant relationship between heterozygosity and the logarithm of population size or range size, but these studies relied on heterozygosity measured from allozyme data (Frankham 1996; Leffler et al. 2012; Nei and Graur 1984; Soulé 1976). Here, I confirm these findings using pairwise diversity estimates from genomic sequence data and the estimated census sizes (Figure 2). The pairwise diversity estimates are from three sources: Leffler et al. (2012), Corbett-Detig et al. (2015), and Romiguier et al. (2014), and are predominantly from either synonymous or non-coding DNA (see Methods: *Diversity and Map Length Data*). Overall, an ordinary least squares (OLS) relationship on a log-log scale fits the data well (Figure 2). The OLS slope estimate is significant and implies a 13% percent increase in differences per basepair for every order of magnitude census size grows (95% confidence interval [12%, 14%], adjusted *R*^2^ = 0.26; see also the OLS fit per-phyla, Figure 1-figure supplement A6).

**Figure 2:**
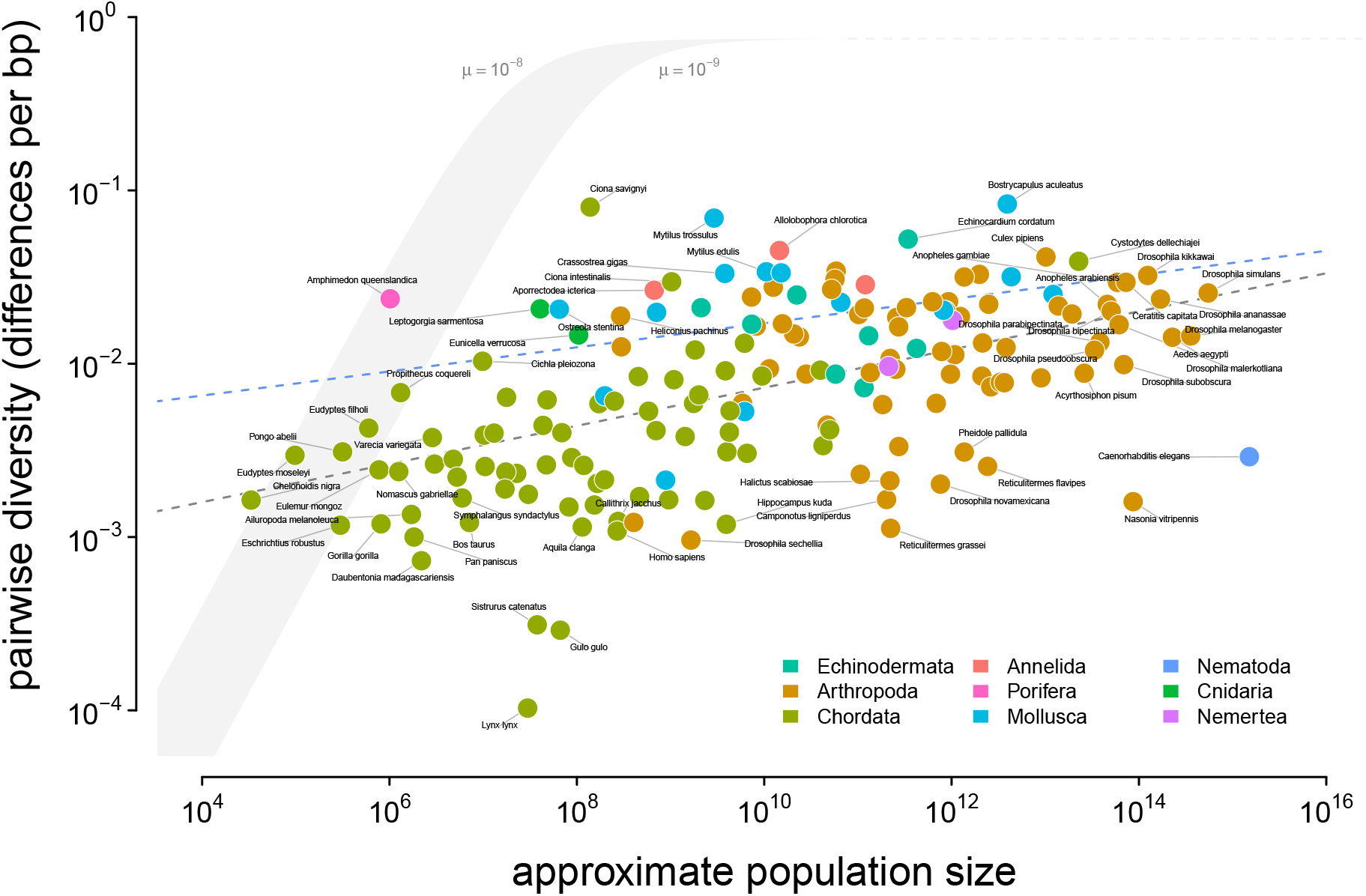
An illustration of Lewontin’s Paradox of Variation. Pairwise diversity (data from Leffler et al. 2012, Corbett-Detig et al. 2015, and Romiguier et al. 2014), which varies around three orders of magnitude, shows a weak relationship with approximate population size, which varies over 12 orders of magnitude. The shaded curve shows the range of expected neutral diversity if *N*_*e*_ were to equal *N*_*c*_ under the fouralleles model, log_10_(*π*) = log_10_(*θ*) − log_10_ (1 + ^4*θ*^/_3_) where *θ* = 4*N*_*c*_*µ*, for two mutation rates, *µ* = 10^−8^ and *µ* = 10^−9^, and the light gray dashed line represents the maximum pairwise diversity under the four alleles model. The dark gray dashed line is the OLS regression fit, and the blue dashed line is the regression fit using a phylogenetic mixed-effects model. Points are colored by phylum. The species *Equus ferus przewalskii* (*N*_*c*_ ≈ 10^3^ and *π* = 3.6 × 10^−3^) was an outlier and excluded from this figure for visual clarity.

Notably, this relationship has few outliers and is relatively homoscedastic. This is in part because of the log-log scale, in contrast to previous work (Nei and Graur 1984; Soulé 1976); see Supplementary Figure A5 for a version on a log-linear scale. However, it is noteworthy that few taxa have diversity estimates below 10^−3.5^ differences per basepair. Those that do, lynx (*Lynx Lynx*), wolverine (*Gulo gulo*), and Massasauga rattlesnake (*Sistrurus catenatus*) face habitat fragmentation and declining population sizes. These three species are all in the IUCN Red List, but are listed as least concern (though their presence in the Red List indicates they are of conservation interest). In Supplementary Information Section *Diversity and IUCN Red List Status*, I explore more about the relationships between IUCN Red List status, diversity, and population size.

### Phylogenetic Non-Independence and the Population Size Diversity Relationship

One limitation of using ordinary least squares is that shared phylogenetic history can create correlation structure in the residuals, which violates an assumption of the regression model (Felsenstein 1985; Revell 2010). To address this shortcoming, I fit this relationship using a phylogenetic mixed-effects model, investigated whether there is a signal of phylogenetic non-independence, estimated the continuous trait values on the phylogeny, and explored how diversity and population size evolve. Prior population genetic comparative studies have lacked time-calibrated phylogenies and assumed unit branch lengths (Whitney and Garland 2010), a shortcoming that has drawn criticism (Lynch 2011). Here, I use a synthetic time-calibrated phylogeny created from the DateLife project (O’Meara et al. 2020) to account for shared phylogenetic history (see Methods: *Phylogenetic Comparative Methods*).

Using a phylogenetic mixed-effects model (Hadfield and Nakagawa 2010; Lynch 1991; Villemereuil and Nakagawa 2014) implemented in Stan (Carpenter et al. 2017; Stan Development Team 2020), I estimated the linear relationship between diversity and population size (on a log-log scale) accounting for phylogeny, for the 166 taxa without missing data and present in the synthetic chronogram. As with the linear regression, this relationship was positive and significant (95% credible interval 0.03, 0.11), though somewhat attenuated compared to the OLS estimates (Figure 3B). Since the population size estimates are based on range and body mass, they are essentially a composite trait; fitting phylogenetic mixed-effects models separately on body mass and range indicates these have significant negative and positive effects, respectively (Supplementary Figure A7; see also Supplementary Figure A1 for the relationship between diversity and the range categories of Leffler et al. 2012).

**Figure 3:**
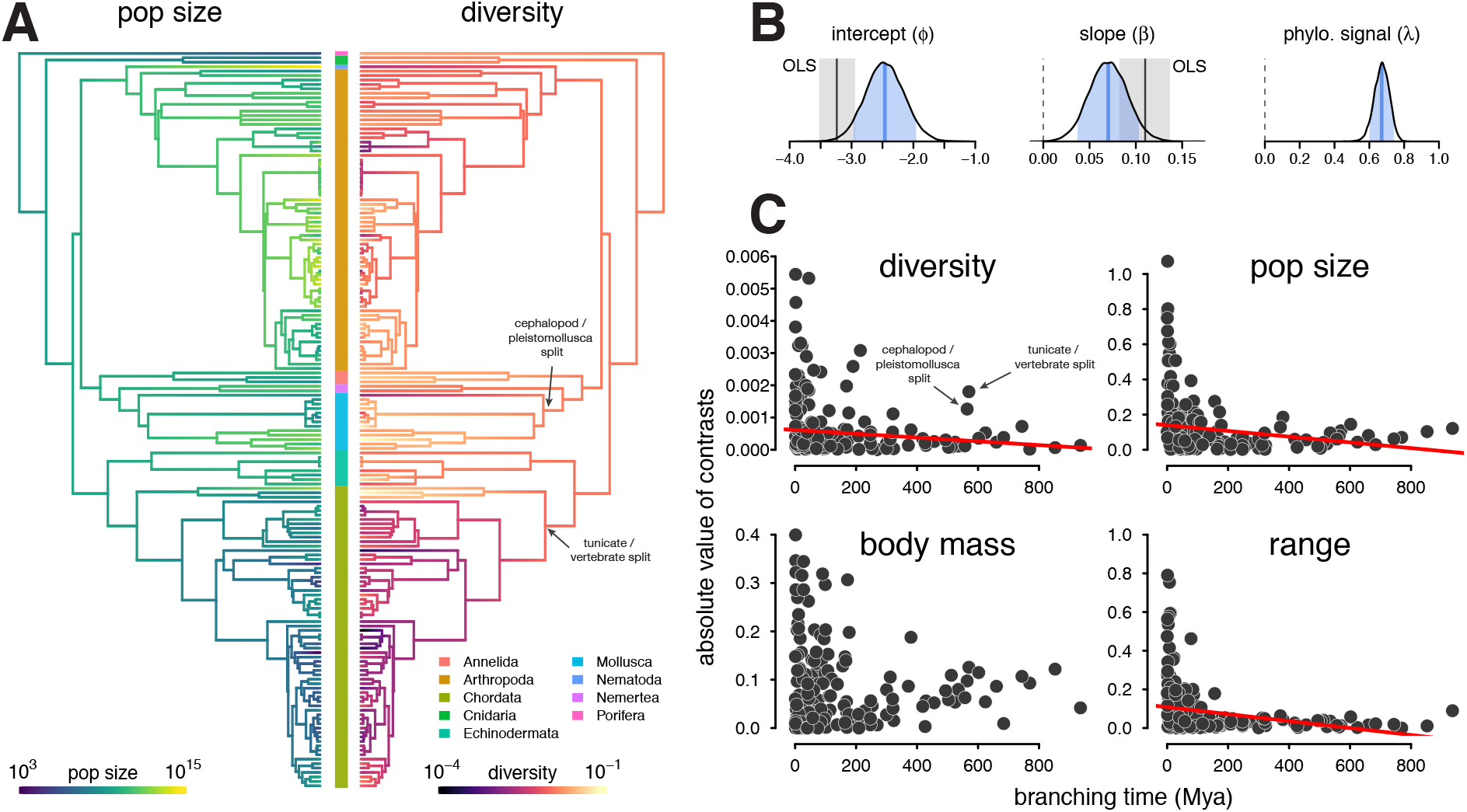
(A) The ancestral continuous trait estimates for the population size and diversity (differences per bp, log scaled) across the phylogeny of 166 taxa. The phyla of the tips are indicated by the color bar in the center. (B) The posterior distributions of the intercept, slope, and phylogenetic signal (*λ*, Villemereuil and Nakagawa 2014) of the phylogenetic mixed-effects model of diversity and population size (log scaled). Also shown are the 90% credible interval (light blue shading), posterior mean (blue line), OLS estimate (gray solid line), and bootstrap OLS confidence intervals (light gray shading). (C) The node-height tests of diversity, population size, and the two components of the population size estimates, body mass, and range (all traits on log scale before contrast was calculated). Each point shows the standardized phylogenetic independent contrast and branching time for a pair of lineages. Red lines are robust regression estimates (and are only shown for statistically significant relationships at the *α* = 0.05 level). Note that some outlier pairs with very high phylogenetic independent contrasts were excluded (in all cases, these outliers were in the genus *Drosophila*).

With the phylogenetic mixed-effects model, I also estimated the variance of the phylogenetic effect 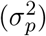 and the residual variance 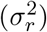, which can be used to estimate a measure of the phylogenetic signal, 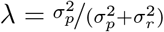 (Lynch 1991; Villemereuil and Nakagawa 2014; see Freckleton et al. 2002 for a comparison to Pagel’s *λ*). If the relationship between diversity and population size was free of shared phylogenetic history, *λ* = 0 and all the variance could be explained by evolution on the tips; this is analogous to Lynch’s conjecture that coalescent times should be free of phylogenetic signal (2011). In the relationship between population size and diversity, the posterior mean of *λ* = 0.67 (90% credible interval [0.58, 0.75]) indicates that the majority of the variance perhaps might be due to shared phylogenetic history (Figure 3B).

This high degree of phylogenetic signal suggests Gillespie’s (1991) concern that the *π*–*N*_*c*_ relationship was driven by chordate-arthropod differences may be valid. A visual inspection of the estimated ancestral continuous values for diversity and population size on the phylogeny indicates the high phylogenetic signal seems to be driven in part by chordates having low diversity and small population sizes compared to non-chordates (Figure 3A). This problem resembles Felsenstein’s worst-case scenario (Felsenstein 1985; Uyeda et al. 2018), where a singular event on a lineage separating two clades generates a spurious association between two traits.

To investigate whether clade-level differences dominated the relationship between diversity and population size, I fit phylogenetic mixed-effects models to phyla-level subsets of the data for clades with sufficient sample sizes (see Methods: *Phylogenetic Comparative Methods*). This analysis shows a significant positive relationship between diversity and population size in arthropods, and positive weak relationships in molluscs and chordates (Supplementary Figure A15). Each of the 90% credible intervals for slope overlap, indicating the relationship between *π* and *N*_*c*_ is similar across these clades.

Additionally, I have explored the rate of trait change through time using node-height tests (Freckleton and Harvey 2006). Node-height tests regress the absolute values of the standardized contrasts between lineages against the branching time (since present) of these lineages. Under Brownian Motion (BM), standardized contrasts are estimates of the rate of character evolution (Felsenstein 1985); if a trait evolves under constant rate BM, this relationship should be flat. For both diversity and population size, node-height tests indicate a significant increase in the rate of evolution towards the present (robust regression p-values 0.023 and 0.00018 respectively; Figure 3C). Considering the constituents of the population size estimate, range and body mass, separately, the rate of evolution of range but not body mass shows a significant increase (p-value 1.03 × 10^−7^) towards the present.

Interestingly, the diversity node-height test reveals two rate shifts at deeper splits (Figure 3C, top left) around 570 Mya. These nodes represent the branches between tunicates and vertebrates in chordates, and cephalopods and pleistomollusca (bivalves and gastropods) in molluscs. While the cephalopod-pleistomollusca split outlier may be an artifact of having a single cephalopod (*Sepia officinalis*) in the phylogeny, the tunicate-vertebrate split outlier is driven by the low diversity of vertebrates and the previously-documented exceptionally high diversity of tunicates (sea squirts; Nydam and Harrison 2010; Small et al. 2007). This deep node representing a rate shift in diversity could reflect a change in either effective population size or mutation rate, and there is some evidence of both in this genus *Ciona* (Small et al. 2007; Tsagkogeorga et al. 2012). Neither of these deep rate shifts in diversity is mirrored in the population size node-height test (Figure 3C, top right). Rather, it appears a trait impacting diversity but not census size (e.g. mutation rate or offspring distributions) has experienced a shift on the lineage separating tunicates and vertebrates. At nearly 600 Mya, these deep nodes illustrate that expected coalescent times can share phylogenetic history, due to phylogenetic inertia in some combination of population size, reproductive system, and mutation rates.

Finally, an important caveat is the increase in rate towards the tips could be caused by measurement noise, or possibly uncertainty or bias in the divergence time estimates deep in the tree. Inspecting the lineage pairs that lead to this increase in rate towards the tips indicates these represent plausible rate shifts, e.g. between cosmopolitan and endemic sister species like *Drosophila simulans* and *Drosophila sechellia*; however, ruling out measurement noise entirely as an explanation would involve considering the uncertainty of diversity and population size estimates.

### Assessing the Impact of Linked Selection on Diversity Across Taxa

The above analyses reemphasize the drastic shortfall of diversity levels as compared to census sizes. Linked selection has been proposed as the mechanism that acts to reduce diversity levels from what we would expect given census sizes (Corbett-Detig et al. 2015; Gillespie 2000; Maynard Smith and Haigh 1974). Here, I test this hypothesis by estimating the scale of diversity reductions expected under background selection and recurrent hitchhiking, and compare these to the observed relationship between *π* and *N*_*c*_.

I quantify the effect of linked selection on diversity as the ratio of observed diversity (*π*) to the estimated diversity in the absence of linked selection (*π*_0_), *R* = ^*π*^*/π*_0_. Here, *π*_0_ would reflect only demographic history and non-heritable variation in reproductive success. There are two difficulties in evaluating whether linked selection could resolve Lewontin’s Paradox. The first difficulty is that *π*_0_ is unobserved. Previous work has estimated *π*_0_ using methods that exploit the spatial heterogeneity in recombination and functional density across the genome to fit linked selection models that incorporate both hitchhiking and background selection (Corbett-Detig et al. 2015; Elyashiv et al. 2016). The second difficulty is understanding of how *R* varies across taxa, since we lack estimates of critical model parameters for most species. Still, I can address a key question: if diversity levels were determined by census sizes (*π*_0_ = 4*N*_*c*_*µ*), are the combined effects of background selection and recurrent hitchhiking sufficient to reduce diversity to observed levels? Furthermore, does the relationship between census size and predicted diversity under linked selection across species, *π*_*BGS*+*HH*_ = *Rπ*_0_, match the observed relationship in Figure 2?

Since we lack estimates of key linked selection parameters across species, I parameterize the hitchhiking and BGS models using estimates from *Drosophila melanogaster*, a species known to be strongly affected by linked selection (Sella et al. 2009). Under a generalized model of hitchhiking and background selection (Coop and Ralph 2012; Elyashiv et al. 2016) and assuming *N*_*e*_ = *N*_*c*_, expected diversity is

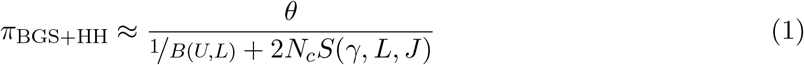

where *θ* = 4*N*_*c*_*µ, B*(*U, L*) is the effect of background selection, and *S*(*γ, L, J*) is the rate of coalescence caused by sweeps (c.f. Elyashiv et al. 2016, equation 1, Coop and Ralph 2012 equation 20). Under background selection models with recombination, the reduction is *B*(*U, L*) = exp(^−*U*^/_*L*_) where *U* is the per diploid genome per generation deleterious mutation rate, and *L* is the recombination map length (Hudson and Kaplan 1995; Hudson and Kaplan 1994; Nordborg et al. 1996). This BGS model is similar to models of effective population size under polygenic fitness variation, and can account for other modes of linked selection (Robertson 1961; Santiago and Caballero 1995, 1998, see Appendix Section A2). The coalescent rate due to sweeps is *S*(*γ, L, J*) = *γ*/_*L*_*J*, where *γ* is the number of adaptive substitutions per generation, and *J* is the probability a lineage is trapped by sweeps as they occur across the genome (c.f. *J*_2,2_ in equation 15 of Coop and Ralph 2012).

Parameterizing the model this way, I then set the key parameters that determine the impact of recurrent hitchhiking and background selection (*γ, J*, and *U*) to high values estimated from *Drosophila melanogaster* by Elyashiv et al. (2016). My estimate of the adaptive substitutions per generation (*γ*_Dmel_) based Elyashiv et al. implies a rate of sweeps per basepair of *ν*_BP,Dmel_ ≈ 2.34 × 10^−11^, which is close to other estimates from *D. melanogaster* (see Supplementary Figure A14A). The rate of deleterious mutations per diploid genome, per generation is parameterized using the estimate from Elyashiv et al., *U*_Dmel_ = 1.6, which is a bit greater than previous estimates based on Bateman-Mukai approaches (Charlesworth 1987; Mukai 1988; Mukai 1985). Finally, the probability that a lineage is trapped in a sweep, *J*_Dmel_, is calculated from the estimated genome-wide average coalescent rate due to sweeps from Elyashiv et al. (see Supplementary Figured A14B and Methods: *Predicted Reductions in Diversity* for more details on parameter estimates). Using these *Drosophila* parameters, I then explore how the predicted range of diversity levels under background selection and recurrent hitchhiking varies across species with recombination map length (*L*) and census population size (*N*_*c*_).

Previous work has found that the impact of linked selection increases with *N*_*c*_ (Corbett-Detig et al. 2015; see also Supplementary Figure A13A), and it is often thought that this is driven by higher rates of adaptive substitutions in larger populations (Ohta 1992), despite equivocal evidence (Galtier 2016). However, there is another mechanism by which species with larger population sizes might experience a greater impact of linked selection: recombinational map length, *L*, is known to correlate with body mass (Burt and Bell 1987) and thus varies inversely with population size. As this is a critical parameter that determines the genome-wide impact of both hitchhiking and background selection, I examine the relationship between recombination map length (*L*) and census population size (*N*_*c*_) across taxa, using available estimates of map lengths across species (Corbett-Detig et al. 2015; Stapley et al. 2017). I find a significant non-linear relationship using phylogenetic mixed-effects models (Figure 4A; see Methods: *Phylogenetic Comparative Methods*). There is also a correlation between map length and genome size (Supplementary Figure A9) and genome size and population size (Supplementary Figure A8). These findings are consistent with both the hypothesis that non-adaptive processes increase genome size in small-*N*_*e*_ species (Lynch and Conery 2003) which in turn could increase map lengths, as well as the hypothesis that map lengths are adaptively longer to more efficiently select against deleterious alleles (Roze 2021). Overall, the negative relationship between map length and census size indicates linked selection is expected to be stronger in short map length, high-*N*_*c*_ species.

**Figure 4:**
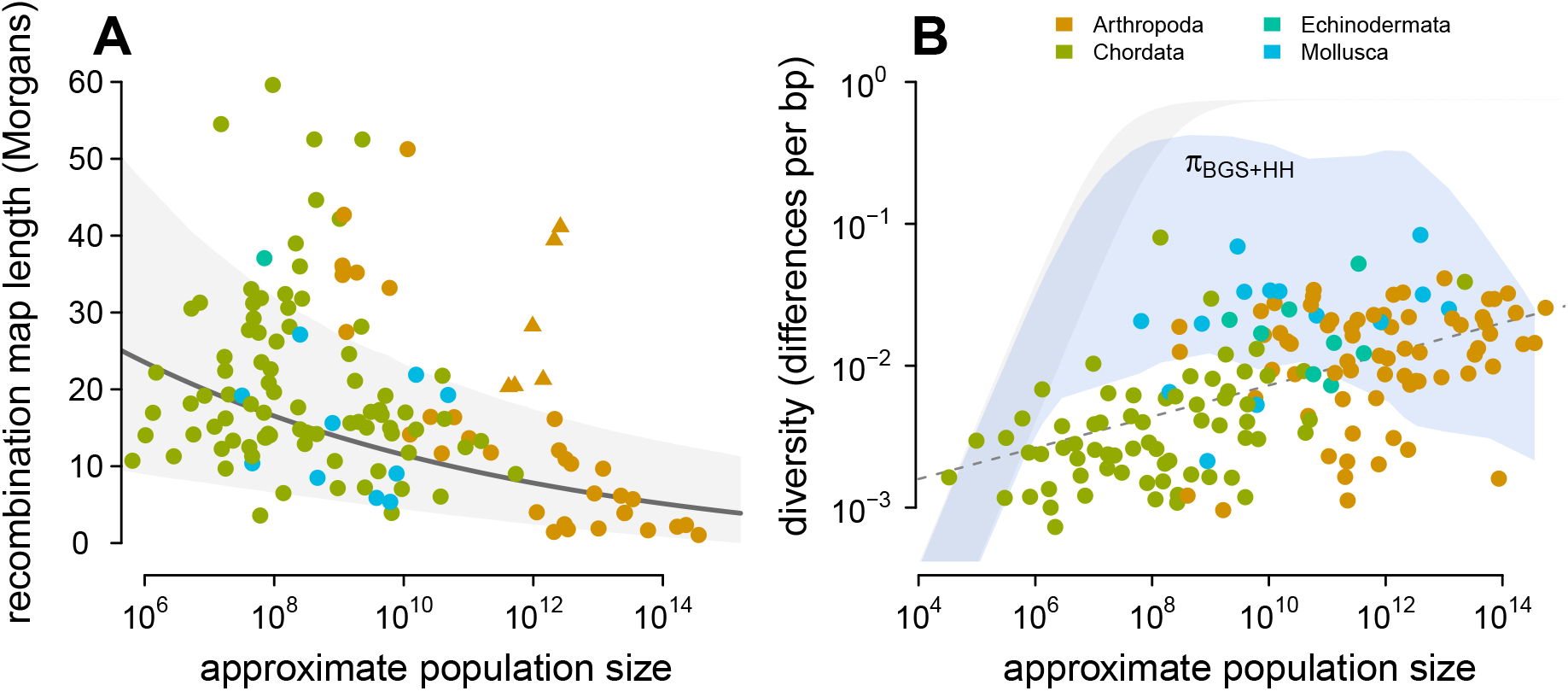
(A) The observed relationship between recombination map length (*L*) and census size (*N*_*c*_) across 136 species with complete data and known phylogeny. Triangle points indicate six social taxa excluded from the model fitting since these have adaptively higher recombination map lengths (Wilfert et al. 2007). The dark gray line is the estimated relationship under a phylogenetic mixed-effects model, and the gray interval is the 95% posterior average. (B) Points indicate the observed *π*–*N*_*c*_ relationship across taxa shown in Figure 2, and the blue ribbon is the range of predicted diversity were *N*_*e*_ = *N*_*c*_ for *µ* = 10^−8^–10^−9^, and after accounting for the expected reduction in diversity due to background selection and recurrent hitchhiking under *Drosophila melanogaster* parameters. In both plots, point color indicates phylum.

Then, I predict the expected diversity (*π*_*BGS*+*HH*_) under background selection and hitchhiking, were *N*_*e*_ = *N*_*c*_, and assuming all species had the rate of sweeps and strength of BGS as *D. melanogaster*. Since neutral mutation rates *µ* are unknown and vary across species, I calculate the range of predicted *π*_*BGS*+*HH*_ estimates for *µ* = 10^−8^–10^−9^ (using the four-alleles model, Tajima 1996), and compare this to the observed relationship between *π* and *N*_*c*_ in Figured 4B. Under these parameters, linked selection begins to appreciably depress diversity around *N*_*c*_ ≈ 10^9^, since *S* ≈ 10^−8^–10^−9^ and linked selection dominates drift when *S >* ^½^_*N*_. Overall, this reveals two problems for the hypothesis that linked selection could solve Lewontin’s Paradox. First, low to mid-*N*_*c*_ species (census sizes between 10^4^–10^10^) have sufficiently long map lengths that their diversity levels are only moderately reduced by linked selection, leading to a wide gap between predicted and observed diversity levels. For this not to be the case, the parameters that determine the strength of background selection and recurrent hitchhiking would need to be *higher* among these species than in *Drosophila melanogaster*. This would require that the rate of adaptive mutations or the deleterious mutation rate be orders of magnitude higher for species within this range than in *Drosophila*, which is incompatible with the rate of adaptive substitutions across species (Galtier 2016) and mutation rates (Lynch 2010). Furthermore, linked selection has been quantified in humans, which fall in this census size range, and has been found to be relatively weak (Boyko et al. 2008; Cai et al. 2009; Hellmann et al. 2008; Hernandez et al. 2011; McVicker et al. 2009). Second, while hitchhiking and BGS can reduce predicted diversity levels for high-*N*_*c*_ species (*N*_*c*_ *>* 10^10^) to observed levels, this would imply available estimates of *π*_0_ are underestimated by several orders of magnitude in *Drosophila* (Supplementary Figured A13B). The high reductions in *π* predicted here (compared to those of Elyashiv et al. 2016) are a result of using *N*_*c*_, rather than *N*_*e*_ = *π*_0_/4*µ* in the denominator of Equation (1), which leads to a very high rate of sweeps in the population. I do not consider selective interference, though the saturation of adaptive substitutions per Morgan would only act to limit the reduction in diversity (Weissman and Barton 2012), and thus these results are conservative. Overall, while linked selection could decouple diversity from census size for high-*N*_*c*_ species, recurrent hitchhiking and background selection seem unlikely to explain the observed patterns of diversity across species under our understanding of the range of parameter estimates.

## Discussion

Nearly fifty years after Lewontin’s description of the Paradox of Variation, how evolutionary, life history, and ecological processes interact to constrain diversity across taxa to a narrow range remains a mystery. I revisit Lewontin’s Paradox by first characterizing the relationship between genomic estimates of pairwise diversity and approximate census population size across 172 metazoan species. Previous surveys have used allozyme-based estimates, fewer taxa, or qualitative measures of population size. My estimates of census population sizes are quite approximate, since they use body size to predict density. An improved estimate might consider vagility (as Soulé 1976 did), though this is harder to do systematically across many taxa. Future work might also use other ecological information, such as total biomass, or species distribution modeling to improve census size estimates (Bar-On et al. 2018; Mora et al. 2011). Still, it seems more accurate estimates would be unlikely to change the qualitative findings here, which resemble those of early surveys (Nei and Graur 1984; Soulé 1976).

One limitation of the dataset in this study is that diversity estimates are collated from a variety of sources rather than estimated with a single bioinformatic pipeline. This leads to technical noise across diversity estimates; perhaps the relationship between *π* and *N*_*c*_ found here could be tighter with a standardized bioinformatic pipeline. In addition to this technical variation, there might be systematic bioinformatic sources of bias in diversity estimates. For example high-diversity sequences may fail to align to the reference genome and end up unaccounted for, leading to a downward bias. Alternatively, high-diversity sequences might map to the reference genome, but adjacent mis-matching SNPs might be mistaken for a short insertion or deletion. While these issues might adversely affect the estimates in high-diversity species, it is unlikely they will qualitatively change the observed *π*–*N*_*c*_ relationship.

### Macroevolution and Across-Taxa Population Genomics

Lewontin’s Paradox arises from a comparison of diversity across species, yet it has been disputed whether such comparisons require phylogenetic comparative methods. Extending previous work that has accounted for phylogeny in particular clades (Leffler et al. 2012), or using taxonomical-level averages (Romiguier et al. 2014), I show that the positive relationship between diversity and census size is significant using a mixed-effects model with a time-calibrated phylogeny. Additionally, I find a high degree of phylogenetic signal, evidence of deep shifts in the rate of evolution of genetic diversity, and that arthropods and chordates form clusters. Overall, this suggests that previous concerns about phylogenetic non-independence in comparative population genetic studies were warranted (Gillespie 1991; Whitney and Garland 2010). Notably, Lynch (2011) has argued that PCMs for pairwise diversity are unnecessary, since mutation rate evolution is fast and thus free of phylogenetic inertia, sampling variance should exceed the variance due to phylogenetic shared history, and coalescent times are much less than divergence times. Since my findings suggest PCMs are necessary in some cases, it is worthwhile to address these points.

First, Lynch has correctly pointed out that while coalescent times are much less than divergence times and should be free of phylogenetic shared history, the factors that determine coalescent times (e.g. mutation rates and effective population size) may not be (2011). In other words, coalescent times are free from phylogenetic shared history *were we to condition* on these causal factors that could be affected by shared phylogenetic history. My estimates of phylogenetic signal in diversity, by contrast, are not conditioned on these factors. Importantly, even “correcting for” phylogeny implicitly favors certain causal interpretations over others (Uyeda et al. 2018; Westoby et al. 1995). Future work could try to untangle what causal factors determine coalescent times across species, as well as how these factors evolve across macroevolutionary timescales. Second, it is a misconception that a fast rate of trait evolution necessarily reduces phylogenetic signal (Revell et al. 2008), and that if either or both variables in a regression are free of phylogenetic signal, PCMs are unnecessary (Revell 2010; Uyeda et al. 2018). The evidence of high phylogenetic signal found in this study suggests PCMs are needed, in part to avoid spurious results from phylogenetic pseudoreplication. Finally, beyond just accounting for phylogenetic non-independence, macroevolution and phylogenetic comparative methods are a promising way to approach across-species population genomic questions. For example, one could imagine that diversification processes could contribute to Lewontin’s Paradox. If large-*N*_*c*_ species were to have a rate of speciation that is greater than the rate at which mutation and drift reach equilibrium (which is indeed slower for large *N*_*c*_ species), this could act to decouple diversity from census population size. That is to say, even if the rate of random demographic bottlenecks were constant across taxa, lineage-specific diversification processes could lead certain clades to be systematically further from demographic equilibrium, and thus have lower diversity than expected for their census population size.

### Spatial and Demographic Processes

One limitation of this study is the inability to quantify the impact of spatial population genetic processes on the relationship between diversity and census population sizes across taxa. The genomic diversity estimates collated in this study unfortunately lack details about the sampling process and spatial data, which can have a profound impact on population genomic summary statistics (Battey et al. 2020). These issues could systematically bias species-wide diversity estimates; for example, if diversity estimates from a cosmopolitan species were primarily from a single subpopulation, diversity would be an underestimate relative to the entire population. However, biased spatial sampling alone seems incapable of explaining the *π*-*N*_*c*_ divergence in high-*N*_*c*_ taxa. In the extreme scenario in which only one subpopulation was sampled, *F*_ST_ would need to be close to one for population subdivision alone to sufficiently reduce the total population heterozygosity to explain the orders-of-magnitude shortfall between predicted and observed diversity levels. This is because the equation for *F*_*ST*_ can be rearranged such that *H*_*S*_ = (1 − *F*_*ST*_)*H*_*T*_, where *H*_*S*_ and *H*_*S*_ are the subpopulation and total population heterozygosities; if *H*_*T*_ = 4*N*_*c*_*µ*, then only *F*_*ST*_ ≈ 1 can reduce *H*_*S*_ several orders of magnitude. Yet, across-taxa surveys indicate that *F*_ST_ is almost never this high within species (Roux et al. 2016). Still, future work could quantify the extent to which spatial processes contribute to Lewontin’s Paradox. For example, high-*N*_*c*_ taxa usually experience range expansions, likely with repeated founder effects and local extinction/recolonization dynamics that doubtlessly depress diversity. In particular, with the appropriate data, one could estimate the empirical relationship between dispersal distance, range size, and coalescent effective population size across taxa.

In this study, I have focused entirely on assessing the role of linked selection, rather than demography, in reducing diversity across taxa. In contrast to demographic models, models of linked selection have comparatively fewer parameters and more readily permit rough estimates of diversity reductions across taxa. Still, a full resolution of Lewontin’s Paradox would require understanding how the demographic processes across taxa with incredibly heterogeneous ecologies and life histories transform *N*_*c*_ into *N*_*e*_. With population genomic data becoming available for more species, this could involve systematically inferring the demographic histories of tens of species and looking for correlations in the frequency and size of bottlenecks with *N*_*c*_ across species.

### How could selection still explain Lewontin’s Paradox?

In this study, my goal was not to accurately estimate the levels in diversity across species, but rather to give linked selection the best possible chance to solve Lewontin’s Paradox. Still, I find that even after parameterizing hitchhiking and background selection with strong selection parameter estimates from *Drosophila melanogaster*, the predicted patterns of diversity under linked selection poorly fit observed patterns of diversity across species. This result extends the analysis by Coop (2016) showing that levels of *π*_0_ estimated by Corbett-Detig et al. (2015) are not decoupled from genome-wide average *π*, as would occur if linked selection were to explain Lewontin’s Paradox. Here, my analysis goes a step further and suggests that models of recurrent hitchhiking and background selection are not capable of explaining the observed relationship between *π* and census size, in part because mid-*N*_*c*_ species have sufficiently long recombination map lengths to diminish the effects of even strong selection. This finding supports the idea the levels of diversity across species are primarily determined by past demographic fluctuations. Overall, while this suggests these two common modes of linked selection seem unlikely to explain across-taxa patterns of diversity, there are three major potential limitations of my approach that need further evaluation.

First, I approximate the reduction in diversity using homogeneous background selection and recurrent hitchhiking models (Coop and Ralph 2012; Hudson and Kaplan 1995; Kaplan et al. 1989), when in reality, there is genome-wide heterogeneity in functional density, recombination rates, and the adaptive substitutions across species. Each of these factors mediate how strongly linked selection impacts diversity across the genome. Despite these model simplifications, the predicted reduction in diversity in *Drosophila melanogaster* is 85% (when using *N*_*e*_, not *N*_*c*_), which is reasonably close to the estimated 77% from the more realistic model of Elyashiv et al. that accounts for the actual position of substitutions, annotation features, and recombination rate heterogeneity (though it should be noted that these both use the same parameter estimates). Furthermore, even though my model fails to capture the heterogeneity of functionality density and recombination rate in real genomes, it is still extraordinary conservative, likely overestimating the effects of linked selection to see if it could be capable of decoupling diversity from census size and explain Lewontin’s Paradox. This is in part because the strong selection parameter estimates from *Drosophila melanogaster* used, but also because I assume that the effective population size is equal to the census size. Even then, this decoupling only occurs in very high–census-size species, and implies that the diversity in the absence of linked selection, *π*_0_, is currently underestimated by several orders of magnitude. Moreover, the study of Corbett-Detig et al. (2015) did consider recombination rate and functional density heterogeneity in estimating the reduction due to linked selection across species, yet their predicted reductions are orders of magnitude weaker than those considered here by assuming that *N*_*e*_ = *N*_*c*_ (Supplementary Figure A13B). Overall, even with more realistic models of linked selection, current models of linked selection seem fundamentally unable to fit the diversity–census-size relationship.

Second, my model here only considers hard sweeps, and ignores the contribution of soft sweeps (e.g. from standing variation or recurrent mutations; Hermisson and Pennings 2005; Pennings and Hermisson 2006), partial sweeps (e.g those that do not reach fixation), and the interaction of sweeps and spatial processes. While future work exploring these alternative types of sweeps is needed, the predicted reductions in diversity found here under the simplified sweep model are likely relatively robust to these other modes of sweeps for a few reasons. First, the shape of the diversity– recombination curve is equivalent under models of partial sweeps and hard sweeps, though these imply different rates of sweeps (Coop and Ralph 2012). Second, in the limit where most fitness variation is due to weak soft sweeps from standing variation scattered across the genome (i.e. due to polygenic fitness variation), levels of diversity are well approximated by quantitative genetic linked selection models (Robertson 1961; Santiago and Caballero 1995, 1998). The reduction in diversity under these models is nearly identical to that under background selection models, in part because deleterious alleles at mutation-selection balance constitute a considerable component of fitness variation (see Appendix Section A2; Charlesworth 2015; Charlesworth and Hughes 2000). Third, the parameters from Elyashiv et al. (2016) are robust to many types of sweeps that result in substitution (e.g. see p. 19 of their Supplementary Online Materials). Finally, I also disregarded the interaction of sweeps and spatial processes. For populations spread over wide ranges, limited dispersal slows the spread of sweeps, allowing for new beneficial alleles to arise, spread, and compete against other segregating beneficial variants (Ralph and Coop 2010; Ralph and Coop 2015). Through limited dispersal should act to “soften sweeps” and not impact my findings for the reasons described above, future work could investigate how these processes impact diversity in ways not captured by hard sweep models.

Third, other selective processes, such as fluctuating selection or hard selective events, could reduce diversity in ways not captured by the background selection and hitchhiking model. Since frequency-independent fluctuating selection generally reduces diversity under most conditions (Novak and Barton 2017), this could lead seasonality and other sources of temporal heterogeneity to reduce diversity in large-*N*_*c*_ species with short generation times more than longer-lived species with smaller population sizes. Future work could consider the impact of fluctuating selection on diversity under simple models (Barton 2000) if estimates of key parameters governing the rate of such fluctuations were known across taxa. Additionally, another mode of selection that could severely reduce diversity across taxa, yet remains unaccounted for in this study, is periodic hard selective events. These selective events could occur regularly in a species’ history yet be indistinguishable from demographic bottlenecks with just population genomic data.

### Measures of Effective Population Size, Timescales, and Lewontin’s Paradox

Lewontin’s Paradox describes the extent to which the effective population sizes implied by diversity, *Ñ*_*e*_, diverge from census population sizes. However, there are a variety other effective population size estimates calculable from different data and summary statistics (Caballero 1994; Caballero 2020; Galtier and Rousselle 2020; Wang et al. 2016). These include estimators based on the site frequency spectrum, observed decay in linkage disequilibrium, or temporal estimators that use the variance in allele frequency change. These alternate estimators capture summaries of the effective population size on shorter timescales than coalescent-based estimators (Wang 2005), and thus could be used to tease apart processes that impact the *N*_*e*_-*N*_*c*_ relationship in the more recent past.

Temporal *N*_*e*_ estimators already play an important role in understanding another summary of the *N*_*e*_-*N*_*c*_ relationship: the ratio *N*_*e*_/*N*_*c*_, which is an important quantity in conservation genetics (Frankham 1995; Mace and Lande 1991) and in understanding evolution in highly fecund marine species. Surveys of the short-term *N*_*e*_/*N*_*c*_ relationship across taxa indicate mean *N*_*e*_/*N*_*c*_ is on order of ≈ 0.1 (Frankham 1995; Palstra and Fraser 2012; Palstra and Ruzzante 2008), though the uncertainty in these estimates is high, and some species with sweepstakes reproduction systems like Pacific Oyster (*Crassostrea gigas*) can have *N*_*e*_/*N*_*c*_ ≈ 10^−6^. Estimates of the *N*_*e*_/*N*_*c*_ ratio are an important, yet under appreciated piece of solving Lewontin’s Paradox. For example, if *N*_*e*_ is estimated from the allele frequency change across a single generation (i.e. Waples 1989), *N*_*e*_/*N*_*c*_ constrains the variance in reproductive success (Nunney 1993, 1996; Wright 1938). This implies that apart from species with sweepstakes reproductive systems, the variance in reproductives success each generation (whether heritable or non-heritable) is likely insufficient to significantly contribute to constraining *N*_*e*_ for most taxa. Still, further work is needed to characterize (1) how *N*_*e*_/*N*_*c*_ varies with *N*_*c*_ across taxa (though see Palstra and Fraser 2012, Figure 2), and (2) the variance of *N*_*e*_/*N*_*c*_ over longer time spans (i.e. how periodic sweepstakes reproductive events act to constrain *N*_*e*_). Overall, characterizing how *N*_*e*_/*N*_*c*_ varies across taxa and correlates with ecology and life history traits could provide clues into the mechanisms that leads propagule size and survivorship curves to be predictive of diversity levels across taxa (Barry et al. 2020; Hallatschek 2018; Romiguier et al. 2014).

Finally, short-term temporal *N*_*e*_ estimators may play an important role in resolving Lewontin’s Paradox. These estimators, along with short-term estimates of the impact of linked selection (Buffalo and Coop 2019, 2020), can inform us how much diversity is depressed across shorter timescales, free from the rare strong selective events or severe bottlenecks that impact pairwise diversity. It could be that in any one generation, selection contributes more to the variance of allele frequency changes than drift, yet across-taxa patterns in diversity are better explained processes acting sporadically on longer timescales, such as colonization, founder effects, and bottlenecks. Thus, the pairwise diversity may not give us the best picture of the generation to generation evolutionary processes acting in a population to change allele frequencies. Furthermore, certain observed adaptations are inexplicable given implied long-term coalescent effective population sizes, and are only possible if short-term effective population sizes are orders of magnitude larger (Barton 2010; Karasov et al. 2010).

## Conclusions

In *Building a Science of Population Biology* (2004), Lewontin laments the difficulty of uniting population genetics and population ecology into a cohesive discipline of population biology. Lewontin’s Paradox of Variation remains a critical unsolved problem at the nexus of these two different disciplines: across species, we fail to understand the processes that connect a central parameter of population ecology, census size, to a central parameter of population genetics, effective population size. Given that selection seems to fall short in explaining Lewontin’s Paradox, a full resolution will require a mechanistic understanding the ecological, life history, and macroevolutionary processes that connect *N*_*c*_ to *N*_*e*_ across taxa. While I have focused exclusively on metazoan taxa since their population densities are more readily approximated from body mass, a full resolution must also include plant species (with the added difficulties of variation in selfing rates, different dispersal strategies, pollination, etc.).

Looking at Lewontin’s Paradox through an macroecological and macroevolutionary lens begets interesting questions outside of the traditional realm of population genetics. Here, I have found that diversity and *N*_*c*_ have a surprisingly consistent relationship without many outliers, despite the wildly disparate ecologies, life histories, and evolutionary histories of the taxa included. Furthermore, taxa with very large census sizes have surprisingly low diversity. Is this explained by macroevolutionary processes, such as different rates of speciation for large-*N*_*c*_ taxa? Or, are the levels of diversity we observe today an artifact of our timing relative to the last glacial maximum, or the last major extinction? Did large-*N*_*c*_ prehistoric animal populations living in other geological eras have higher levels of diversity than our present taxa? Or, does ecological competition occur on shorter timescales such that strong population size contractions transpire and depress diversity, even if a species is undisturbed by climatic shifts or mass extinctions? Overall, patterns of diversity across taxa are determined by many overlaid evolutionary and ecological processes occurring on vastly different timescales. Lewontin’s Paradox of Variation may persist unresolved for some time because the explanation requires synthesis and model building at the intersection of all these disciplines.

## Methods

### Diversity and Map Length Data

The data used in this study are collated from a variety of previously published surveys. Of the 172 taxa with diversity estimates, 14 are from Corbett-Detig et al. (2015), 96 are from Leffler et al. (2012), and 62 are from Romiguier et al. (2014). The Corbett-Detig et al. data is estimated from four-fold degenerate sites, the Romiguier et al. data is synonymous sites, and the Leffler et al. data is estimated predominantly from silent, intronic, and non-coding sites. All types of diversity estimates from Leffler et al. (2012) were included to maximize the taxa in the study, since the variability of diversity across functional categories is much less than the diversity across taxa. Multiple diversity estimates per taxa were averaged. The total recombination map length data were from both Stapley et al. (2017; 127 taxa), and Corbett-Detig et al. (2015; 9 taxa). Both studies used sex-averaged recombination maps estimated with cross-based approaches; in some cases errors in the original data were found, documented, and corrected. These studies also included genome size estimates used to create Supplementary Figures A8 and A9.

### Macroecological Estimates of Population Size

A rough approximation for total population size (census size) is *N*_*c*_ = *DR*, where *D* is the population density in individuals per km^2^ and *R* is the range size in km^2^. Since population density estimates are not available for many taxa included in this study, I used the macroecological abundance-body size relationship to predict population density from body size. Since body length measurements are more readily available than body mass, I collated body length data from various sources (see https://github.com/vsbuffalo/paradox_variation/); body lengths were averaged across sexes for sexually dimorphic species, and if only a range of lengths was available, the midpoint was used.

Then, I re-estimated the relationship between body mass and population density using the data in the appendix table of Damuth (1987), which includes 696 taxa with body mass and population density measurements across mammals, fish, reptiles, amphibians, aquatic invertebrates, and terrestrial arthropods. Though the abundance-body size relationship can be noisy at small spatial or phylogenetic scales (Chapter 5, Gaston and Blackburn 2008), across deeply diverged taxa such as those included in this study and Damuth (1987), the relationship is linear and homoscedastic (see Supplementary Figure A4). Using Stan (Stan Development Team 2020), I jointly estimated the relationship between body mass from body length using the Romiguier et al. (2014) taxa, and used this relationship to predict body mass for the taxa in this study. These body masses were then used to predict population density simultaneously, using the Damuth (1981) relationship. The code of this routine (`pred popsize missing centered.stan`) is available in the GitHub repository (https://github.com/vsbuffalo/paradox_variation/).

To estimate range, I first downloaded occurrence records from Global Biodiversity Information Facility (*GBIF Occurrence Download* 2020) using the `rgbif` R package (Chamberlain and Boettiger 2017; Chamberlain et al. 2014). Using the occurrence locations, I inferred whether a species was marine or terrestrial, based on whether the majority of their recorded occurrences overlapped a continent using `rnaturalearth` and the `sf` packages (Pebesma 2018; South 2017). For each taxon, I estimated its range by finding the minimum *α*-shape containing these occurrences. The *α* parameters were set more permissive for marine species since occurrence data for marine taxa were sparser. Then, I intersected the inferred ranges for terrestrial taxa with continental polygons, so their ranges did not overrun landmasses (and likewise with marine taxa and oceans). I inspected diagnostic plots for each taxa for quality control (all of these plots are available in `paradox variation` GitHub repository), and in some cases, I manually adjusted the *α* parameter or manually corrected the range based on known range maps (these changes are documented in the code `data/species ranges.r` and `data/species range fixes.r`). The range of *C. elegans* was conservatively approximated as the area of the Western US and Western Europe based on the map in Frézal and Félix (2015). *Drosophila* species ranges are from the Drosophila Speciation Patterns website, (Yukilevich 2017; Yukilevich 2012). To further validate these range estimates, I have compared these to the qualitative range descriptions Leffler et al. (2012) (Supplementary Figure A11) and compared my *α*-shape method to a subset of taxa with range estimates from IUCN Red List (Chamberlain 2020; IUCN 2020; Supplementary Figure A10). Each census population size is then estimated as the product of range and density.

### Phylogenetic Comparative Methods

Of the full dataset of 172 taxa with diversity and population size estimates, a synthetic calibrated phylogeny was created for 166 species that appear in phylogenies in DateLife project (O’Meara et al. 2020; Sanchez-Reyes and O’Meara 2019). This calibrated synthetic phylogeny was then subset for the analyses based on what species had complete trait data. The diversity-population size relationship assessed by a linear phylogenetic mixed-effects model implemented in Stan (Stan Development Team 2020), according to the methods described in (Villemereuil and Nakagawa 2014, see `stan/phylo_mm_regression.stan` in the GitHub repository). This same Stan model was used to estimate the same relationship between arthropod, chordate, and mollusc subsets of the data, though a reduced model was used for the chordate subset due to identifiability issues leading to poor MCMC convergence (Supplementary Figure A15).

The relationship between recombination map length and the logarithm of population size is non-linear and heteroscedastic, and was fit using a lognormal phylogenetic mixed-effects model on the 130 species with complete data. Since social insects have longer recombination map lengths (Wilfert et al. 2007), social taxa were excluded when fitting this model. All `Rhat` (Vehtari et al. 2019) values were below 1.01 and the effective number of samples was over 1,000, consistent with good mixing; details about the model are available in the GitHub repository (`phylo mm lognormal.stan`). Continuous trait maps (Figure 3A and Supplementary Information Figure A17) were created using `phytools` (Revell 2012). Node-height tests were implemented based on the methods in `Geiger` (Harmon et al. 2008; Pennell et al. 2014), and use robust regression to fit a linear relationship between phylogenetic independent contrasts and branching times.

### Predicted Reductions in Diversity

The predicted reductions in diversity due to linked selection are approximated using selection and deleterious mutation parameters from *Drosophila melanogaster*, and the recombination map length estimates from Stapley et al. (2017) and Corbett-Detig et al. (2015). The mathematical details of the simplified sweep model are explained in the Appendix Section A1. I use estimates of the number of substitutions, *m*, in genic regions between *D. melanogaster* and *D. simulans* from Hu et al. (2013). Following Elyashiv et al. (2016), only substitutions in UTRs and exons are included, since they found no evidence of sweeps in introns. Then, I average over annotation classes to estimate the mean proportion of substitutions that are beneficial, *α*_Dmel_ = 0.42, which are consistent with the estimates of Elyashiv et al. and estimates from MacDonald–Kreitman test approaches (see Eyre-Walker 2006, Table 1). Then, I use divergence time estimates between *D. melanogaster* and *D. simulans* of 4.2 × 10^6^ and estimate of ten generations per year (Obbard et al. 2012), calculating there are *γ*_Dmel_ = ^*αm*^/_2*T*_ = 2.26 × 10^−3^ substitutions per generation. Given the length of the *Drosophila* autosomes, *G*, this implies that the rate of beneficial substitutions per basepair, per generation is *ν*_*BP,Dmel*_ = *γ*_Dmel_/_*G*_ = 2.34 × 10^−11^. Finally, I estimate *J*_Dmel_ from the estimate of genome-wide average rate of sweeps from Elyashiv et al. (Supplementary Table S6) and assuming *Drosophila N*_*e*_ = 10^6^. These *Drosophila melanogaster* hitchhiking parameter estimates are close to other previously-published estimates (Supplementary Figure A14). Finally, I use *U*_Dmel_ = 1.6, from Elyashiv et al. (2016). With these parameter estimates from *D. melanogaster*, the recombination map lengths across species, and Equation (1), I estimate *π*_BGS+HH_ (assuming *N*_*c*_ = *N*_*c*_) across all species. This leads to a range of predicted diversity ranges across species corresponding to *µ* = 10^−8^–10^−9^; to visualize these, I take a convex hull of all diversity ranges and smooth this with R’s `smooth.spline` function.

## Supporting information

A PDF of the differences between this version and the last version.

## Acknowledgments

I would like to thank Andy Kern and Peter Ralph for helpful discussions and supporting me during this work, and Graham Coop for inspiration and helpful feedback during socially distanced nature walks at Yolo Basin. I thank Jessica Stapley for kindly providing the recombination map length data, and Yaniv Brandvain, Amy Collins, Doc Edge, Tyler Kent, Chuck Langley, Matt Osmond, Sally Otto, Jeff Ross-Ibarra, Aaron Stern, Anastasia Teterina, Michael Turelli, Margot Wood, and my Kern-Ralph labmates for helpful discussions. Sarah Friedman, Katherine Corn, and Josef Uyeda provided very useful advice about phylogenetic comparative methods; yet I take full responsibility for any shortcomings of my analysis. Finally, I am indebted to Guy Sella, Matt Pennell, and two other anonymous reviewers for helpful feedback. I would like to also thank UO librarian Dean Walton for helping me track down some rather difficult to find older papers. This work was supported by an NIH Grant (1R01GM117241) awarded to Andrew Kern.

## Appendix

### A1 Simplified Sweep Effects Model

I use a simplified model of the effects of recurrent hitchhiking and background selection (BGS) occurring uniformly along a genome. Expected diversity is given by

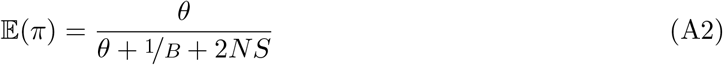

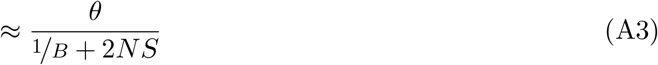

(cf. equation 1 Elyashiv et al. 2016, and equation 20 of Coop and Ralph 2012). The BGS component is given by Hudson and Kaplan (1995),

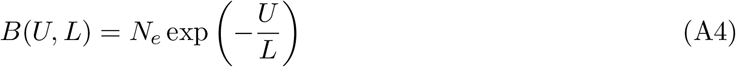

and the hitchhiking component is

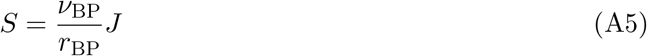

(cf. Coop and Ralph 2012 equation 20) where *J* is the probability that two lineages coalesce down to one, given sweeps occur uniformly along the genome. Under this homogeneous sweep model, *J* is

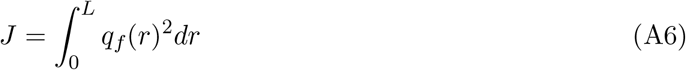

where *q*_*f*_ (*r*) is the approximate probability that a lineage is trapped by a sweep to frequency *f* when it is *r* recombination fraction away from this sweep (cf. Coop and Ralph 2012 equation 15). Since I use *Drosophila melanogaster* parameter estimates from Elyashiv et al. (2016), I now reconcile their model’s *S* term with the simple model above. They estimate *S* in *Drosophila melanogaster* using a composite likelihood model that considers hitchhiking and background selection simultaneously, using substitutions and stratifying by annotation. For a neutral position at site *x*, the coalescent rate due to sweeps is given by Elyashiv et al.’s equation 3,

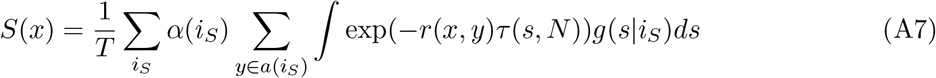

where *T* is the number of generations that substitutions accrue, *i*_*S*_ = 1, …, *I*_*S*_ is the annotation class (e.g. exons, introns, UTRs), *α*(*i*_*S*_) is the fraction of substitutions in annotation class *i*_*S*_ that are beneficial, *a*(*i*_*S*_) is the set of all substitutions in annotation class *i*_*S*_, *τ* (*s, N*) is the fixation time of a site with additive effect *s*, and *g*(*s*|*i*_*S*_) is the distribution of selection coefficients for annotation class *i*_*S*_.

Note, that we can recover the model of Coop and Ralph (2012) from this expression. Suppose there is only one annotation class, and *α* fraction of substitutions are beneficial, and one selection coefficient 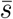, (i.e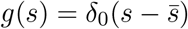), then

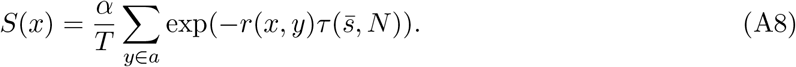

Let the number of substitutions be *m* := |*a*|, and imagine their positions are uniformly distributed on a segment of length *G* basepairs with the focal site is the middle at position *x* = 0. Then, each substitution *y* is a random distance *l*_*y*_ ∼ *U* (−*G*/_2_, ^*G*^/_2_) away from the focal site. Assuming the recombination rate is a constant *r*_BP_ per basepair, and approximating the sum with an integral, we have,

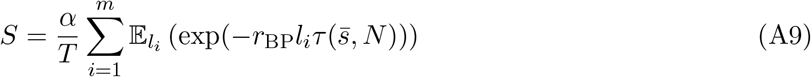

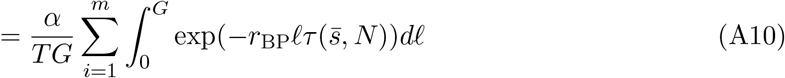

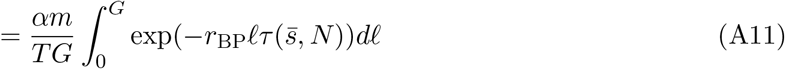

Using *u*-substitution with *r* = *ℓr*_BP_ this simplifies to

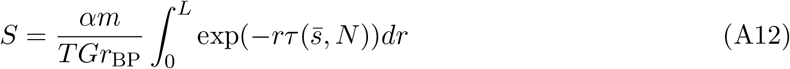

where *L* = *Gr*_BP_.

To simplify this notation, note that the rate of adaptive substitutions per basepair per generation is *ν*_BP_ = *αm*/_*GT*_, so

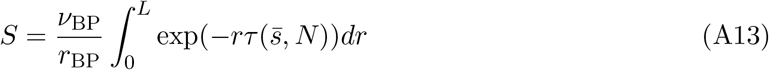

This is analogous to the second term of Coop and Ralph (2012) equation 17, with *k* = *i* = 2 and *x* = 1 (e.g. conditioning on a sweep to fixation). Note that there appears to be a factor of two error in Elyashiv et al. (2016) compared to Coop and Ralph (2012); here I include the factor of two. Then,

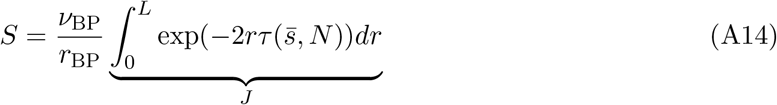

where the integral is equal to *J* (c.f. *J*_2,2_ of equation 15 in Coop and Ralph 2012) since a simple model of *q*_*f*_ (*r*) = *f* exp(−2*rτ* (*s, N*)) and if we condition on fixation, *f* = 1. This expression is useful to generalize across species, since we know *N* and *L*. Additionally, we have estimates of *α* and ^*m*^/_*T*_ in *Drosophila* and other species. In Elyashiv et al, they consider the number of substitutions per generation in genic regions only; it should be noted that the number of coding basepairs varies little across species. For convenience, I define *γ* = *αm*/_*T*_ as the number of adaptive substitutions per generation per entire genome, such that *S*(*γ, L, J*) = *γ*/_*L*_ *J* used in the main text. Using the estimates of *m* ≈ 4.5 × 10^5^, *α* ≈ 0.42, and *T* ≈ 8.4 × 10^7^ from the Supplementary Material of Elyashiv et al., I arrive at *γ* ≈ 0.00226 adaptive substitutions per generation, per genome. For a ≈ 100 megabase genome, this translates to a *ν*_BP_ ≈ 2.34 × 10^−11^, which is close to previous estimates (Supplementary Figure A14). For *J*, I use an empirical estimate calculated from the genome-wide average of the rate of coalescent events due to sweeps, from Supplementary Table S6 of Elyashiv et al. (*r*_*s*_ = 2*NS* ≈ 0.92). This implies *J* ≈ 4.46 × 10^−4^. Alternatively, I have tried using the estimated distribution of selection coefficients from Elyashiv et al., but this led to a weaker estimate of *J*, since the adaptive substitutions considered tend to cluster around genic regions. Note that these *Drosophila* sweep parameters I have used are close to previous estimates (Supplementary Figures A14 A and B).

### A2 Background Selection and Polygenic Fitness Models

Throughout the main text, I use recurrent hitchhiking and background selection models to estimate the reduction in diversity due to linked selection. Another class of linked selection models, which I refer to as quantitative genetic linked selection models (QGLS; Robertson 1961; Santiago and Caballero 1995, 1998), can also depress genome-wide diversity. Furthermore, these models may depress diversity at neutral sites unlinked to the regions containing fitness variation. While I did not explicitly incorporate these models into my estimates of the diversity reductions, their effect is implicit in background selection models because they are analytically nearly identical. Here, I briefly sketch out the connection between BGS and QGLS models.

Under the Santiago and Caballero (1998) model, the effective population size is 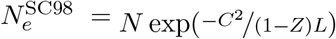, where *C*^2^ is the standardized heritable fitness variation, 1 − *Z* is the decay of genetic variance through time, and *L* is the recombination map length. This model can accommodate a variety of modes of selection such as selection on an infinitesimal trait (Santiago and Caballero 1995, p. 1016), and the flux of either weakly advantageous or deleterious alleles (Santiago and Caballero 1998, p. 2109). If the source of fitness variation is entirely the input of new deleterious mutations with heterozygous effect *sh* at rate *U* per diploid genome per generation, then under mutation-selection balance, the equilibrium relative variance in reproductive success *C*^2^ = *Ush* (Crow and Kimura 1970; Caballero 2020, p. 167), and *Z* = 1 − *sh* − ½*N*_*c*_ (Santiago and Caballero 1998). Thus, if ½*N*_*c*_ *<< sh <<* 1, then *C*^2^/(1−*Z* ≈ *U* and 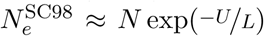, which is the BGS model used in the main text and is a result of many background selection models with similar assumptions (Hudson and Kaplan 1994 eqn. 15; Hudson and Kaplan 1995 eqn. 9; Nordborg et al. 1996 eqn. 4; Barton 1995 eqn. 22b). Intuitively, the similarity of these models reflects the fact that a substantial proportion of heritable fitness variation is caused by the continual flux of deleterious alleles across the genome under mutation-selection balance (Charlesworth 2015; Charlesworth and Hughes 2000).

## Supplementary Information

### S1 Population Size Validation

I validated the approximate census sizes I have estimated here using a few approaches. First, to check broad consistency, I compared the implied biomass from my estimates of body mass and census size per species with estimates of the total carbon biomass on earth by phylum (Bar-On et al. 2018). For species *i* with wet body mass *m*_*i*_ and census size *N*_*i*_, the implied biomass is *m*_*i*_*N*_*i*_. For all species in a phylum *S*, this total sample biomass is *b*_*S*_ =∑_*i∈S*_ *m*_*i*_*N*_*i*_. I then compare this wet biomass to the carbon biomasses by phylum by Bar-On et al. (2018). Across animal species, the ratio of dry to wet body mass, and carbon body mass dry body mass varies little. In their study, Bar-On et al. assume wet body mass has a 70% water content, and 50% of dry body mass is carbon mass, leading to a wet body mass to carbon mass factor of 1−0.7/0.5 = 0.15. I use this factor to convert the total wet biomass to carbon biomass per phylum.

**Table S1:**
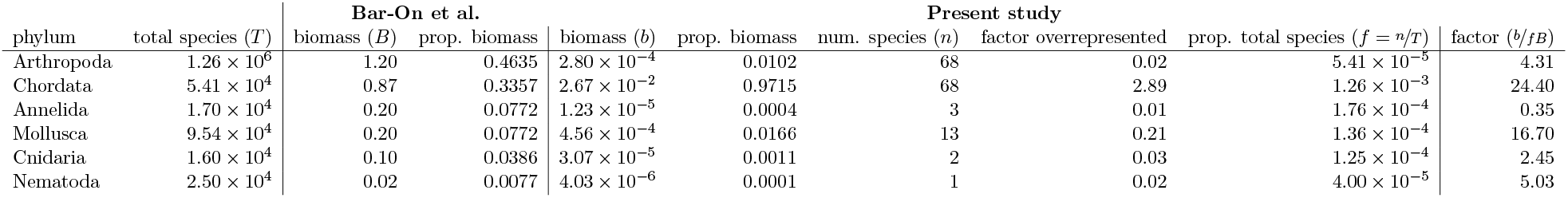
How the total carbon biomass estimates by phylum from Bar-On et al. (2018) compare to the implied biomass estimates from this study. All biomass estimates are carbon biomass, and the proportions are of total biomass with respect to the study. The proportion of biomass in this study compared to the Bar-On et al. estimates Bar-On et al. (2018) indicates chordates are overrepresented and arthropods are underrepresented in the present study; the factor that each phylum is overrepresented is given in the eighth column. Total species by phylum estimates are from Chapman et al. (2009), Nicol (1969), Reaka-Kudla et al. (1996), and Zhang (2013). The ratio column is the ratio of total biomass implied by the *N*_*c*_ estimates of each species in a phylum to the actual biomass of that phylum.

First, I compared the relative carbon biomass in this study to the relative carbon biomass on earth per phylum. This shows that this study’s sample over represents chordate biomass (by a factor of ∼3), and under represents in arthropod biomass (by a factor of 0.02) relative to the proportion of carbon biomass of these phyla on earth (see column eight of Supplementary Table S1. Second, to check whether the carbon biomass per phylum in the sample was broadly consistent with the total on earth by phylum (*B*_*S*_ for phylum *S*), I calculated the expected sample biomass if species were sampled randomly from the total species in a phylum, (*B*_*S*_ × *N*_*S*_*/T*_*S*_, where *n*_*S*_ is the total number of species in the sample in phylum *S, T*_*S*_ is the total number of species in phylum *S* on earth). The fraction of total species on earth included in the sample in this study is depicted in Supplementary Figure A2.

Next, I look at the ratio of sample biomass per phylum, *b*_*S*_ to this expected biomass per phylum (Supplementary Table S1). The consistency is quite close for this rough approach and the non-random sample of taxa included in this study. The carbon biomass estimates for chordates implied by the census size estimates are ∼24-fold higher than expected, but is well within reasonable expectations given that the chordate sample includes many larger-bodied domesticated species (and is a biased sample in other ways). Similarly, the implied arthropod carbon biomass is quite close to what one would expect. Overall, these values indicate that the census size estimates here do not lead to implied biomasses per phylum that are outside the range of plausibility.

Additionally, note that the body mass based estimates of density for *Drosophila* are similar to previously used estimates in surveys of census size and diversity. Nei and Graur (1984) suggested a maximum of 5 *Drosophila* per m^2^, including regions of the range that are not inhabitable. Across *Drosophila*, the body mass based estimates suggest 10^6.7^ − 10^7.6^ individuals per km^2^, or 4.5 − 36.3 individuals per m^2^, which are consistent with this previous estimate. Nei and Graur’s estimates of *Drosophila pseudoobscura*’s census size are four orders of magnitude smaller than mine, but their approach uses a speculated ratio of population sizes of different *Drosophila* species rather than range sizes (Nei and Graur 1984, p. 81).

As another consistency check, I looked at the rank order of mammals by biomass. Whale species have the first and third highest biomass with 11.4 and 3.9 megatons of carbon biomass (for *Balaenoptera bonaerensis* and *Eschrichtius robustus*, respectively). While this seems high, a recent study shows that across whale species, pre-whaling carbon biomass was at the tens of megatons level (Pershing et al. 2010, Table 1 and Figure 1). Given that my census size estimates represent populations at a macroecological equilibrium, they would not reflect reduced density due to whaling or other anthropogenic causes. Humans had the second largest biomass, followed by wolf species (*Canis lupus* and *C. latrans*); as with whales, the population sizes for wolf species represent pre-anthropogenic densities and are overestimates compared to current population sizes, as expected.

Finally, there are other estimates of approximate population sizes for some species that I compared my estimates to. The United Nation’s FAOSTAT database estimates the total number of horses (*Equus caballus*) on earth as ∼60 million; the estimate in this study is close to 40 million. For other domesticated species like chicken (*Gallus gallus*), estimates range from 25 million to 19.6 billion (*FAOSTAT statistics database* 2021; Robinson et al. 2014); the present study’s estimate lies in the middle at ∼175 million. Again, this is a known limitation of this method, as the range is estimated from occurrence data and does not consider species’ niches. This present study’s estimate of the number of king penguins (*Aptenodytes patagonicus*) is about 3 million; the population size was recently estimated as 2.23 million pairs (Shirihai 2008).

### S2 Sensitivity Analysis

### S3 Diversity and IUCN Red List Status

I also investigated the relationship between species’ IUCN Red List categories (an ordinal scale of how threatened a species is) and both diversity and population size, finding that species categorized as more threatened have both smaller population sizes and reduced diversity, compared to non-threatened species (Supplementary Figure A3) consistent with past work (Spielman et al. 2004). A linear model of diversity regressed on population size has lower AIC when the IUCN Red List categories are included, and the estimates of the effect of IUCN status are all negative on diversity, though not all are significant in part because some categories have three or fewer species (Supplementary Table S3).

**Table S2:**
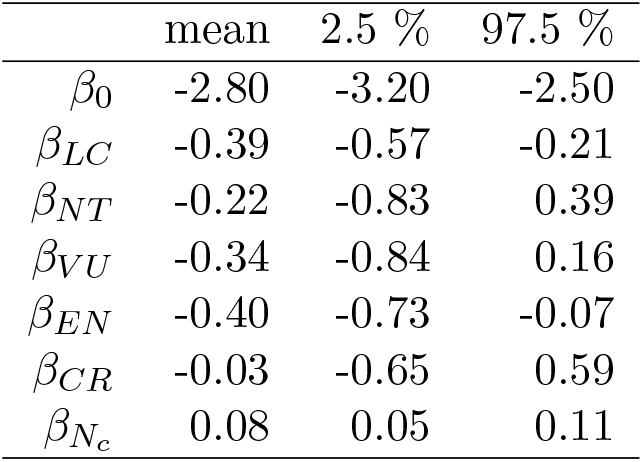
The regression estimates of full IUCN Red List population size model for diversity, 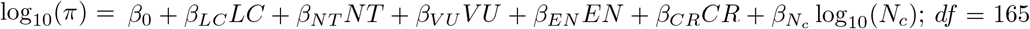. Using AIC to compare this full model to a reduced model of 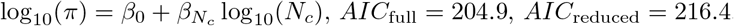.

|  | mean | 2.5 % | 97.5 % |
| --- | --- | --- | --- |
| $\beta_0$ | -2.80 | -3.20 | -2.50 |
| $\beta_{LC}$ | -0.39 | -0.57 | -0.21 |
| $\beta_{NT}$ | -0.22 | -0.83 | 0.39 |
| $\beta_{VU}$ | -0.34 | -0.84 | 0.16 |
| $\beta_{EN}$ | -0.40 | -0.73 | -0.07 |
| $\beta_{CR}$ | -0.03 | -0.65 | 0.59 |
| $\beta_{N_c}$ | 0.08 | 0.05 | 0.11 |

**Figure A1:**
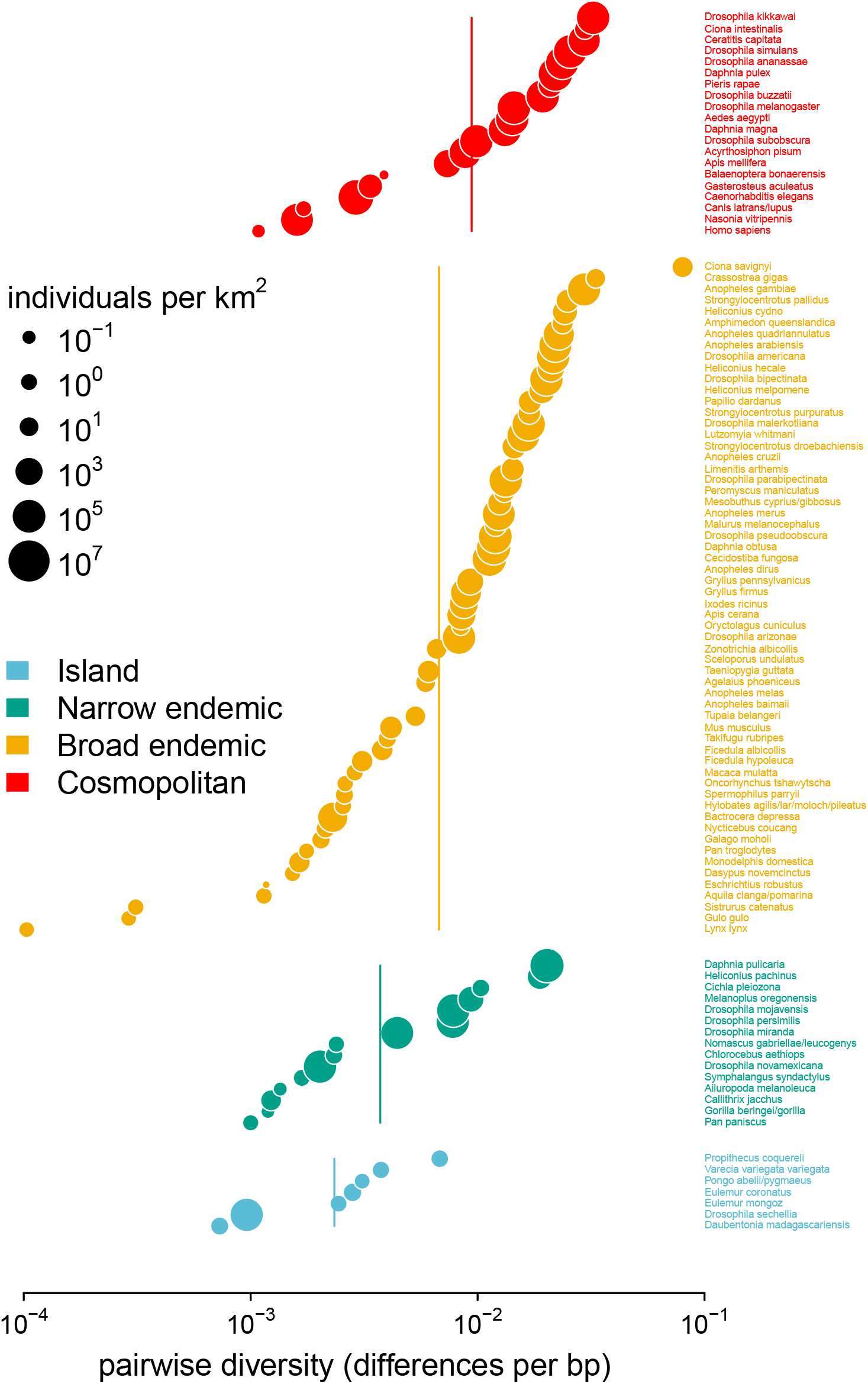
Pairwise diversity grouped by the range categories from Leffler et al. (2012), with point size indicating the predicted population density. The vertical lines are the range category group means.

**Figure A2:**
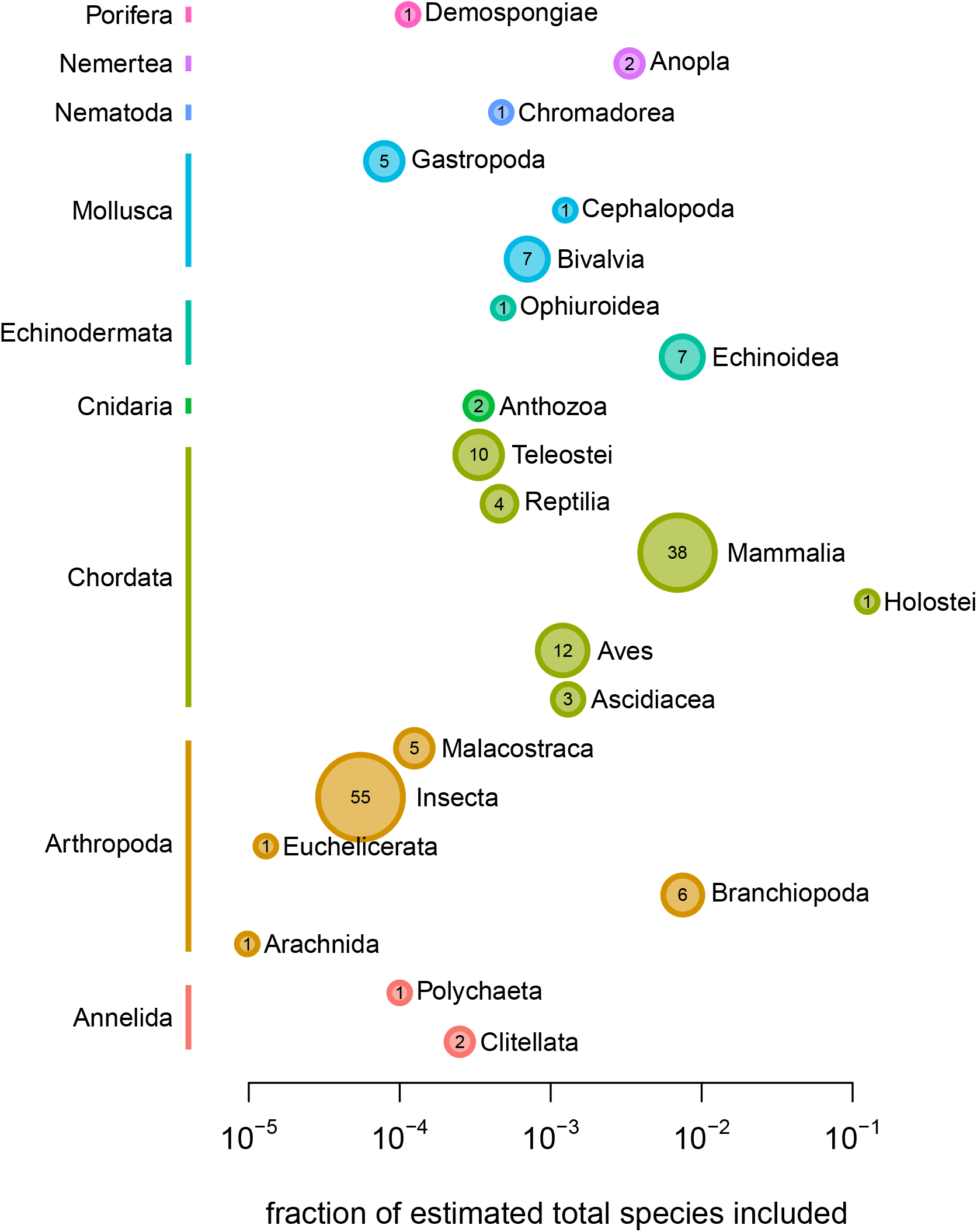
The fraction of total species on earth included in this study’s sample, per class. The color of the points represents phylum, and the size of the point represents the absolute number of species by class.

**Figure A3:**
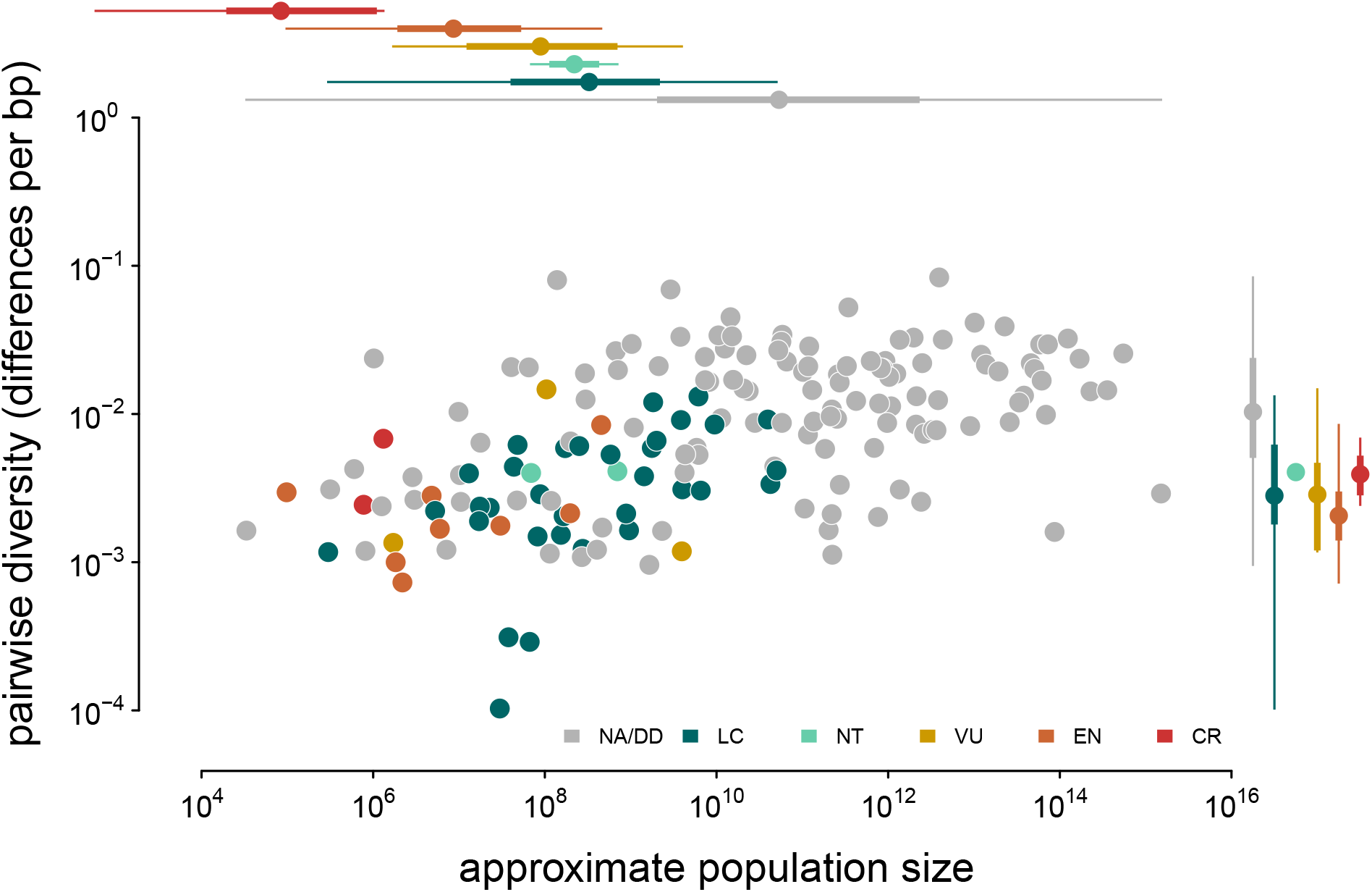
A version of Figure 2 with points colored by their IUCN Red List conservation status. Margin boxplots show the diversity and population size ranges (thin lines) and interquartile ranges (thick lines) for each category. NA/DD indicates no IUCN Red List entry, or Red List status Data Deficient; LC is Least Concern, NT is Near Threatened, VU is Vulnerable, EN is Endangered, and CR is Critically Endangered.

**Figure A4:**
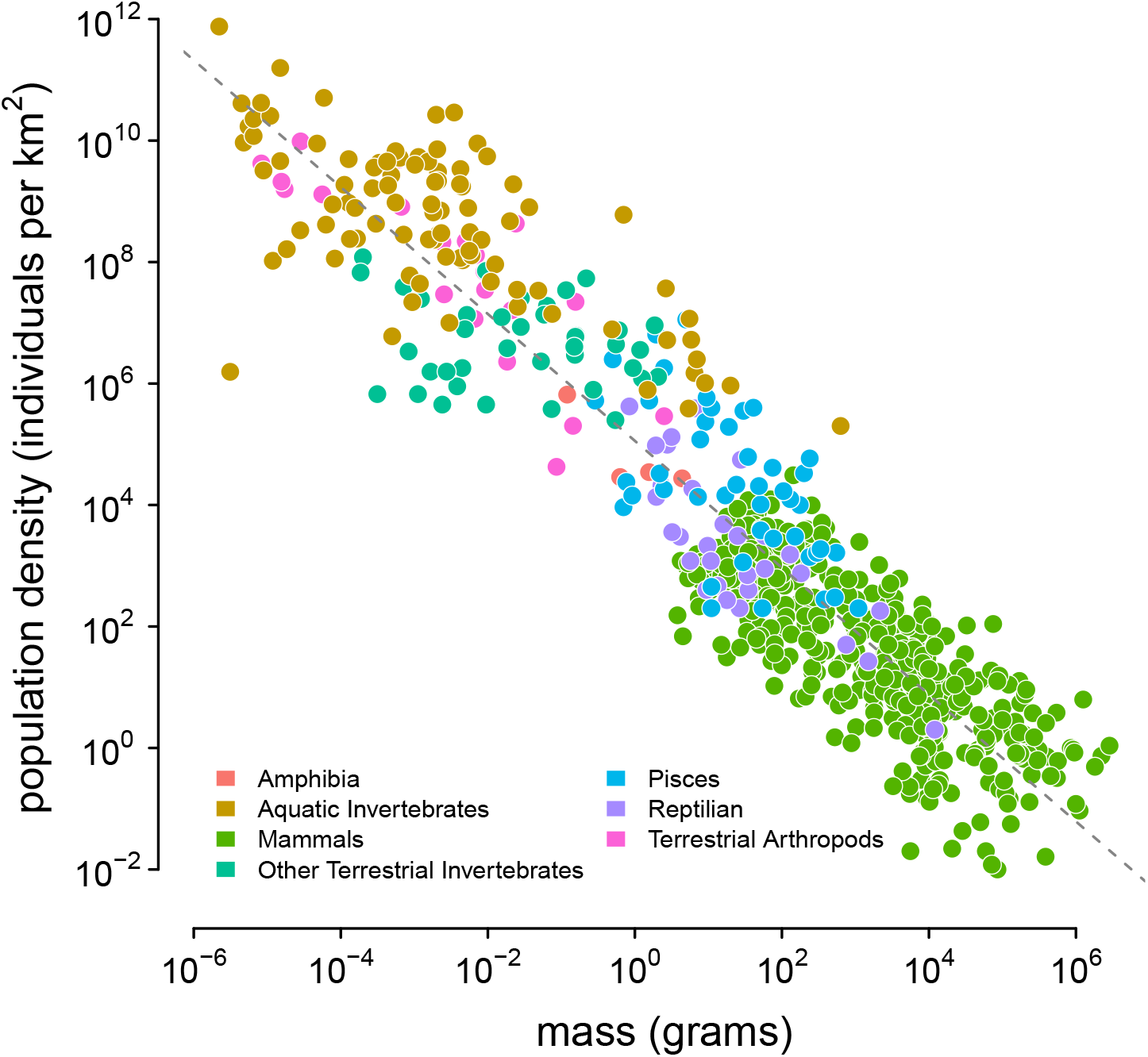
The appendix table of Damuth (1987); the color indicates Damuth’s original group labels. The dashed line was estimated using a lognormal regression model in Stan. References to each measurement are available in Damuth (1987).

**Figure A5:**
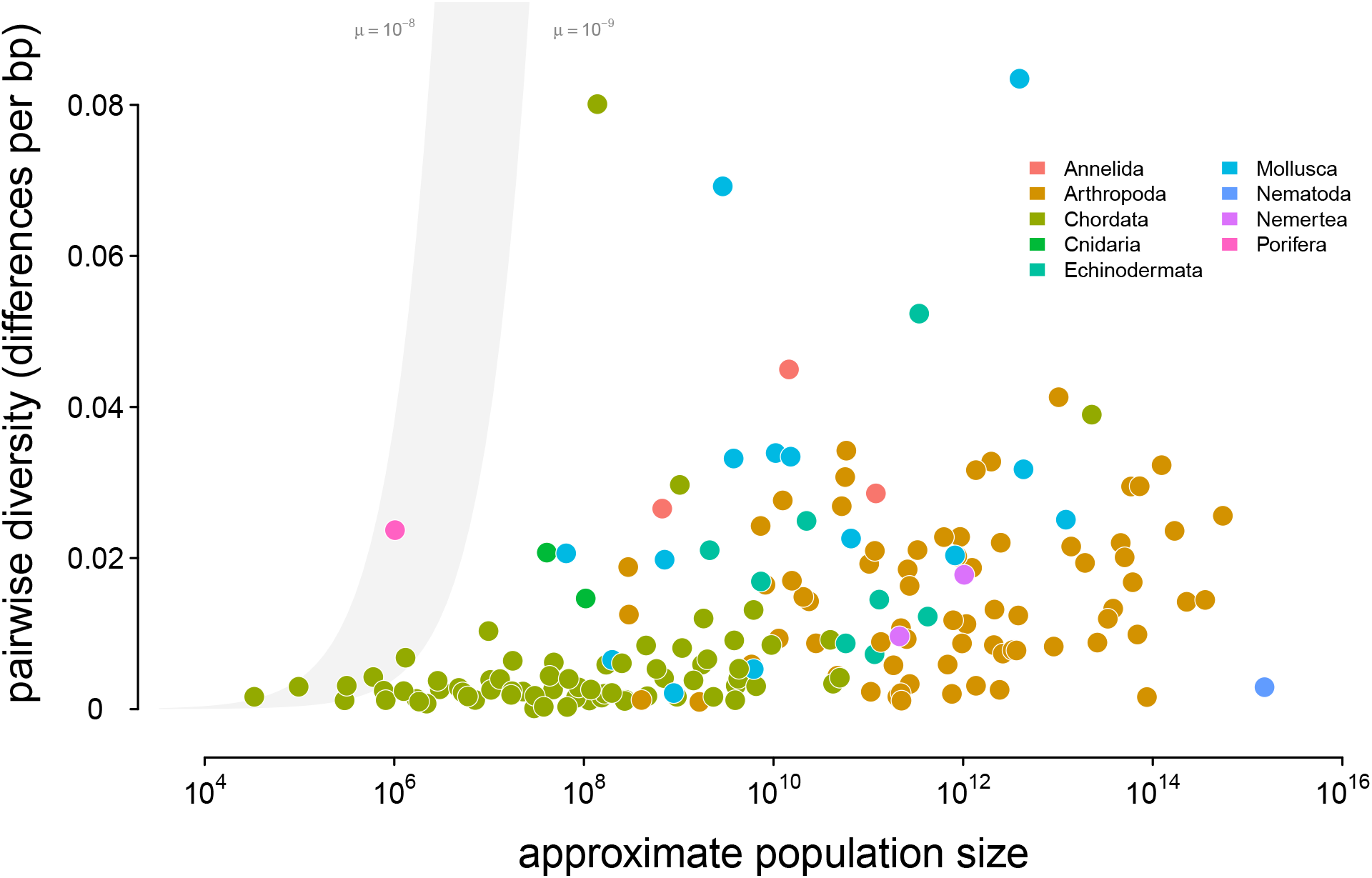
A version of Figure 2 with diversity on a linear, rather than log, scale. Points are colored by phylum, and the shaded region is the predicted neutral level of diversity assuming *N*_*e*_ = *N*_*c*_ with mutation range ranging between 10^*−*10^ ≤ *µ* ≤ 10^*−*8^.

**Figure A6:**
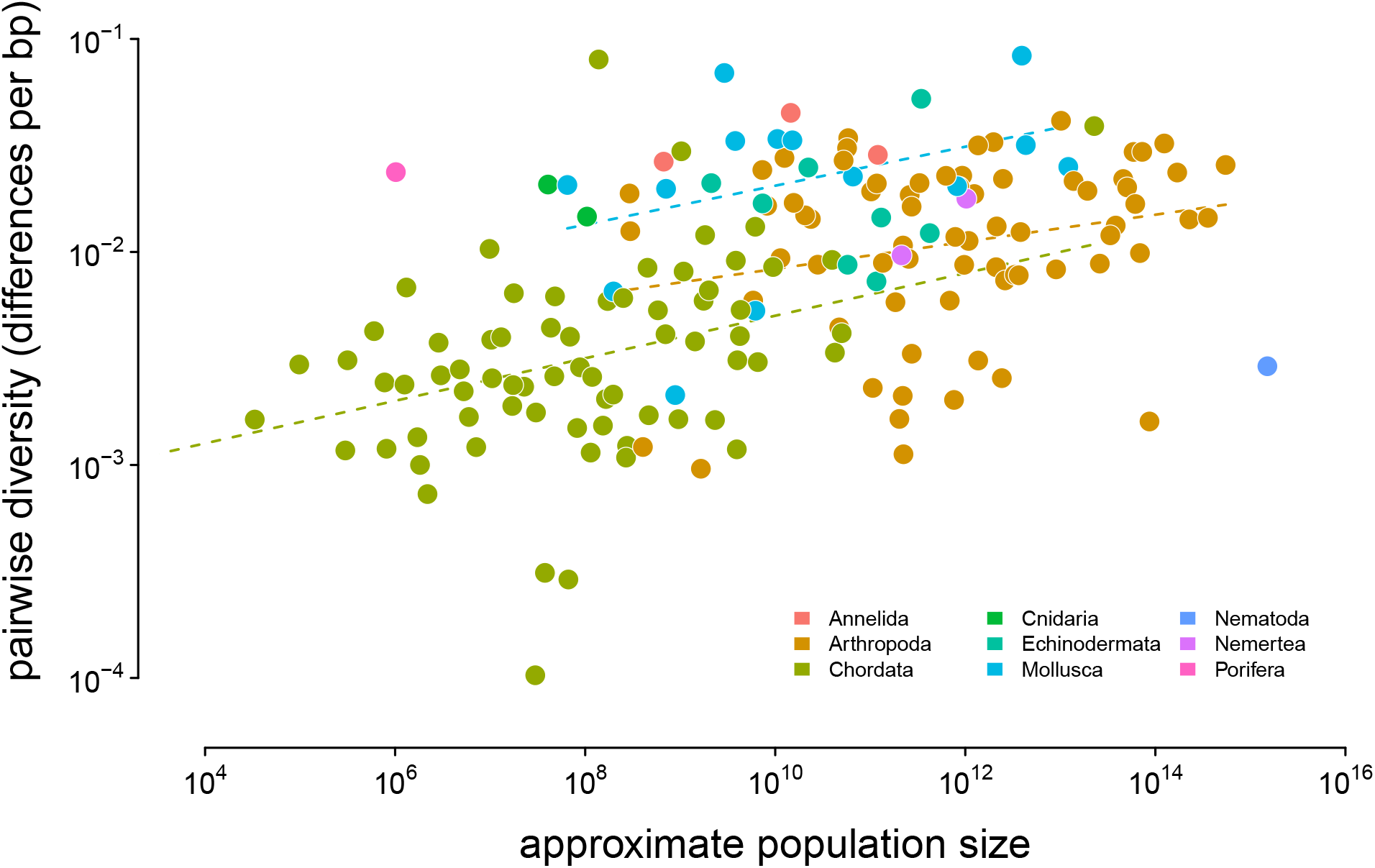
Diversity and approximate population size for 172 taxa, colored by phylum; the dashed lines indicate the non-phylogenetic OLS estimates of the relationship between population size and diversity grouped by phyla.

**Figure A7:**
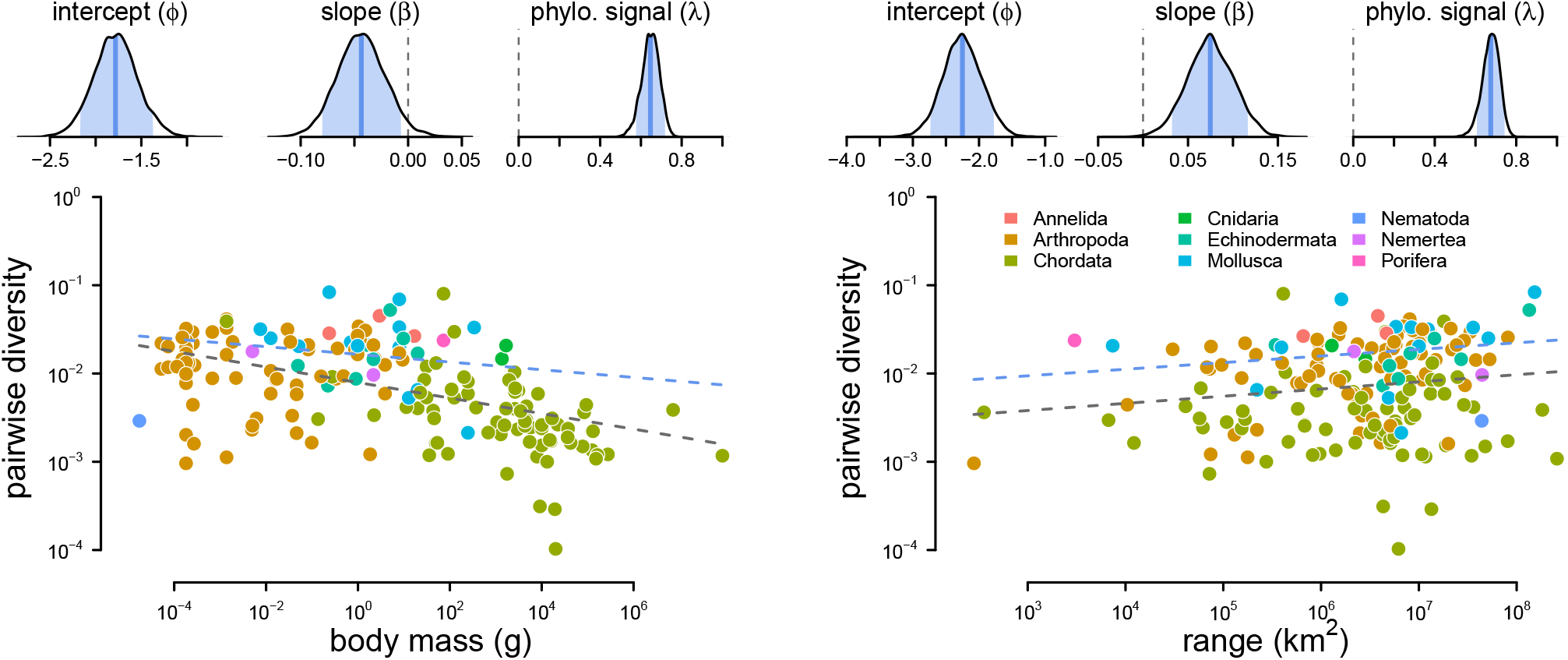
The relationship between diversity (differences per basepair) and body mass (left) and range (right) across 172 species. The top row are posterior distributions of parameters estimated using the phylogenetic mixed-effects model using 166 taxa in the synthetic phylogeny for the intercept, slope, and phylogenetic signal from the mixed-effects model. The bottom row contain each species as a point, colored by phyla. The gray dashed line is the non-phylogenetic standard regression estimate, and the blue dashed line is the relationship fit by the phylogenetic mixed-effects model.

**Figure A8:**
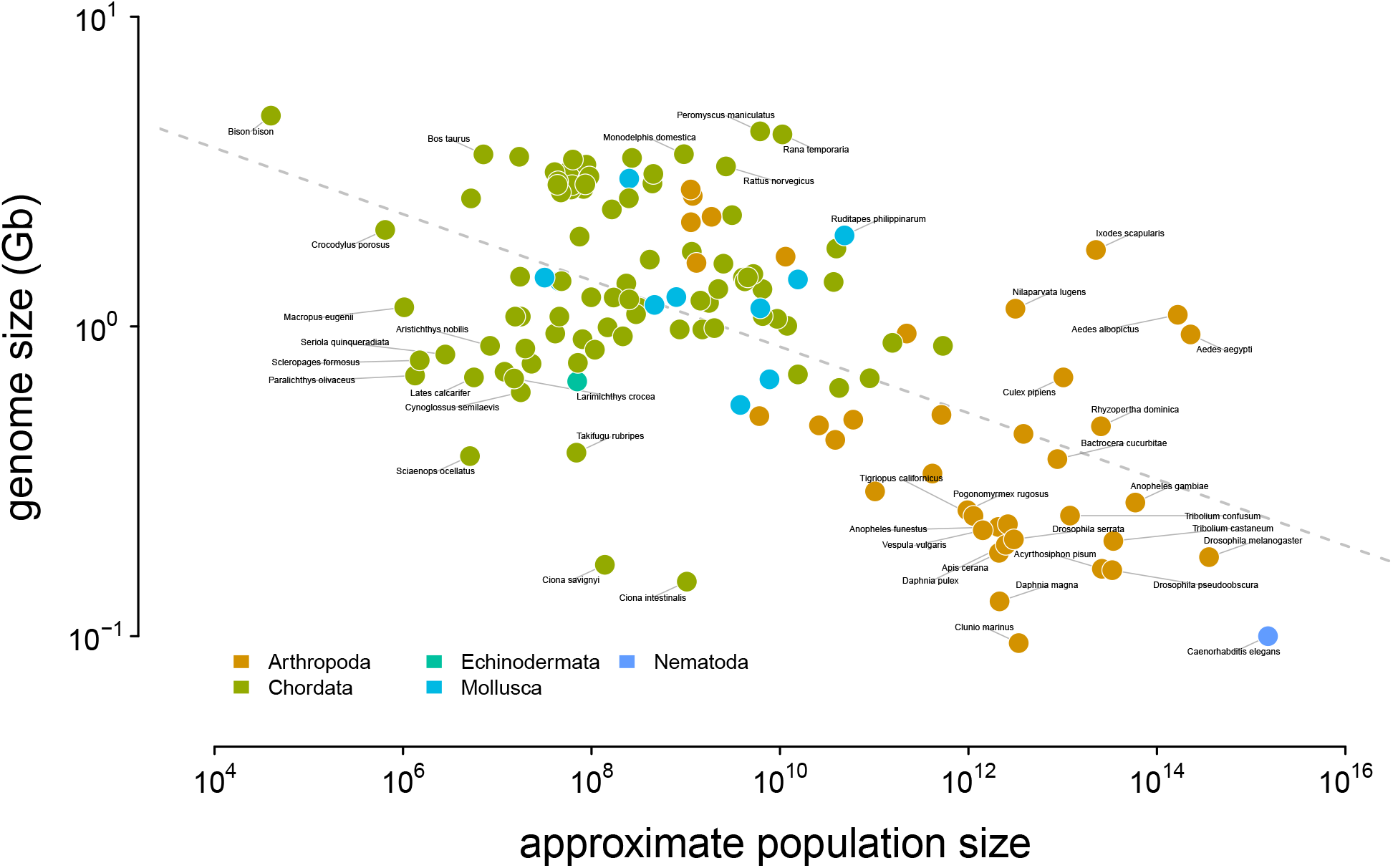
The relationship between genome size and approximate census population size. The dashed gray line indicates the OLS fit. Tiger salamander (*Ambystoma tigrinum*) was excluded because of its exceptionally large genome size (30Gbp).

**Figure A9:**
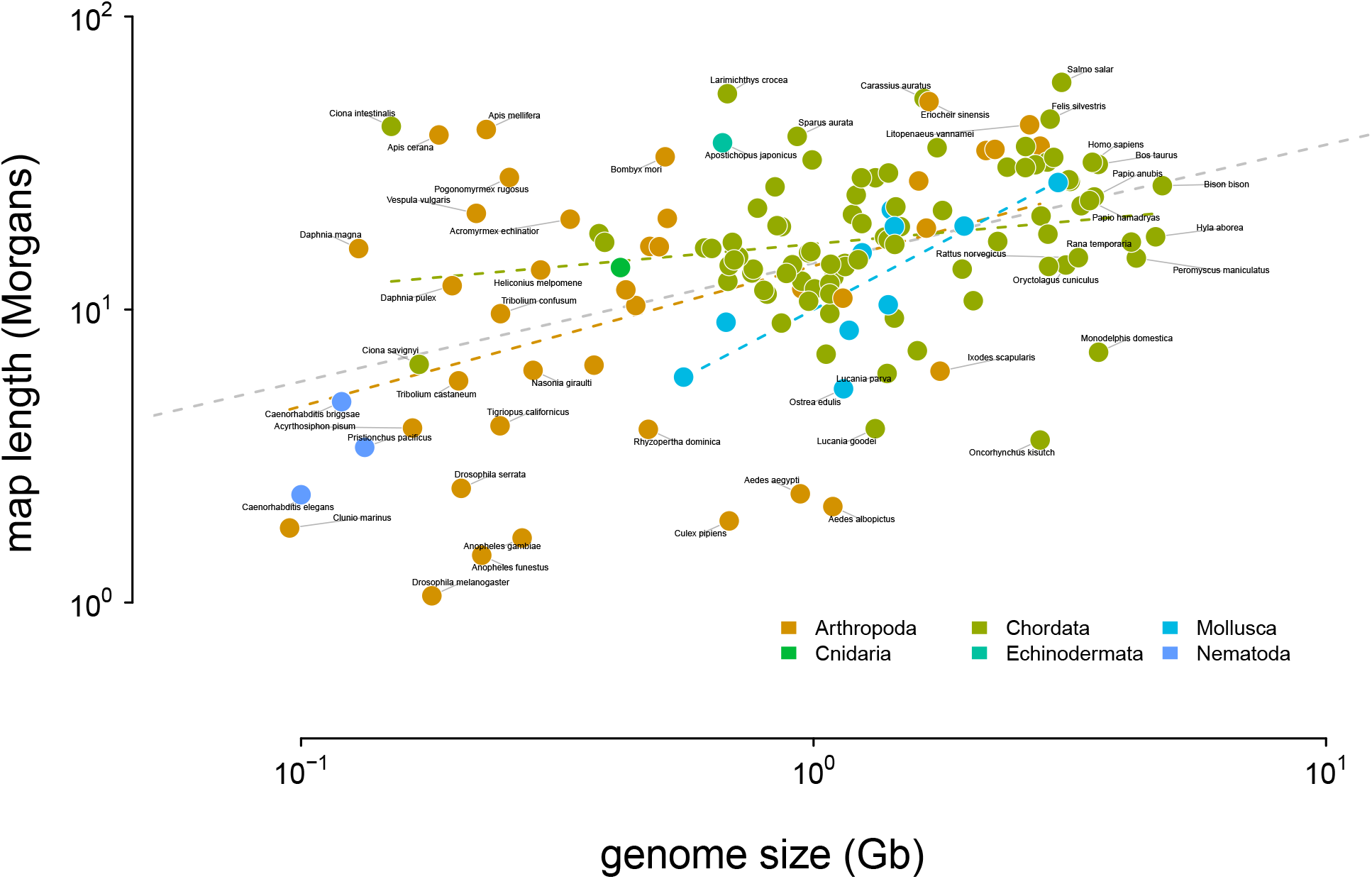
The relationship between genome size and recombination map length. The dashed gray line indicates the OLS fit for all taxa, and the dashed colored dashed lines indicate the linear relationship fit by phyla. Tiger salamander (*Ambystoma tigrinum*) was excluded because of its exceptionally large genome size (30Gbp).

**Figure A10:**
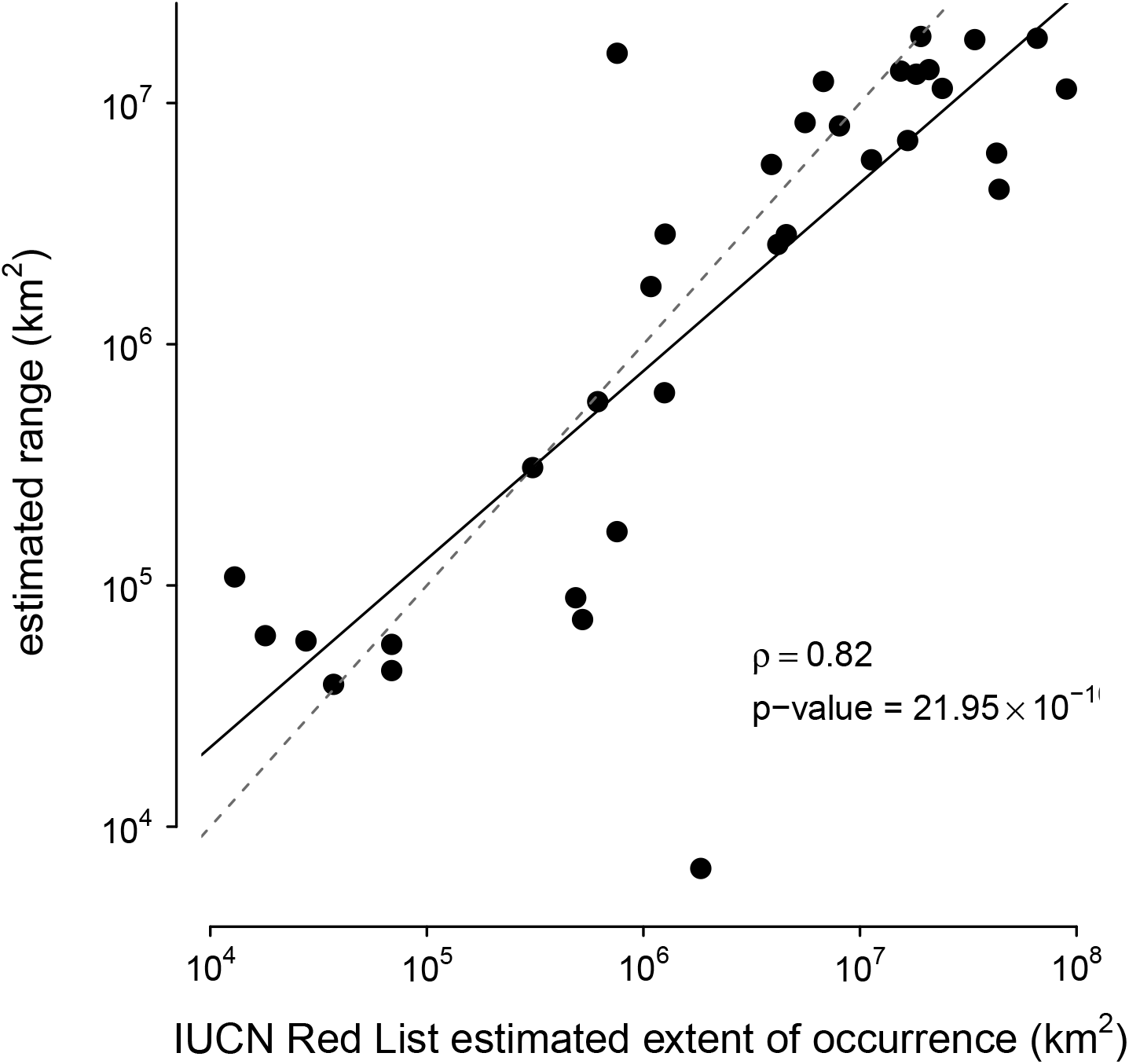
The correspondence between the ranges estimated with the alpha hull method applied to GBIF data used in this paper and IUCN Red List’s Extent of Occurrence for the subset of species in both datasets. Note that the IUCN Red List contains predominantly endangered species, which leads to ascertainment bias; still, the high correlation between the estimated ranges shows the alpha hull method works well.

**Figure A11:**
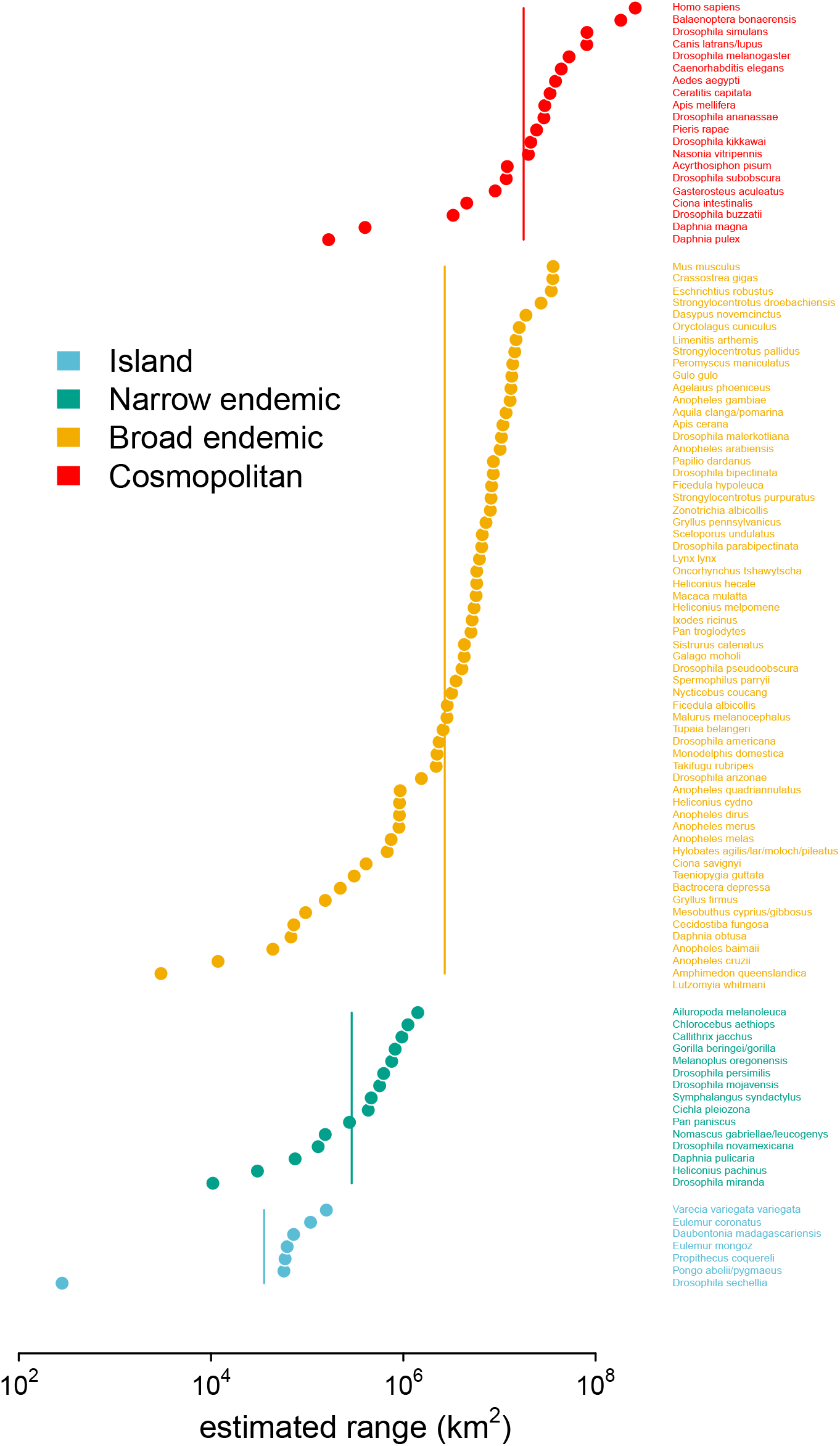
The estimated ranges using GBIF occurrence data, ordered within and colored by the original range category labels assigned in Leffler et al. (2012).

**Figure A12:**
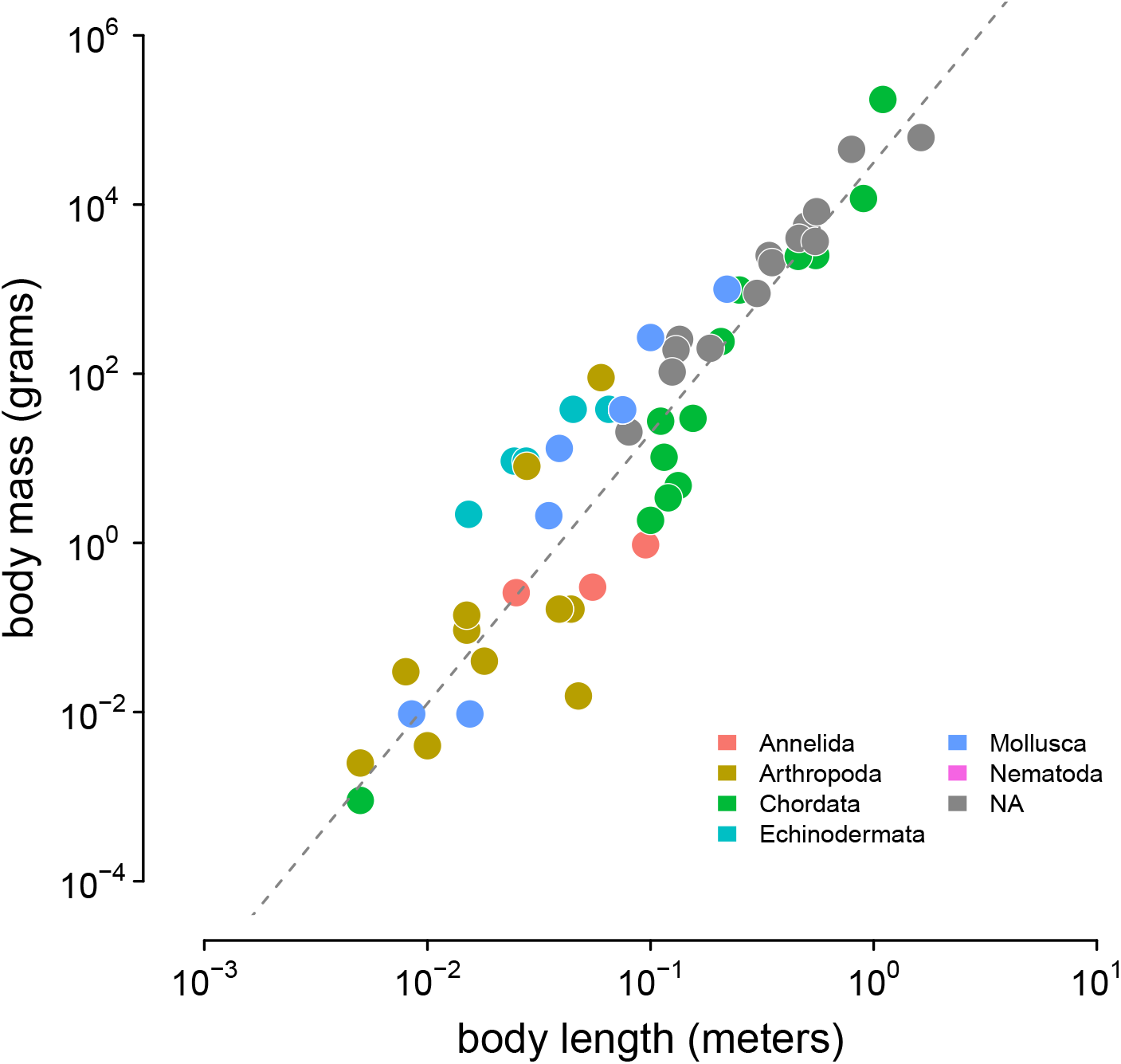
The relationship between body length (meters) and body mass (grams) in the Romiguier et al. (2014) data set, used to infer body masses for taxa. The gray dashed line is the line of best fit inferred using Stan.

**Figure A13:**
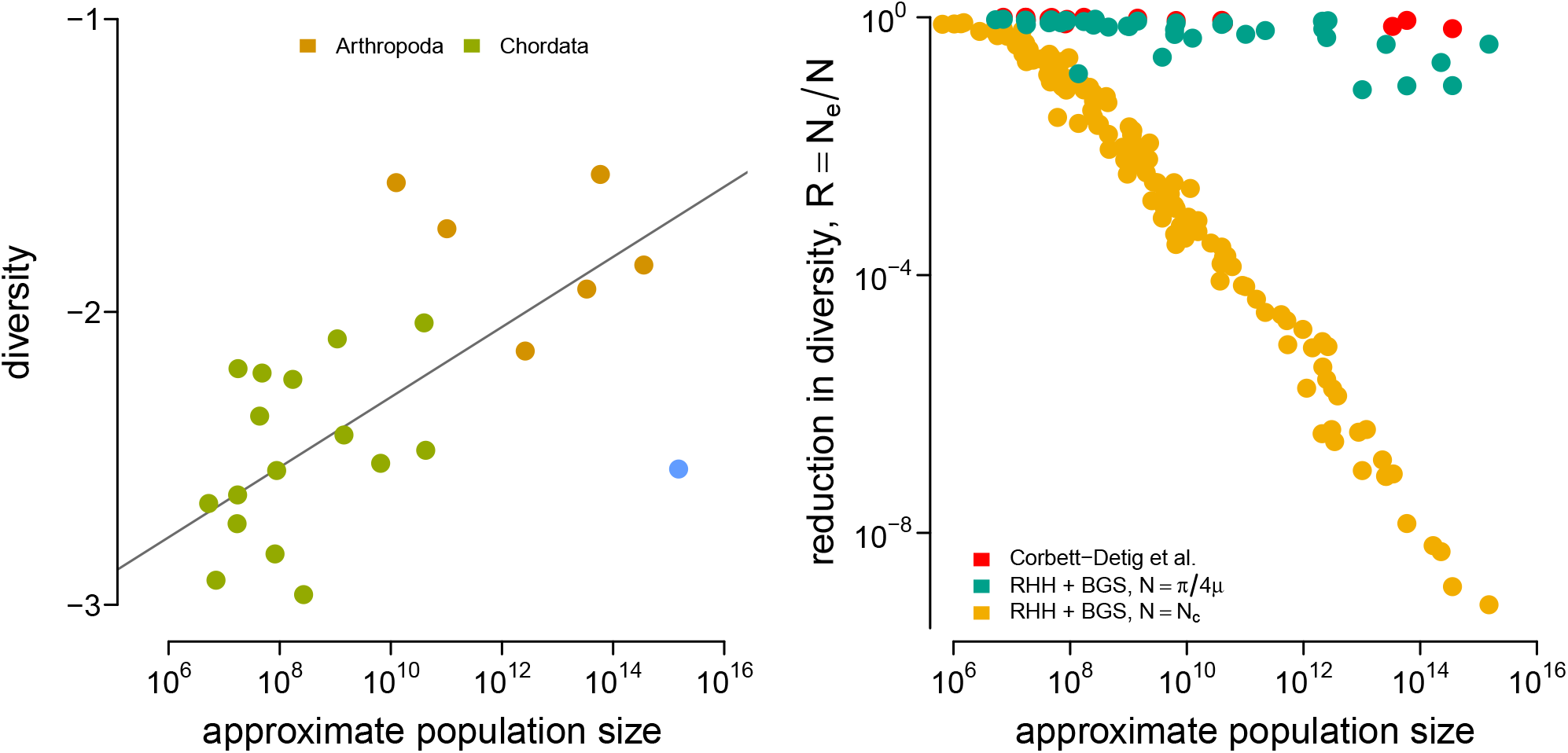
(A) The diversity data from Corbett-Detig et al. (2015) and the census population size estimated here for metazoan taxa. (B) The reductions in diversity, *R* = *N*_*e*_*/N*, plotted against census size across species. The red points are the reductions estimated by Corbett-Detig et al. (2015). This confirms Corbett-Detig et al.’s (2015) finding that the impact of selection (*I* = 1 − *R*) increases with census population size (though, in the original paper size body size and range were used as separate proxy variables for census population size). The green and red points are the predicted reduction in diversity under the recurrent hitchhiking (RHH) and background selection (BGS) model using the *Drosophila melanogaster* parameters as described in the main text. The reduction in the diversity due to sweeps, from Equation (1), is determined by the term 2*NS*. Green points treat *N* as the implied effective population size from diversity 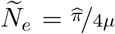, assuming *µ* = 10^*−*9^. Yellow points treat *N* as the census size, *N* = *N*_*c*_. Overall, using the census size, e.g. 2*N*_*c*_*S*, leads to reductions in diversity that far exceed the empirical estimates of Corbett-Detig et al. and reasonable model-based predictions from *Ñ*_*e*_.

**Figure A14:**
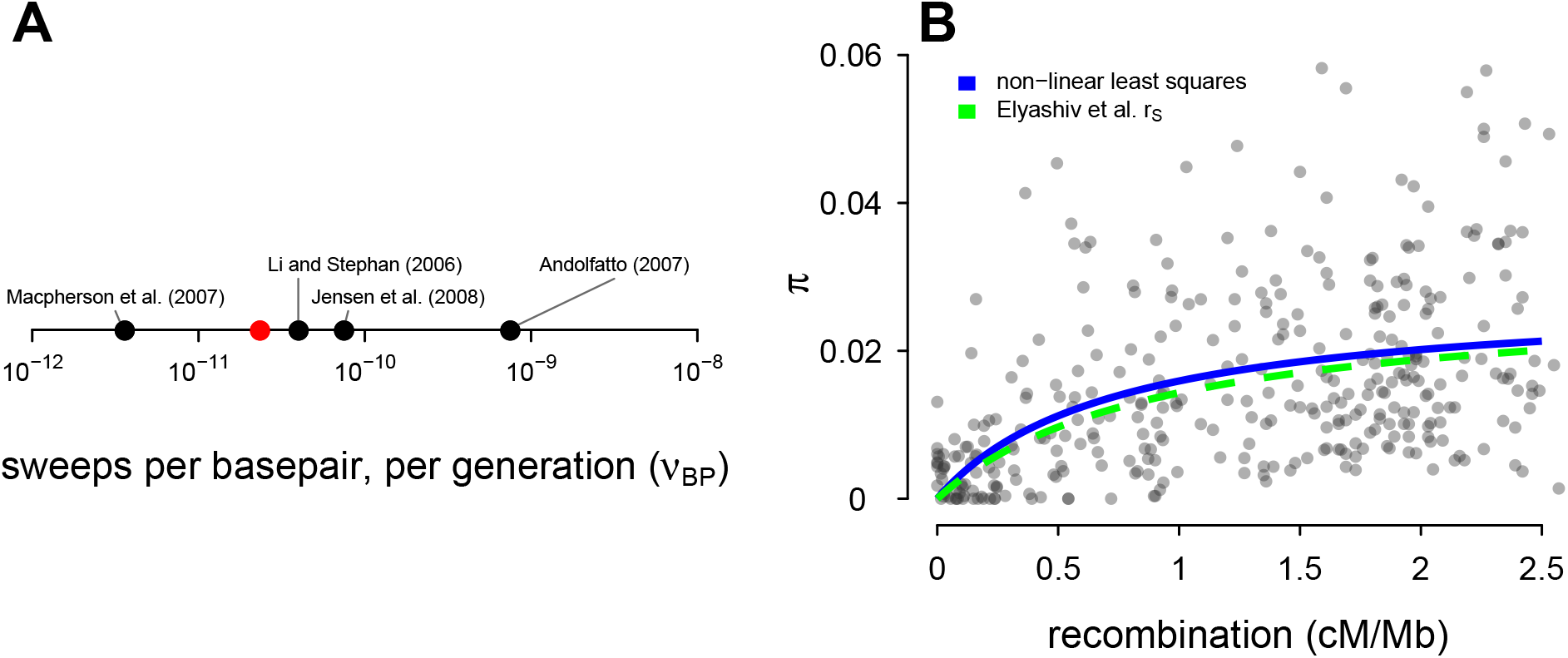
(A) The estimate of the number of sweeps per basepair, per genome (*ν*_BP_) from Table 2 of Elyashiv et al. (2016) (the studies included are Andolfatto 2007; Li and Stephan 2006; Macpherson et al. 2007 and Jensen et al. 2008); the red point is my estimate used in this paper. (B) Points are the data from Shapiro et al. (2007). The blue line is the non-linear least squares fit to the data, and the green dashed line is the sweep model parameterized by the genome-wide average sweep coalescent rate 2*NS* ≈ 0.92 from the classic sweep and background selection model of Elyashiv et al. (2016) (*r*_*s*_ in Supplementary Table S6).

**Figure A15:**
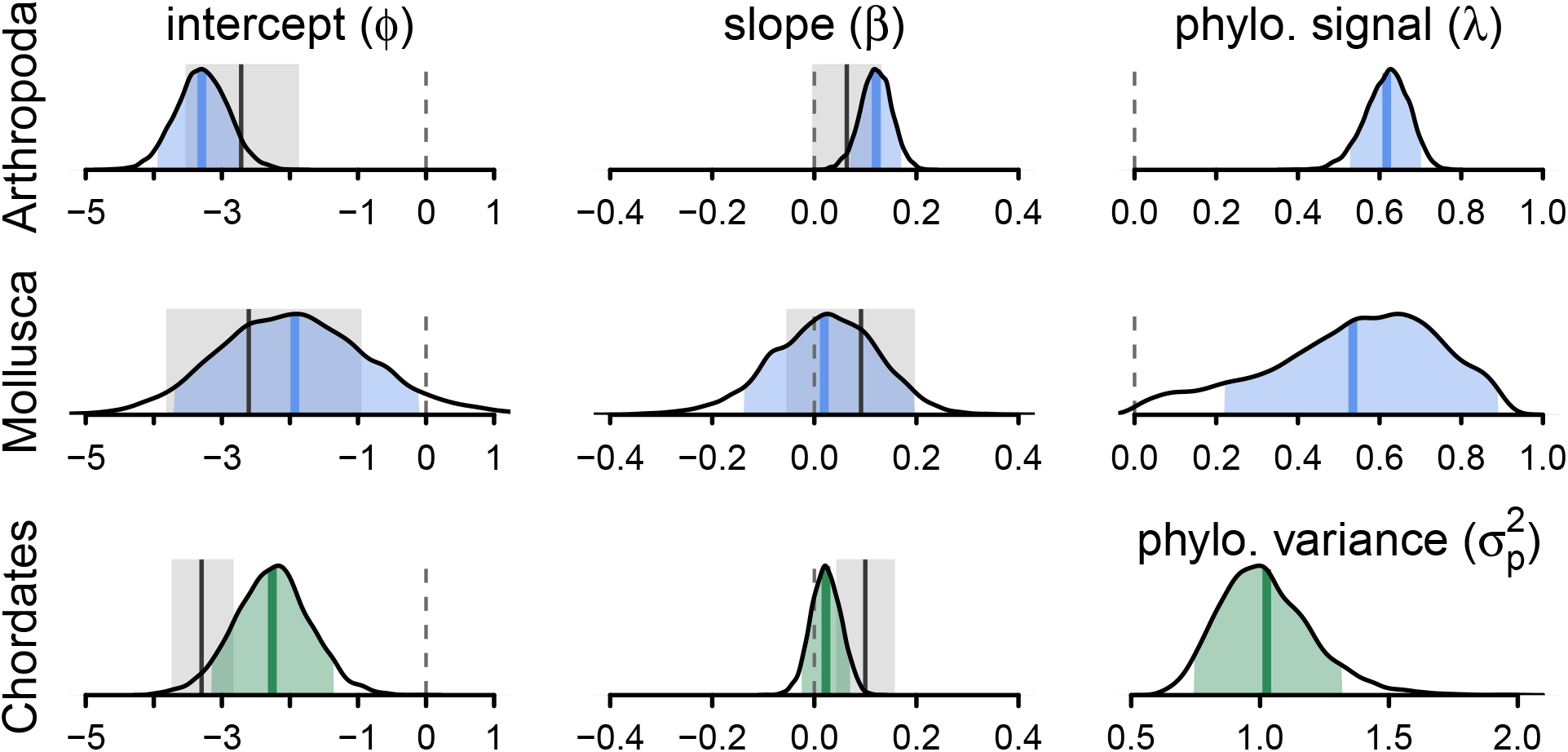
The posterior distributions for the parameters of the phylogenetic mixed-effects model of diversity and population size (this is analogous to Figure 3B) fit separately on chordates (*n* = 68), molluscs (*n* = 13), and arthropods (*n* = 68). The phylogenetic mixed-effects model for chordates indicated the best-fitting model had no residual variance 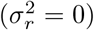, so an alternate model without this variance component was used to ensure proper convergence; this model is shown in green. The light blue (green) shaded regions are the 90% credible intervals, the blue (green) lines the posterior averages, the gray shaded regions the OLS bootstrap 95% confidence intervals, and the gray lines the OLS estimate. Note that unlike Figure 3, the OLS estimate uses all taxa, not just those present in the phylogeny, since splitting the data by phyla reduces sample sizes (OLS with just the subset of taxa in the phylogeny is not significant for either chordates and arthropods). The vertical dashed gray line indicates zero.

**Figure A16.**
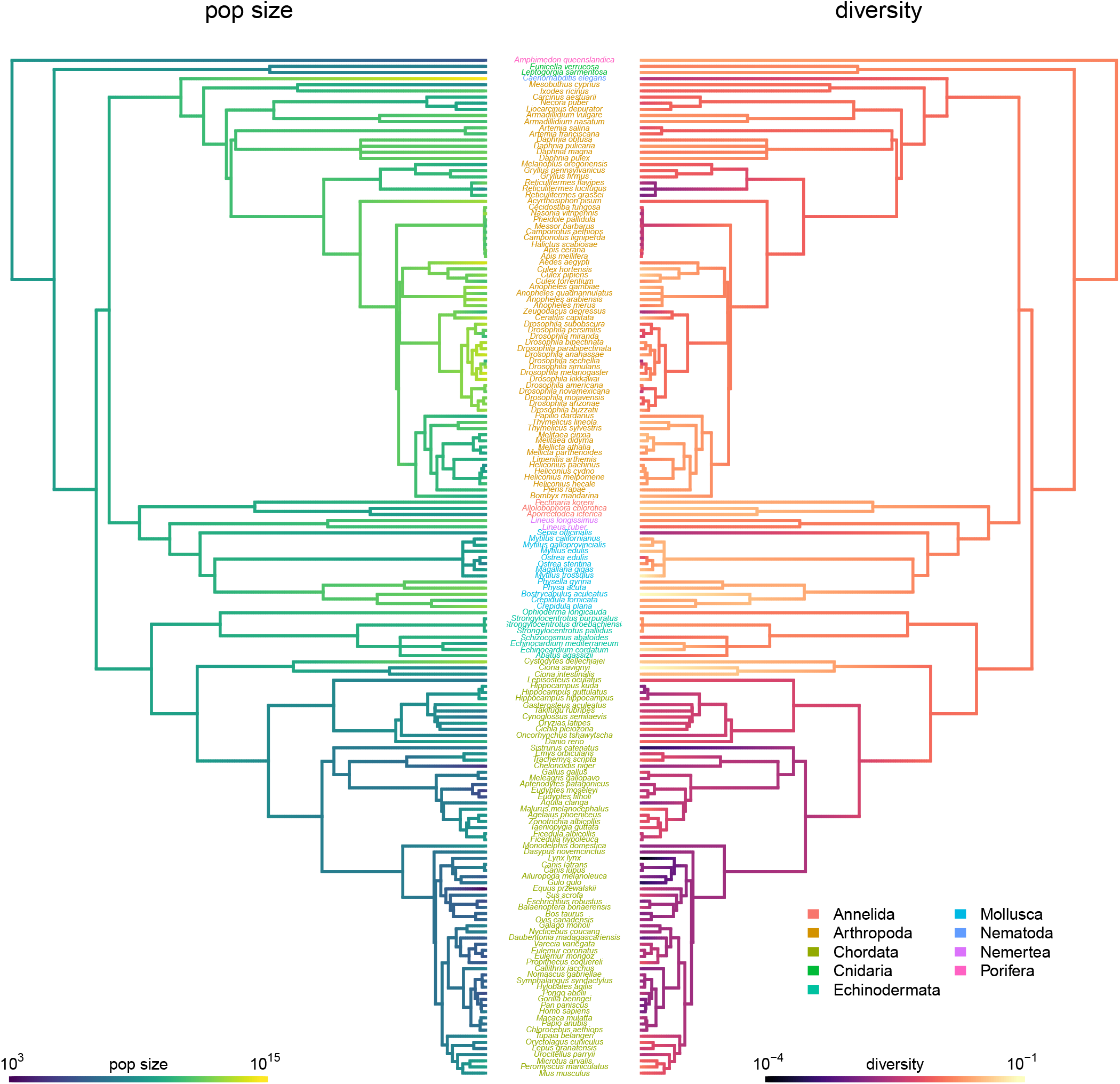

**Figure A17.**
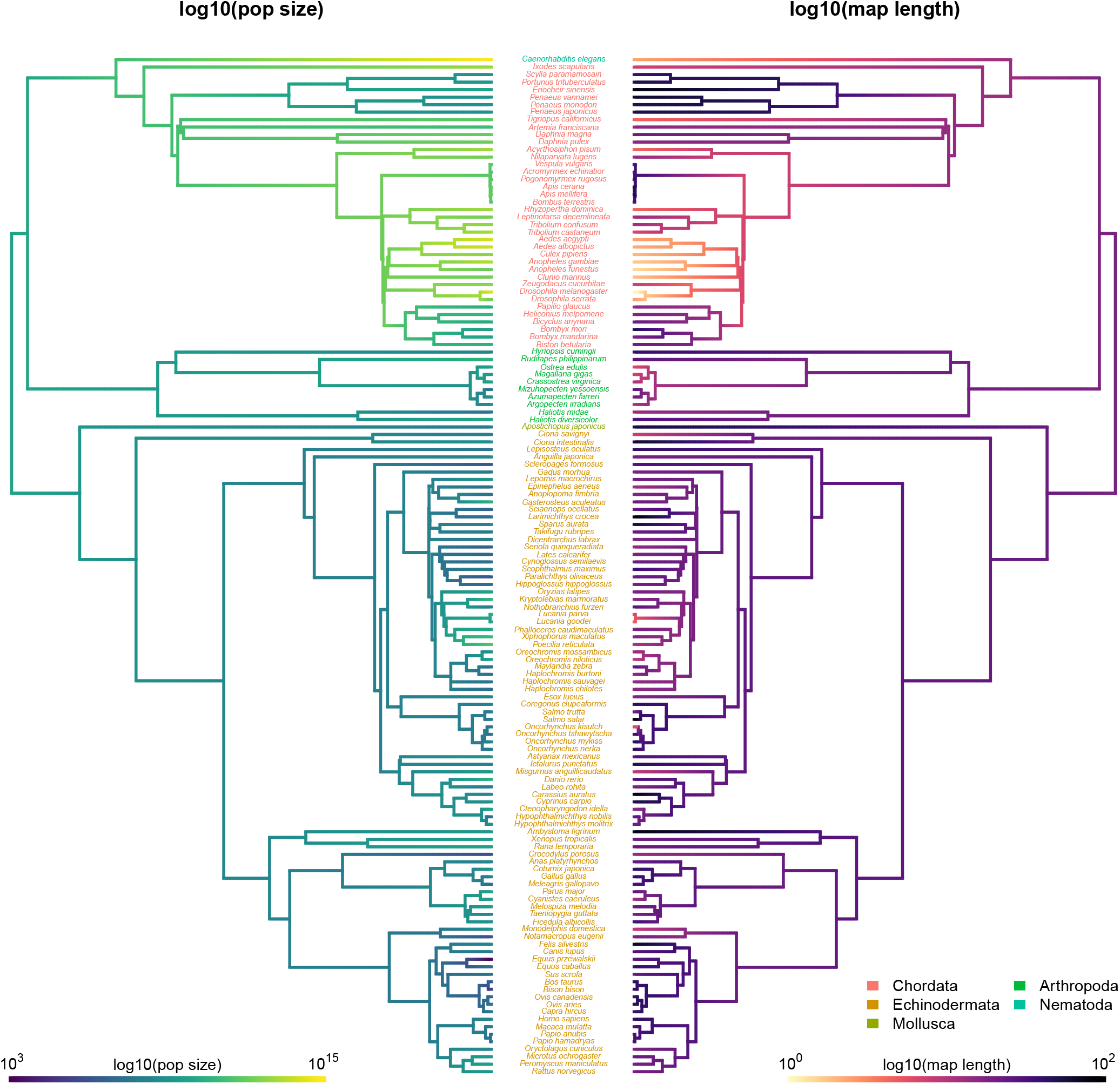

