## Supplementary material for "Natural Selection is Unlikely to Explain Why Species Get a Thin Slice of *π*": A PDF of the differences between this version and the last version.

### Why do species get a thin slice of $\pi$? Revisiting Lewontin's Paradox of Variation


Vince Buffalo

University of Oregon, Institute of Ecology and Evolution
Eugene, Oregon



May 26, 2021


#### Abstract

~~Under neutral theory , the level of polymorphism in an equilibrium population is expected to increase~~ Neutral theory predicts that genetic diversity increases with population size~~. However,~~ , yet observed levels of diversity across metazoans vary only two orders of magnitude ~~, while census population sizes ($N_c$) are expected to~~ while population sizes vary over several. This unexpectedly narrow range of diversity is ~~a longstanding enigma in evolutionary genetics~~ known as Lewontin's Paradox of Variation (1974). ~~Since Lewontin's observation, it has been argued that~~ While some have suggested selection constrains diversity~~across species, yet~~ , tests of this hypothesis seem to fall short~~of explaining the orders-of-magnitude reduction in diversity observed in nature. In this work~~. Here, I revisit Lewontin's Paradox ~~and~~ to assess whether current models of linked selection are ~~likely to constrain~~ capable of reducing diversity to this extent. To quantify the discrepancy between pairwise diversity and census population sizes across species, I combine ~~genetic data~~ previously-published estimates of pairwise diversity from 172 metazoan taxa with estimates of census sizes~~from geographic occurrence data and population densities estimated from body mass. Next, I fit the relationship between previously-published estimates of genomic diversity and these approximate census sizes to quantify Lewontin's Paradox. While previous across-taxa population genetic studies have avoided accounting for phylogenetic non-independence, I use phylogenetic comparative methods to investigate the diversity-census size relationship, estimate phylogenetic signal , and explore how diversity changes along the phylogeny. I consider whether the reduction in diversity predicted by models of recurrent hitchhiking and background selection could explain the observed pattern of diversity across species. Since the impact of linked selection is mediated by recombination map length, I also investigate how map lengths vary with census sizes. I find species with large census sizes have shorter map lengths, leading these species to~~. Using phylogenetic comparative methods, I show this relationship is significant accounting for phylogeny, but with high phylogenetic signal and evidence that some lineages experience shifts in the evolutionary rate of diversity deep in the past. Additionally, I find a negative relationship between recombination map length and census size, suggesting abundant species have less recombination and experience greater reductions in diversity due to linked selection. ~~Even after using high estimates of the strength of sweeps and background selection , I find linked selection likely cannot explain the shortfall between predicted and observed diversity levels across metazoan species. Furthermore, the predicted diversity under linked selection does not fit the observed diversity-census-size relationship, implying that processes other than background selection and recurrent hitchhiking must be limiting diversity.~~ However, I show that even using strong selection parameter estimates, models of linked selection are unlikely to explain the observed relationship between diversity and census sizes across species.

Past work looking at the relationship between $\pi$ and $N_c$ has ~~largely ignored~~ been unable to fully account for phylogenetic non-independence across taxa (Felsenstein 1985). To address this shortcoming, I ~~account for phylogenetic non-independence across taxa using a synthetic time-calibrated phylogeny and phylogenetic~~ use phylogenetic comparative methods (PCMs) with a synthetic time-calibrated phylogeny to account for shared phylogenetic history. Moreover, ~~Lynch (2011) has argued that~~ it is disputed whether considering phylogenetic non-independence is necessary in population genetics, since coalescent times are much less than divergence times ~~, considering phylogenetic non-independence is unnecessary for traits like effective population size~~(Lynch 2011; Whitney and Garland 2010). Using PCMs, I ~~test this conjecture by estimating~~ address this by estimating the degree of phylogenetic signal in the diversity census size relationship, and investigating how these traits evolve along the phylogeny.

Finally, I explore whether the predicted reductions of diversity under background selection and recurrent hitchhiking are sufficiently strong ~~enough~~ to resolve Lewontin's Paradox. These predicted reductions in diversity across species are ~~generously estimated using selection strength~~ estimated using strong selection parameters from *Drosophila melanogaster*, a species known to be strongly affected by linked selection. Given the effects of linked selection are mediated by recombination map length, I investigate how recombination map lengths vary with census population size using data from a previously-published survey (Stapley et al. 2017). I find map lengths are typically shorter in large–census-size species, increasing the effects of linked selection in these species, which might further decouple diversity from census size. Still, I find the combined impact of these selection models with available parameter estimates falls short in explaining Lewontin's Paradox, and discuss future avenues through which the Paradox of Variation could be fully resolved.

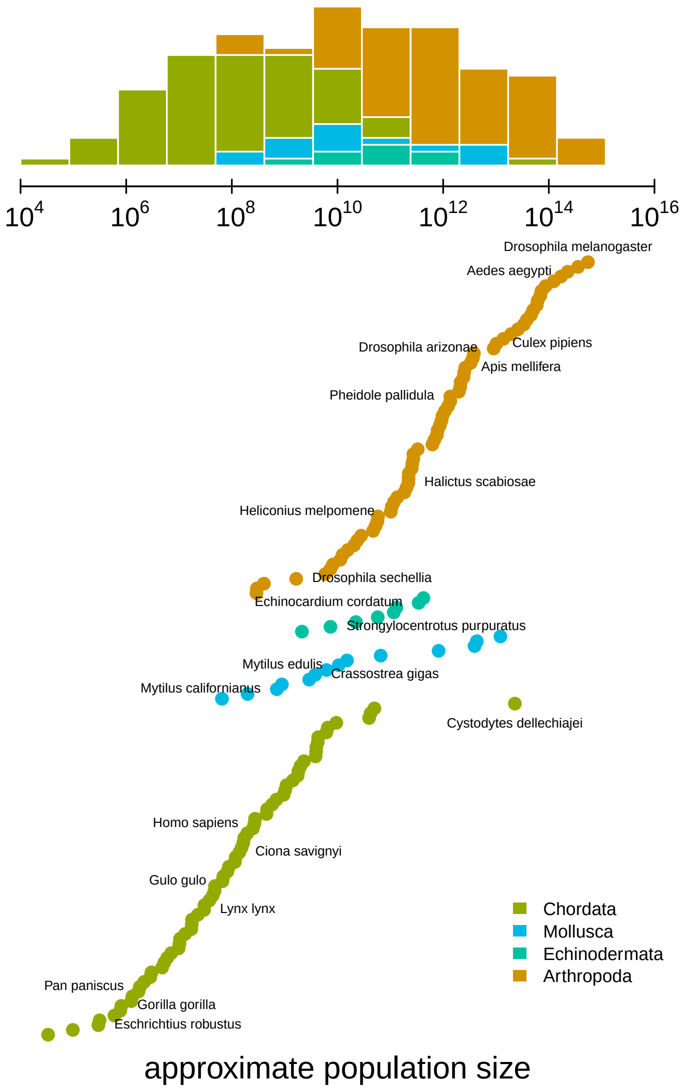


**Figure 1:** The distribution of approximate census population sizes estimated by this study. Some phyla containing few species were excluded for clarity.

### Results

#### Estimates of Census Population Size

A major impediment in quantifying Lewontin's Paradox has been estimating census population sizes across many taxa, especially for extremely abundant, cosmopolitan species that define the upper limit of ranges. Previous work has surveyed the literature for census size estimates (Frankham 1996; Nei and Graur 1984; Soulé 1976), or used range~~and body size~~, body size, or qualitative categories as proxies for census size (Corbett-Detig et al. 2015; Leffler et al. 2012). To quantify the relationship between genomic estimates of diversity and census population sizes, I first approximate census population sizes for 172 metazoan taxa (Figure 1). My approach predicts population densities from body sizes using a previously-observed linear relationship that holds across metazoans (Supplementary Figure S11; Damuth 1981, 1987). Then, from geographic occurrence data, I estimate range sizes. Finally, I estimate population size as the product of these predicted densities and range estimates (see Methods: *Macroecological Estimates of Population Size*). Note that the relationship between population density and body size is driven by energy budgets, and thus reflects macroecological equilibria (Damuth 1987)~~; consequently~~. Consequently, population sizes are underestimated for taxa like humans and their domesticated species~~are underestimated. ~~, and overestimated for species with anthropogenically reduced densities or fragmented ranges. For example, the population size of *Lynx lynx* is likely around 50,000 (IUCN 2020) which is around two orders of magnitude smaller than my estimate. Additionally, the range size estimates do not consider whether an area has unsuitable habitat, and thus may be overestimated for species with particular niches or patchy habitats. While these methods to estimate census size are crude and approximate, they can be efficiently calculated for numerous taxa and are sufficient to estimate the scale of Lewontin's Paradox (see Supplementary Information *Population Size Validation* for more on validation based on biomass and other approaches).

#### Quantifying Lewontin's Paradox

To determine which ecological or evolutionary processes could decouple diversity from census population size, we first need to quantify this relationship across a wide variety of taxa. Previous work has found there is a significant relationship between heterozygosity and the logarithm of population size or range size, but these studies relied on heterozygosity measured from allozyme data (Frankham 1996; Leffler et al. 2012; Nei and Graur 1984; Soulé 1976). Here, I confirm these findings using pairwise diversity estimates from genomic sequence data and the estimated census sizes (Figure 2). The pairwise diversity estimates are from three sources: Leffler et al. (2012), Corbett-Detig et al. (2015), and Romiguier et al. (2014), and are predominantly from either synonymous or non-coding DNA (see Methods: *Diversity and Map Length Data*). Overall, an ordinary least squares (OLS) relationship on a log-log scale fits the data well (Figure 2). The OLS slope estimate is significant and implies ~~an increase in diversity of 0.09~~ a 13% percent increase in differences per basepair for every order of magnitude census size grows (95% confidence interval ~~[0.08, 0.12]~~[12%, 14%], adjusted $R^2 = 0.26$; see also the OLS fit per-phyla, Supplementary Figure S5).

Notably, this relationship has few outliers and is relatively homoscedastic. This is in part because of the log-log scale, in contrast to previous work (Nei and Graur 1984; Soulé 1976); see Supplementary Figure S4 for a version on a log-linear scale. However, it is noteworthy that few taxa have diversity estimates below $10^{-3.5}$ differences per basepair. Those that do, lynx (*Lynx Lynx*), wolverine (*Gulo gulo*), and Massasauga rattlesnake (*Sistrurus catenatus*) face habitat fragmentation and declining population sizes. These three species are all in the IUCN Red List, but are listed as least concern (though their presence in the Red List indicates they are of conservation interest). ~~I also investigated the relationship between species'~~ In Supplementary Information Section *Diversity and IUCN Red List Status*, I explore more about the relationships between IUCN Red List ~~categories (an ordinal scale of how threatened a species is) and both diversity~~status, diversity, and population size~~, finding that species categorized as more threatened have both smaller population sizes and reduced diversity, compared to non-threatened species (Figure ??) consistent with past work . A linear model of diversity regressed on population size has lower AIC when the IUCN Red List categories are included, and the estimates of the effect of IUCN status are all negative on diversity, though not all are significant in part because some categories have three or fewer species~~

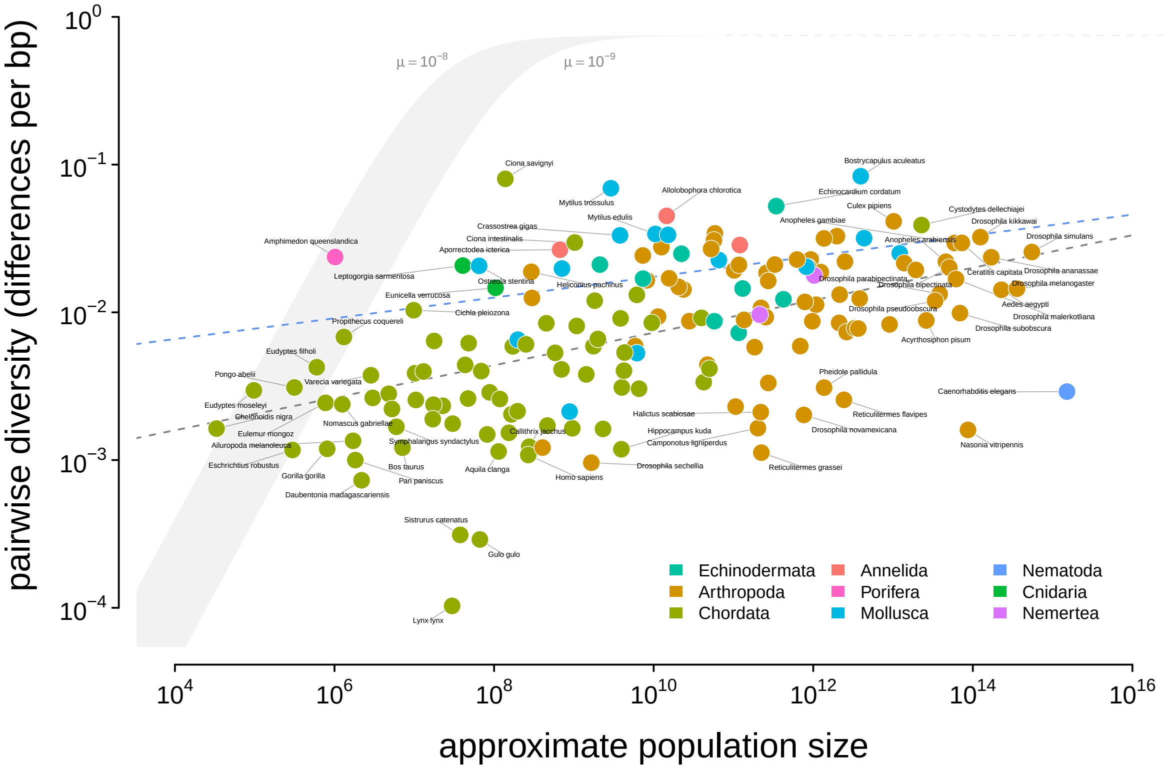


**Figure 2:** An illustration of Lewontin's Paradox of Variation. Pairwise diversity (data from Leffler et al. 2012, Corbett-Detig et al. 2015, and Romiguier et al. 2014), which varies around three orders of magnitude, shows a weak relationship with approximate population size, which varies over 12 orders of magnitude. The shaded curve shows the range of expected neutral diversity if $N_e$ were to equal $N_c$ under the four-alleles model, $\log_{10}(\pi) = \log_{10}(\theta) - \log_{10}(1 + 4\theta/3)$ where $\theta = 4N_c\mu$, for two mutation rates, $\mu = 10^{-8}$ and $\mu = 10^{-9}$, and the light gray dashed line represents the maximum pairwise diversity under the four alleles model. The dark gray dashed line is the OLS regression fit, and the blue dashed line is the regression fit using a phylogenetic mixed-effects model. Points are colored by phylum. The species *Equus ferus przewalskii* ($N_c \approx 10^3$ and $\pi = 3.6 \times 10^{-3}$) was an outlier and excluded from this figure for visual clarity.

~~(Supplementary Table S3).~~ .

~~A version of Figure 2 with points colored by their IUCN Red List conservation status. Margin boxplots show the diversity and population size ranges (thin lines) and interquartile ranges (thick lines) for each category. NA/DD indicates no IUCN Red List entry, or Red List status Data Deficient; LC is Least Concern, NT is Near Threatened, VU is Vulnerable, EN is Endangered, and CR is Critically Endangered.~~

#### Phylogenetic Non-Independence and the Population Size Diversity Relationship

~~In quantifying Lewontin's Paradox, I have additionally fit some simple trait evolution models that account for phylogenetic non-independence~~One limitation of using ordinary least squares is that shared phylogenetic history can create correlation structure in the residuals, which violates an assumption of the regression model (Felsenstein 1985; Revell 2010). To address this shortcoming, I

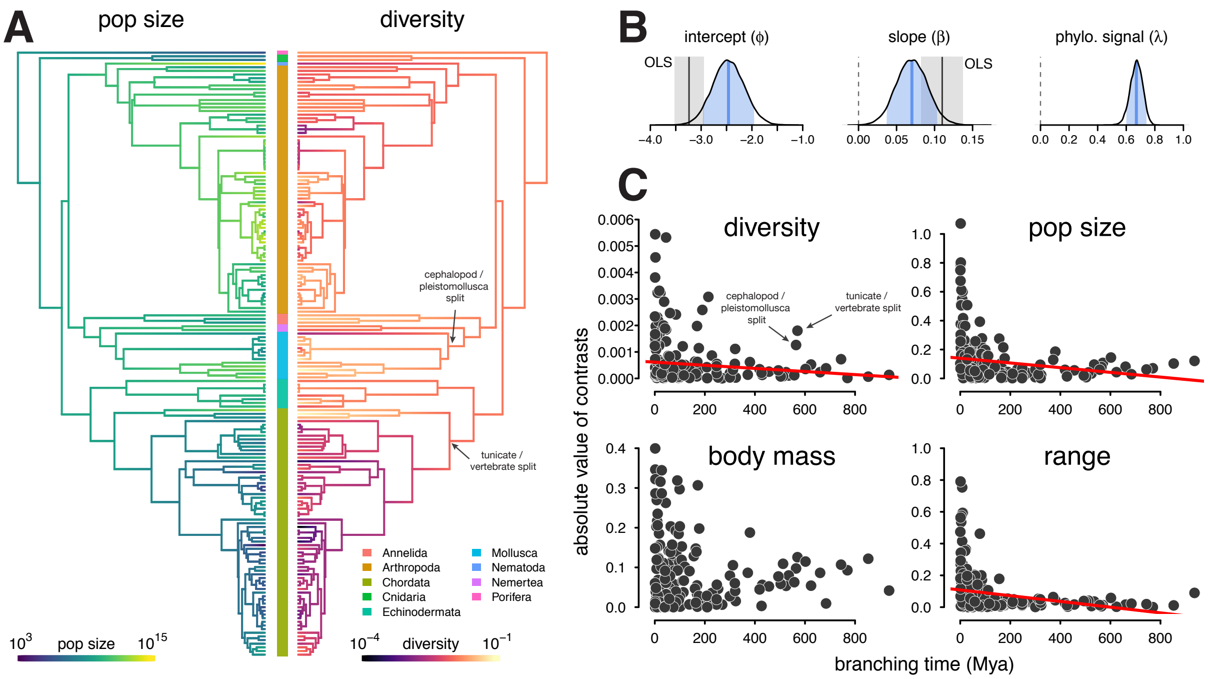


**Figure 3:** (A) The ancestral continuous trait estimates for the population size and diversity (differences per bp, log scaled) across the phylogeny of 166 taxa. The phyla of the tips are indicated by the color bar in the center. (B) The posterior distributions of the intercept, slope, and phylogenetic signal ($\lambda$, Villemereuil and Nakagawa 2014) of the phylogenetic mixed-effects model of diversity and population size (log scaled). Also shown are the 90% credible interval (light blue shading), posterior mean (blue line), OLS estimate (gray solid line), and bootstrap OLS confidence intervals (light gray shading). (C) The node-height tests of diversity, population size, and the two components of the population size estimates, body mass, and range (all traits on log scale before contrast was calculated). Each point shows the standardized phylogenetic independent contrast and branching time for a pair of lineages. Red lines are robust regression estimates (and are only shown for statistically significant relationships at the $\alpha = 0.05$ level). Note that some outlier pairs with very high phylogenetic independent contrasts were excluded (in all cases, these outliers were in the genus *Drosophila*).

fit this relationship using a phylogenetic mixed-effects model, investigated whether there is a signal of phylogenetic non-independence, estimated the continuous trait values on the phylogeny, and ~~assessed~~ explored how diversity and population size evolve. Prior population genetic comparative studies have lacked time-calibrated phylogenies and assumed unit branch lengths (Whitney and Garland 2010), a shortcoming that has drawn criticism (Lynch 2011). Here, I use a synthetic time-calibrated phylogeny created from the DateLife project (O'Meara et al. 2020) to account for shared phylogenetic history (see Methods: *Phylogenetic Comparative Methods*).

Using a phylogenetic mixed-effects model (Hadfield and Nakagawa 2010; Lynch 1991; Villemereuil and Nakagawa 2014) implemented in Stan (Carpenter et al. 2017; Stan Development Team 2020), I estimated the linear relationship between diversity and population size (on a log-log scale) accounting for phylogeny, for the 166 taxa ~~with non-missing~~ without missing data and present in the synthetic chronogram. As with the ~~non-phylogenetic~~ linear regression, this relationship was positive and significant (95% credible interval ~~0.04, 0.11~~0.03, 0.11), though somewhat attenuated compared to the OLS estimates (Figure ~~2~~3B). Since the population size estimates are based on range and body mass, they are essentially a composite trait; fitting phylogenetic mixed-effects models separately on body mass and range indicates these have significant negative and positive effects, respectively (Supplementary Figure S6; see also Supplementary Figure S1 for the relationship between diversity and the range categories of Leffler et al. 2012).

With the phylogenetic mixed-effects model, I also estimated the variance of the phylogenetic effect ($\sigma_p^2$) and the residual variance ($\sigma_r^2$), which can be used to estimate a measure of the phylogenetic signal, $\lambda = \sigma_p^2/(\sigma_p^2+\sigma_r^2)$ (Lynch 1991; Villemereuil and Nakagawa 2014; see Freckleton et al. 2002 for a comparison to Pagel's $\lambda$). If the relationship between diversity and population size was free of shared phylogenetic history, $\lambda = 0$ and all the variance could be explained by evolution on the tips; this is analogous to Lynch's conjecture that coalescent times should be free of phylogenetic signal (2011). In the relationship between population size and diversity, the posterior mean of $\lambda = 0.67$ (90% credible interval ~~[0.59, 0.75]~~[0.58, 0.75]) indicates that the majority of the variance perhaps might be due to shared phylogenetic history (Figure 3B).

~~A closer~~ This high degree of phylogenetic signal suggests Gillespie's (1991) concern that the $\pi$–$N_c$ relationship was driven by chordate-arthropod differences may be valid. A visual inspection of the estimated ancestral continuous values for diversity and population size on the phylogeny indicates the high phylogenetic signal seems to be driven in part by chordates having low diversity and small population sizes compared to non-chordates (Figure 3A). This ~~suggests Gillespie's (1991) earlier critique that the $\pi$–$N_c$ relationship was driven by chordate-arthropod differences may be valid. This~~ problem resembles Felsenstein's worst-case scenario (Felsenstein 1985; Uyeda et al. 2018), where a singular event on a lineage separating two clades generates a spurious association between two traits. ~~To further~~

To investigate whether clade-level differences dominated the relationship between diversity and population size, I fit phylogenetic mixed-effects models to phyla-level subsets of the data for clades with sufficient sample sizes (see Methods: *Phylogenetic Comparative Methods*). This analysis shows a significant positive relationship between diversity and population size in arthropods, and positive weak relationships in molluscs and chordates (Supplementary Figure S15). Each of the 90% credible intervals for slope overlap, indicating the relationship between $\pi$ and $N_c$ is similar across these clades.

~~One limitation of the phylogenetic mixed-effects models employed here is that they assume traits evolve under constant-rate Brownian motion. To test this assumption, I performed~~ Additionally, I have explored the rate of trait change through time using node-height tests (Freckleton and Harvey 2006). Node-height tests regress the absolute values of the standardized contrasts between lineages against the branching time (since present) of these lineages. Under Brownian Motion (BM), standardized contrasts are estimates of the rate of character evolution (Felsenstein 1985); if a trait evolves under constant rate BM, this relationship should be flat. For both diversity and population size, node-height tests indicate a significant increase in the rate of evolution towards the present (robust regression p-values ~~0.028 and 0.00070~~ 0.023 and 0.00018 respectively; Figure 3C). Considering the constituents of the population size estimate, range and body mass, separately, the rate of evolution of range but not body mass shows a significant increase (p-value ~~$1.9 \times 10^{-7}$) in rate~~ $1.03 \times 10^{-7}$) towards the present.

Parameterizing the model this way, I then set the key parameters that determine the impact of recurrent hitchhiking and background selection ($\gamma$, $J$, and $U$) to high values estimated from *Drosophila melanogaster* by Elyashiv et al. (2016). My estimate of the adaptive substitutions per generation ($\gamma_{\text{Dmel}}$) based Elyashiv et al. implies a rate of sweeps per basepair of $\nu_{\text{BP,Dmel}} \approx 2.34 \times 10^{-11}$, which is close to other estimates from *D. melanogaster* (see Supplementary Figure S14A). The rate of deleterious mutations per diploid genome, per generation is parameterized using the estimate from Elyashiv et al., $U_{\text{Dmel}} = 1.6$, which is a bit greater than previous estimates based on Bateman-Mukai approaches (Charlesworth 1987; Mukai 1988; Mukai 1985). Finally, the probability that a lineage is trapped in a sweep, $J_{\text{Dmel}}$, is calculated from the estimated genome-wide average coalescent rate due to sweeps from Elyashiv et al. (see Supplementary Figured S14B and Methods: *Predicted Reductions in Diversity* for more details on parameter estimates). Using these *Drosophila* parameters, I then explore how the predicted range of diversity levels under background selection and recurrent hitchhiking varies across species with recombination map length ($L$) and census population size ($N_c$).

Previous work has found that the impact of linked selection increases with $N_c$ (Corbett-Detig et al. 2015; see also Supplementary Figure S13A), and it is often thought that this is driven by higher rates of adaptive substitutions in larger populations (Ohta 1992), despite equivocal evidence (Galtier 2016). However, there is another mechanism by which species with larger population sizes might experience a greater impact of linked selection: recombinational map length, $L$, is known to correlate with body mass (Burt and Bell 1987) and thus varies inversely with population size. As this is a critical parameter that determines the genome-wide impact of both hitchhiking and background selection, I examine the relationship between recombination map length ($L$) and census population size ($N_c$) across taxa, using available estimates of map lengths across species (Corbett-Detig et al. 2015; Stapley et al. 2017). I find a significant non-linear relationship using phylogenetic mixed-effects models (Figure 4A; see Methods: *Phylogenetic Comparative Methods*). There is also a correlation between map length and genome size (Supplementary Figure S8) and genome size and population size (Supplementary Figure S7). These findings are consistent with both the hypothesis that non-adaptive processes increase genome size in small-$N_e$ species (Lynch and Conery 2003) which in turn could increase map lengths, as well as the hypothesis that map lengths are adaptively longer to more efficiently select against deleterious alleles (Roze 2021). Overall, the negative relationship between map length and census size indicates linked selection is expected to be stronger in short map length, high-$N_c$ species.

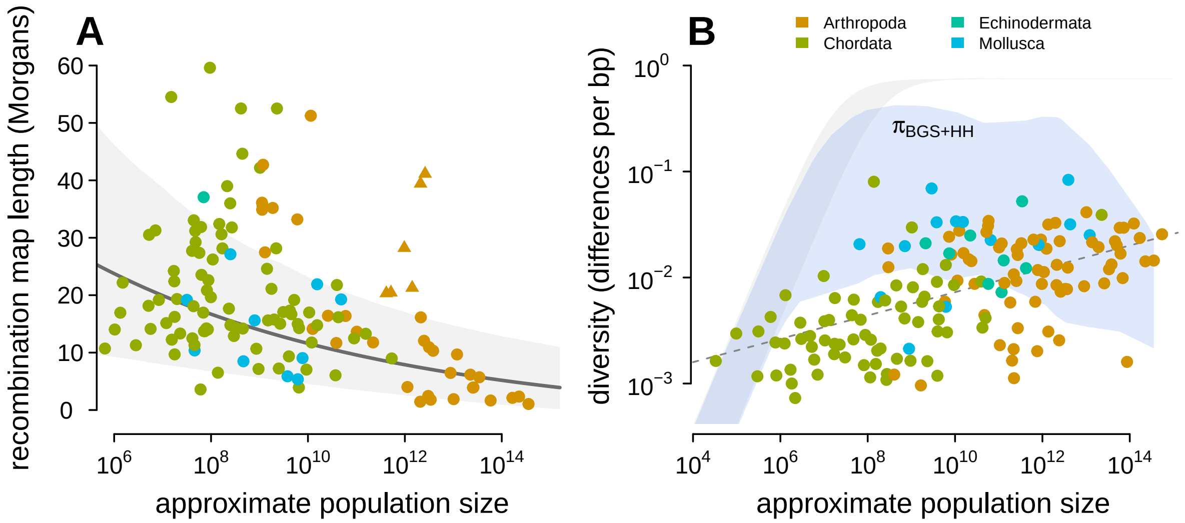


**Figure 4:** (A) The observed relationship between recombination map length ($L$) and census size ($N_c$) across ~~131~~ 136 species with complete data and known phylogeny. Triangle points indicate six social taxa excluded from the model fitting since these have adaptively higher recombination map lengths (Wilfert et al. 2007). The dark gray line is the estimated relationship under a phylogenetic mixed-effects model, and the gray interval is the 95% posterior average. (B) Points indicate the observed $\pi$–$N_c$ relationship across taxa shown in Figure 2, and the blue ribbon is the range of predicted diversity were $N_e = N_c$ for $\mu = 10^{-8}$–$10^{-9}$, and after accounting for the expected reduction in diversity due to background selection and recurrent hitchhiking under *Drosophila melanogaster* parameters. In both plots, point color indicates phylum.

### Discussion

Nearly fifty years after Lewontin's description of the Paradox of Variation, how evolutionary, life history, and ecological processes interact to constrain diversity across taxa to a narrow range remains a mystery. ~~Since Wright (1931; 1938), population geneticists have appreciated that various demographic processes shrink effective population sizes compared to census sizes, yet it has remained unclear whether these neutral processes alone can explain~~ I revisit Lewontin's Paradox ~~and across-taxa diversity patterns. Alternatively, selective processes that act more strongly in larger populations could account for the observed narrow range of diversity. A critical first step to discerning the processes that act to transform census sizes to diversity levels across species is characterizing the observed $\pi$–$N_c$ relationship .~~

~~Here, for 172 taxa, I estimated the relationship~~ by first characterizing the relationship between genomic estimates of pairwise diversity and approximate census population size across 172 metazoan species. Previous surveys have used allozyme-based estimates, fewer taxa, or qualitative measures of population size. My estimates of census population sizes are quite approximate, since they use body size to predict density. An improved estimate might consider vagility (as Soulé 1976 did), though this is harder to do systematically across many taxa. Future work might also use other ecological information, such as total biomass~~and estimated numbers of species within phyla,~~ , or species distribution modeling to improve census size estimates (Bar-On et al. 2018; Mora et al. 2011). Still, it seems more accurate estimates would be unlikely to change the qualitative findings here, which resemble those of early surveys (Nei and Graur 1984; Soulé 1976).

One limitation of the dataset in this study is that diversity estimates are collated from a variety of sources rather than estimated with a single bioinformatic pipeline. This leads to technical noise across diversity estimates; perhaps the relationship between $\pi$ and $N_c$ found here could be tighter with a standardized bioinformatic pipeline. In addition to this technical variation, there might be systematic bioinformatic sources of bias in diversity estimates. For example high-diversity sequences may fail to align to the reference genome and end up unaccounted for, leading to a downward bias. Alternatively, high-diversity sequences might map to the reference genome, but adjacent mis-matching SNPs might be mistaken for a short insertion or deletion. While these issues might adversely affect the estimates in high-diversity species, it is unlikely they will qualitatively change the observed $\pi$–$N_c$ relationship.

#### Macroevolution and Across-Taxa Population Genomics

Lewontin's Paradox arises from a comparison of diversity across species, yet ~~surprisingly, previous work on this problem has not considered the impact of phylogenetic non-independence. I have~~

addressed this limitation, showing that diversity does have a significant positive relationship with census size , after accounting for shared phylogenetic history among taxait has been disputed whether such comparisons require phylogenetic comparative methods. Extending previous work that has accounted for phylogeny in particular clades (Leffler et al. 2012), or using taxonomical-level averages (Romiguier et al. 2014), I show that the positive relationship between diversity and census size is significant using a mixed-effects model with a time-calibrated phylogeny. Additionally, I find a high degree of phylogenetic signal, evidence of deep shifts in the rate of evolution of genetic diversity, and that arthropods and chordates form clusters, showing previous concern . Overall, this suggests that previous concerns about phylogenetic non-independence was warranted . Finally, this high degree of phylogenetic signal, as well as evidence of shifts in the rate of evolution of genetic diversity on deep timescales in molluscs and chordates, seem to contradict Lynch 's in comparative population genetic studies were warranted (Gillespie 1991; Whitney and Garland 2010). Notably, Lynch (2011) claim that since has argued that PCMs for pairwise diversity are unnecessary, since mutation rate evolution is fast and thus free of phylogenetic inertia, sampling variance should exceed the variance due to phylogenetic shared history, and coalescent times are much less than divergence times, they are not affected by shared phylogenetic history. . Since my findings suggest PCMs are necessary in some cases, it is worthwhile to address these points.

One can reconcile my findings with Lynch's claim by considering what evolutionary, ecological, life history, and demographic causal factors determine coalescent rates across species, and how these factors evolve across deep timescales. Lynch's conjecture that coalescent times First, Lynch has correctly pointed out that while coalescent times are much less than divergence times and should be free of phylogenetic signal may be true shared history, the factors that determine coalescent times (e.g. mutation rates and effective population size) may not be (2011). In other words, coalescent times are free from phylogenetic shared history *were we to condition* on these causal factors that could be affected by shared phylogenetic history. In contrast, my My estimates of phylogenetic signal in diversity, by contrast, are not conditioned on these factors. Importantly, even "correcting for" phylogeny implicitly favors certain causal interpretations over others (Uyeda et al. 2018; Westoby et al. 1995). Future work could try to untangle what causal factors determine coalescent times across species, as well as how these factors evolve across macroevolutionary timescales. Second, it is a misconception that a fast rate of trait evolution necessarily reduces phylogenetic signal (Revell et al. 2008), and that if either or both variables in a regression are free of phylogenetic signal, PCMs are unnecessary (Revell 2010; Uyeda et al. 2018). The evidence of high phylogenetic signal found in this study suggests PCMs are needed, in part to avoid spurious results from phylogenetic pseudoreplication.

I have compared these to the qualitative range descriptions Leffler et al. (2012) (Supplementary Figure S10) and compared my $\alpha$-shape method to a subset of taxa with range estimates from IUCN Red List (Chamberlain 2020; IUCN 2020; Supplementary Figure S9). Each census population size is then estimated as the product of range and density.

$$\mathbb{E}(\pi) = \frac{\theta}{\theta + {}^{1}\!/_{B} + 2NS} \tag{A2}$$

$$\approx \frac{\theta}{{}^{1}\!/_{B} + 2NS} \tag{A3}$$

(cf. equation 1 Elyashiv et al. 2016, and equation 20 of Coop and Ralph 2012). The BGS component is given by Hudson and Kaplan (1995),

$$B(U, L) = N_e \exp\left(-\frac{U}{L}\right) \tag{A4}$$

and the hitchhiking component is

$$S = \frac{\nu_{\text{BP}}}{r_{\text{BP}}} J \tag{A5}$$

(cf. Coop and Ralph 2012 equation 20) where $J$ is the probability that two lineages coalesce down to one, given sweeps occur uniformly along the genome. Under this homogeneous sweep model, $J$ is

$$J = \int_0^L q_f(r)^2 dr \tag{A6}$$

where $q_f(r)$ is the approximate probability that a lineage is trapped by a sweep to frequency $f$ when it is $r$ recombination fraction away from this sweep (cf. Coop and Ralph 2012 equation 15).

Since I use *Drosophila melanogaster* parameter estimates from Elyashiv et al. (2016), I now reconcile their model's $S$ term with the simple model above. They estimate $S$ in *Drosophila melanogaster* using a composite likelihood model that considers hitchhiking and background selection simultaneously, using substitutions and stratifying by annotation. For a neutral position at site $x$, the coalescent rate due to sweeps is given by Elyashiv et al.'s equation 3,

$$S(x) = \frac{1}{T} \sum_{i_S} \alpha(i_S) \sum_{y \in a(i_S)} \int \exp(-r(x, y)\tau(s, N)) g(s|i_S) ds \tag{A7}$$

where $T$ is the number of generations that substitutions accrue, $i_S = 1, \ldots, I_S$ is the annotation class (e.g. exons, introns, UTRs), $\alpha(i_S)$ is the fraction of substitutions in annotation class $i_S$ that are beneficial, $a(i_S)$ is the set of all substitutions in annotation class $i_S$, $\tau(s, N)$ is the fixation time of a site with additive effect $s$, and $g(s|i_S)$ is the distribution of selection coefficients for annotation class $i_S$.

Note, that we can recover the model of Coop and Ralph (2012) from this expression. Suppose there is only one annotation class, and $\alpha$ fraction of substitutions are beneficial, and one selection coefficient $\bar{s}$, (i.e. $g(s) = \delta_0(s - \bar{s})$), then

$$S(x) = \frac{\alpha}{T} \sum_{y \in a} \exp(-r(x, y)\tau(\bar{s}, N)). \tag{A8}$$

Let the number of substitutions be $m := |a|$, and imagine their positions are uniformly distributed on a segment of length $G$ basepairs with the focal site is the middle at position $x = 0$. Then, each substitution $y$ is a random distance $l_y \sim U(-G/2, G/2)$ away from the focal site. Assuming the recombination rate is a constant $r_{\text{BP}}$ per basepair, and approximating the sum with an integral, we have,

$$\begin{aligned}
S &= \frac{\alpha}{T} \sum_{i=1}^{m} \mathbb{E}_{l_i} \left( \exp(-r_{\text{BP}} l_i \tau(\bar{s}, N)) \right) && \text{(A9)} \\
&= \frac{\alpha}{TG} \sum_{i=1}^{m} \int_0^G \exp(-r_{\text{BP}} \ell \tau(\bar{s}, N)) d\ell && \text{(A10)} \\
&= \frac{\alpha m}{TG} \int_0^G \exp(-r_{\text{BP}} \ell \tau(\bar{s}, N)) d\ell && \text{(A11)}
\end{aligned}$$

Using $u$-substitution with $r = \ell r_{\text{BP}}$ this simplifies to

$$S = \frac{\alpha m}{TGr_{\text{BP}}} \int_0^L \exp(-r\tau(\bar{s}, N))dr \tag{A12}$$

where $L = Gr_{\text{BP}}$.

To simplify this notation, note that the rate of adaptive substitutions per basepair per generation is $\nu_{\text{BP}} = {}^{\alpha m}/_{GT}$, so

$$S = \frac{\nu_{\text{BP}}}{r_{\text{BP}}} \int_0^L \exp(-r\tau(\bar{s}, N))dr \tag{A13}$$

This is analogous to the second term of Coop and Ralph (2012) equation 17, with $k = i = 2$ and $x = 1$ (e.g. conditioning on a sweep to fixation). Note that there appears to be a factor of two error in Elyashiv et al. (2016) compared to Coop and Ralph (2012); here I include the factor of two. Then,

$$S = \frac{\nu_{\text{BP}}}{r_{\text{BP}}} \underbrace{\int_0^L \exp(-2r\tau(\bar{s}, N))dr}_{J} \tag{A14}$$

where the integral is equal to $J$ (c.f. $J_{2,2}$ of equation 15 in Coop and Ralph 2012) since a simple model of $q_f(r) = f\exp(-2r\tau(s, N))$ and if we condition on fixation, $f = 1$. This expression is useful to generalize across species, since we know $N$ and $L$. Additionally, we have estimates of $\alpha$ and ${}^{m}/_{T}$ in *Drosophila* and other species. In Elyashiv et al, they consider the number of substitutions per generation in genic regions only; it should be noted that the number of coding basepairs varies little across species. For convenience, I define $\gamma = {}^{\alpha m}/_{T}$ as the number of adaptive substitutions per generation per entire genome, such that $S(\gamma, L, J) = {}^{\gamma}/_{L}\, J$ used in the main text. Using the estimates of $m \approx 4.5 \times 10^5$, $\alpha \approx 0.42$, and $T \approx 8.4 \times 10^7$ from the Supplementary Material of Elyashiv et al., I arrive at $\gamma \approx 0.00226$ adaptive substitutions per generation, per genome. For a $\approx 100$ megabase genome, this translates to a $\nu_{\text{BP}} \approx 2.34 \times 10^{-11}$, which is close to previous estimates (Supplementary Figure S14). For $J$, I use an empirical estimate calculated from the genome-wide average of the rate of coalescent events due to sweeps, from Supplementary Table S6 of Elyashiv et al. ($r_s = 2NS \approx 0.92$). This implies $J \approx 4.46 \times 10^{-4}$. Alternatively, I have tried using the estimated distribution of selection coefficients from Elyashiv et al., but this led to a weaker estimate of $J$, since the adaptive substitutions considered tend to cluster around genic regions. Note that these *Drosophila* sweep parameters I have used are close to previous estimates (Supplementary Figures S14 A and B).

| | | Bar-On et al. | | Present study | | | | | |
|---|---|---|---|---|---|---|---|---|---|
| phylum | total species ($T$) | biomass ($B$) | prop. biomass | biomass ($b$) | prop. biomass | num. species ($n$) | factor overrepresented | prop. total species ($f = {}^{n}/_{T}$) | factor (${}^{b}/_{fB}$) |
| Arthropoda | $1.26 \times 10^{6}$ | 1.20 | 0.4635 | $2.80 \times 10^{-4}$ | 0.0102 | 68 | 0.02 | $5.41 \times 10^{-5}$ | 4.31 |
| Chordata | $5.41 \times 10^{4}$ | 0.87 | 0.3357 | $2.67 \times 10^{-2}$ | 0.9715 | 68 | 2.89 | $1.26 \times 10^{-3}$ | 24.40 |
| Annelida | $1.70 \times 10^{4}$ | 0.20 | 0.0772 | $1.23 \times 10^{-5}$ | 0.0004 | 3 | 0.01 | $1.76 \times 10^{-4}$ | 0.35 |
| Mollusca | $9.54 \times 10^{4}$ | 0.20 | 0.0772 | $4.56 \times 10^{-4}$ | 0.0166 | 13 | 0.21 | $1.36 \times 10^{-4}$ | 16.70 |
| Cnidaria | $1.60 \times 10^{4}$ | 0.10 | 0.0386 | $3.07 \times 10^{-5}$ | 0.0011 | 2 | 0.03 | $1.25 \times 10^{-4}$ | 2.45 |
| Nematoda | $2.50 \times 10^{4}$ | 0.02 | 0.0077 | $4.03 \times 10^{-6}$ | 0.0001 | 1 | 0.02 | $4.00 \times 10^{-5}$ | 5.03 |

**Table S1:** How the total carbon biomass estimates by phylum from Bar-On et al. (2018) compare to the implied biomass estimates from this study. All biomass estimates are carbon biomass, and the proportions are of total biomass with respect to the study. The proportion of biomass in this study compared to the Bar-On et al. estimates Bar-On et al. (2018) indicates chordates are overrepresented and arthropods are underrepresented in the present study; the factor that each phylum is overrepresented is given in the eighth column. Total species by phylum estimates are from Chapman et al. (2009), Nicol (1969), Reaka-Kudla et al. (1996), and Zhang (2013). The ratio column is the ratio of total biomass implied by the $N_c$ estimates of each species in a phylum to the actual biomass of that phylum.

| | mean | 2.5 % | 97.5 % |
|---|---|---|---|
| $\beta_0$ | ~~-2.70~~ -2.80 | ~~-3.00~~ -3.20 | ~~-2.40~~ -2.50 |
| $\beta_{LC}$ | ~~-0.37~~ -0.39 | ~~-0.55~~ -0.57 | ~~-0.19~~ -0.21 |
| $\beta_{NT}$ | ~~-0.19~~ -0.22 | ~~-0.80~~ -0.83 | ~~0.41~~ 0.39 |
| $\beta_{VU}$ | ~~-0.31~~ -0.34 | ~~-0.81~~ -0.84 | ~~0.19~~ 0.16 |
| $\beta_{EN}$ | ~~-0.39~~ -0.40 | ~~-0.71~~ -0.73 | ~~-0.06~~ -0.07 |
| $\beta_{CR}$ | -0.03 | -0.65 | 0.59 |
| $\beta_{N_c}$ | ~~0.06~~ 0.08 | ~~0.04~~ 0.05 | ~~0.09~~ 0.11 |

**Table S2:** The regression estimates of full IUCN Red List population size model for diversity, $\log_{10}(\pi) = \beta_0 + \beta_{LC}LC + \beta_{NT}NT + \beta_{VU}VU + \beta_{EN}EN + \beta_{CR}CR + \beta_{N_c}\log_{10}(N_c)$; *df* = 165. Using AIC to compare this full model to a reduced model of $\log_{10}(\pi) = \beta_0 + \beta_{N_c}\log_{10}(N_c)$, ~~$AIC_{\text{full}} = 203.99$~~ $AIC_{\text{full}} = 204.9$, ~~$AIC_{\text{reduced}} = 212.46$~~ $AIC_{\text{reduced}} = 216.4$.

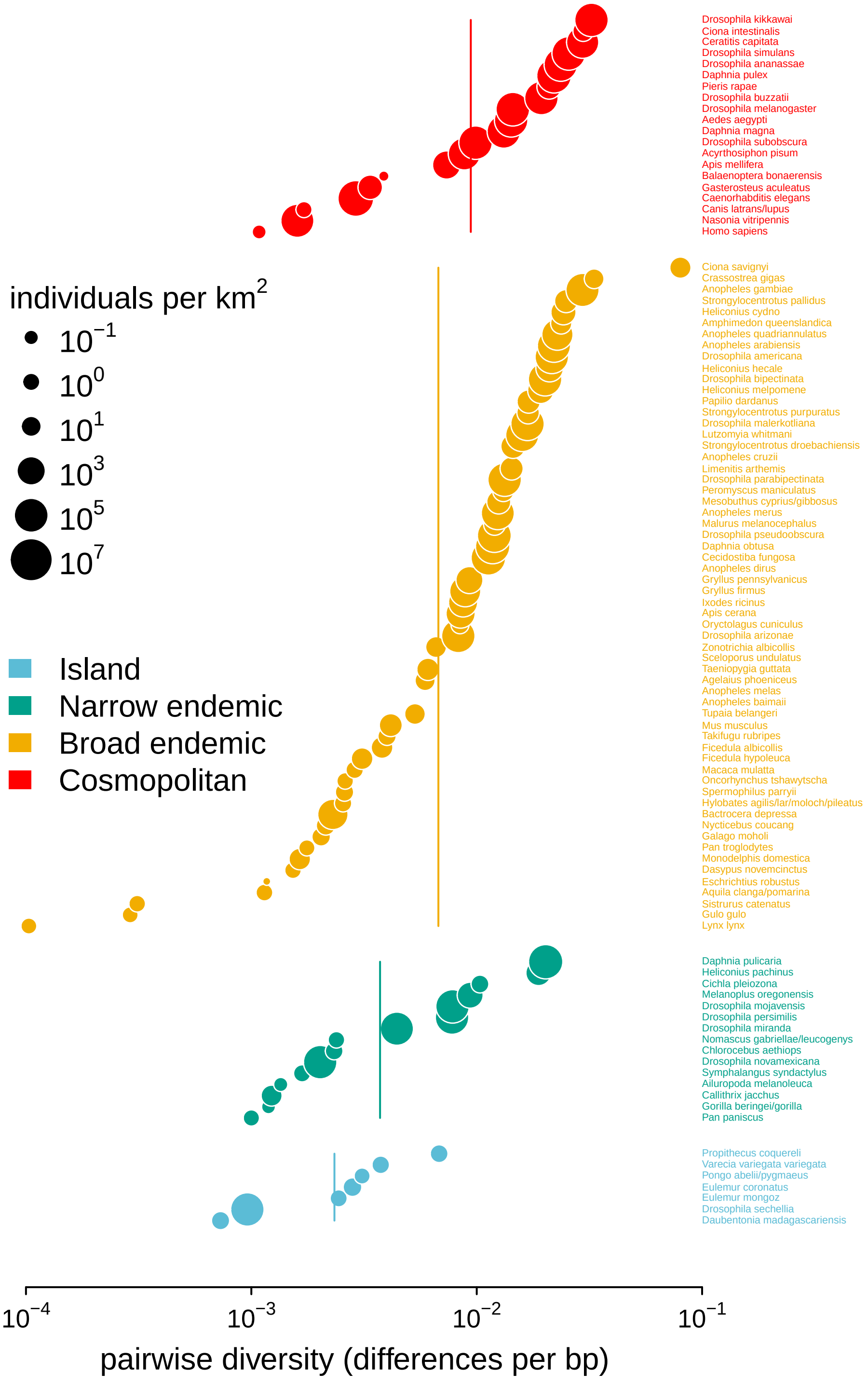


**Figure S1:** Pairwise diversity grouped by the range categories from Leffler et al. (2012), with point size indicating the predicted population density. The vertical lines are the range category group means.

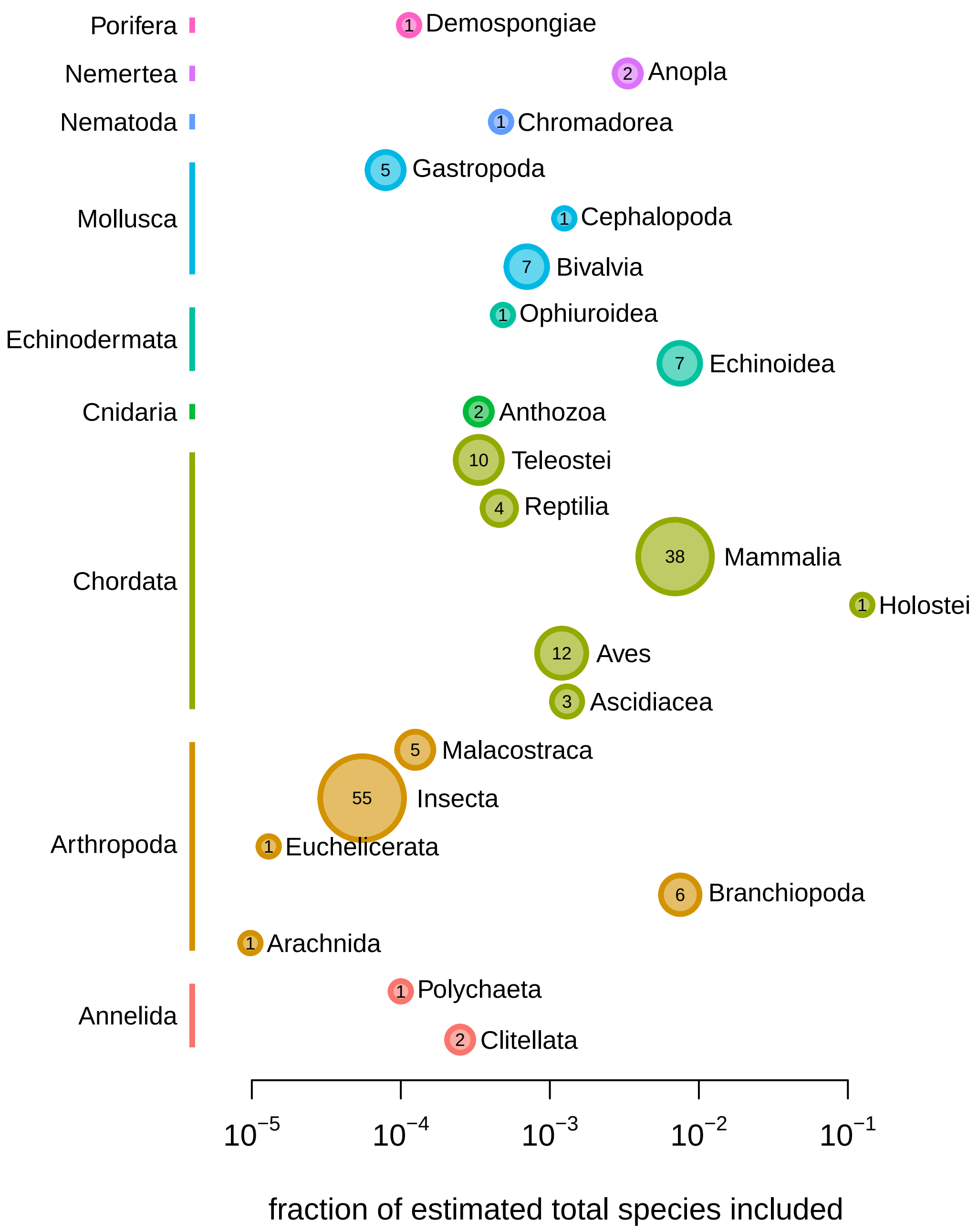


**Figure S2:** The fraction of total species on earth included in this study's sample, per class. The color of the points represents phylum, and the size of the point represents the absolute number of species by class.

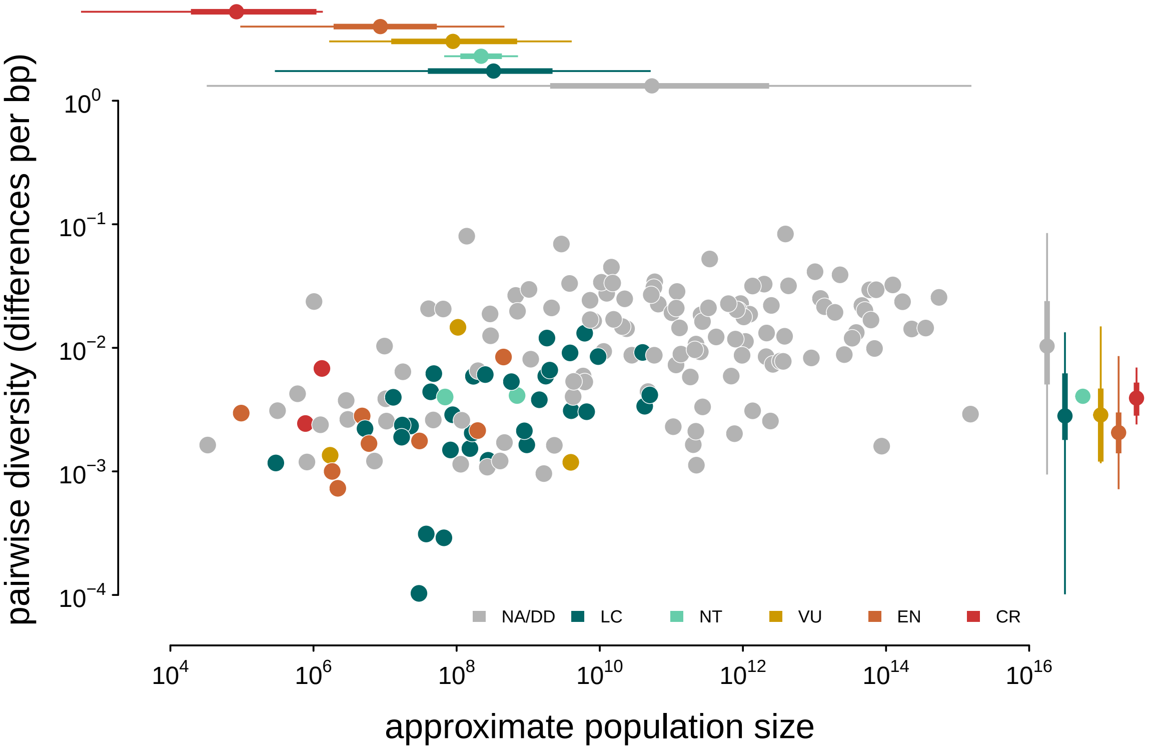


**Figure S3:** A version of Figure 2 with points colored by their IUCN Red List conservation status. Margin boxplots show the diversity and population size ranges (thin lines) and interquartile ranges (thick lines) for each category. NA/DD indicates no IUCN Red List entry, or Red List status Data Deficient; LC is Least Concern, NT is Near Threatened, VU is Vulnerable, EN is Endangered, and CR is Critically Endangered.

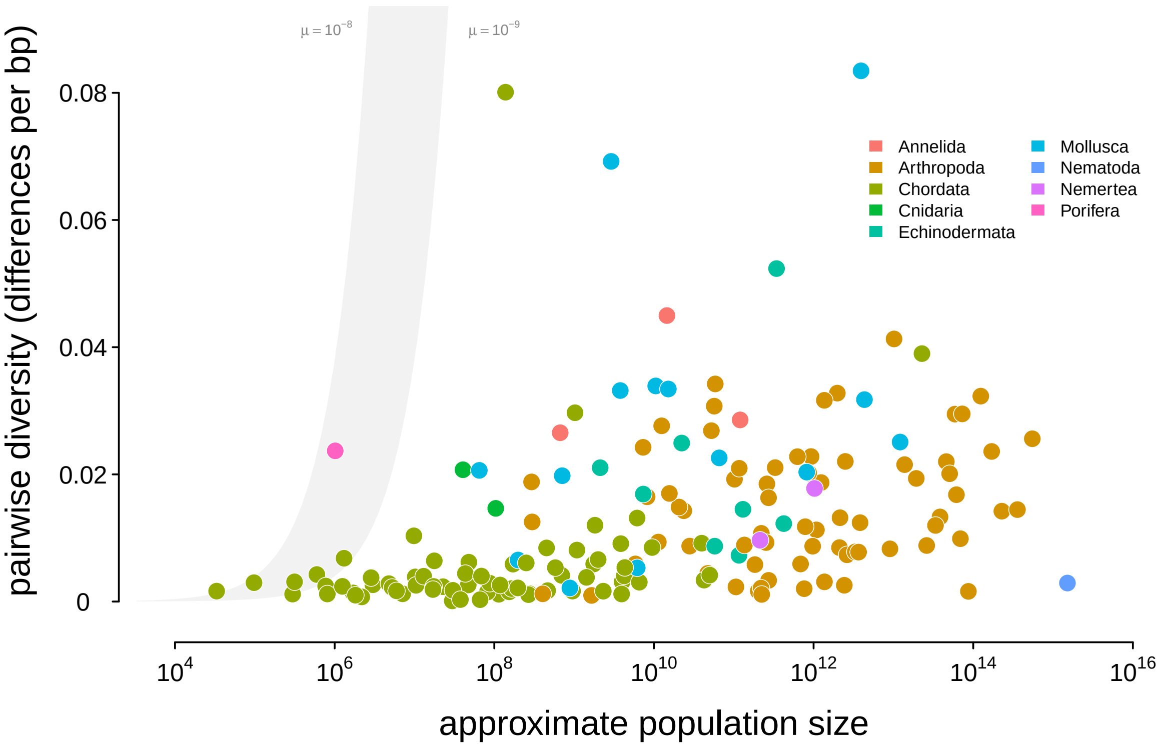


**Figure S4:** A version of Figure 2 with diversity on a linear, rather than log, scale. Points are colored by phylum, and the shaded region is the predicted neutral level of diversity assuming $N_e = N_c$ with mutation range ranging between $10^{-10} \leq \mu \leq 10^{-8}$.

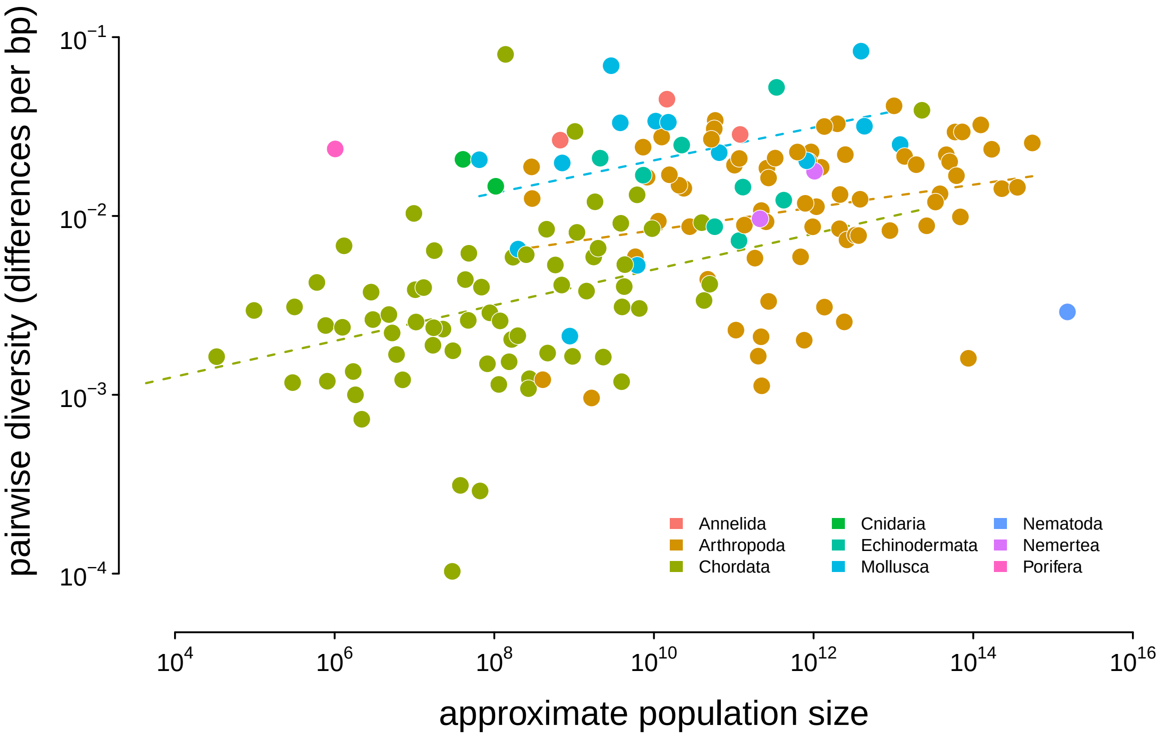


**Figure S5:** Diversity and approximate population size for 172 taxa, colored by phylum; the dashed lines indicate the non-phylogenetic OLS estimates of the relationship between population size and diversity grouped by phyla.

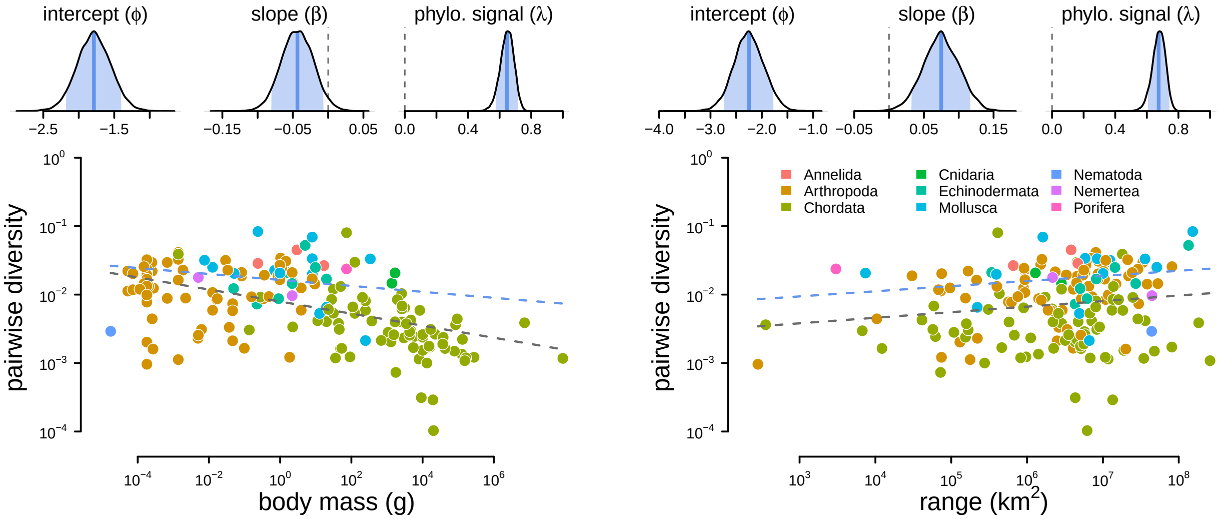


**Figure S6:** The relationship between diversity (differences per basepair) and body mass (left) and range (right) across 172 species. The top row are posterior distributions of parameters estimated using the phylogenetic mixed-effects model using 166 taxa in the synthetic phylogeny for the intercept, slope, and phylogenetic signal from the mixed-effects model. The bottom row contain each species as a point, colored by phyla. The gray dashed line is the non-phylogenetic standard regression estimate, and the blue dashed line is the relationship fit by the phylogenetic mixed-effects model.

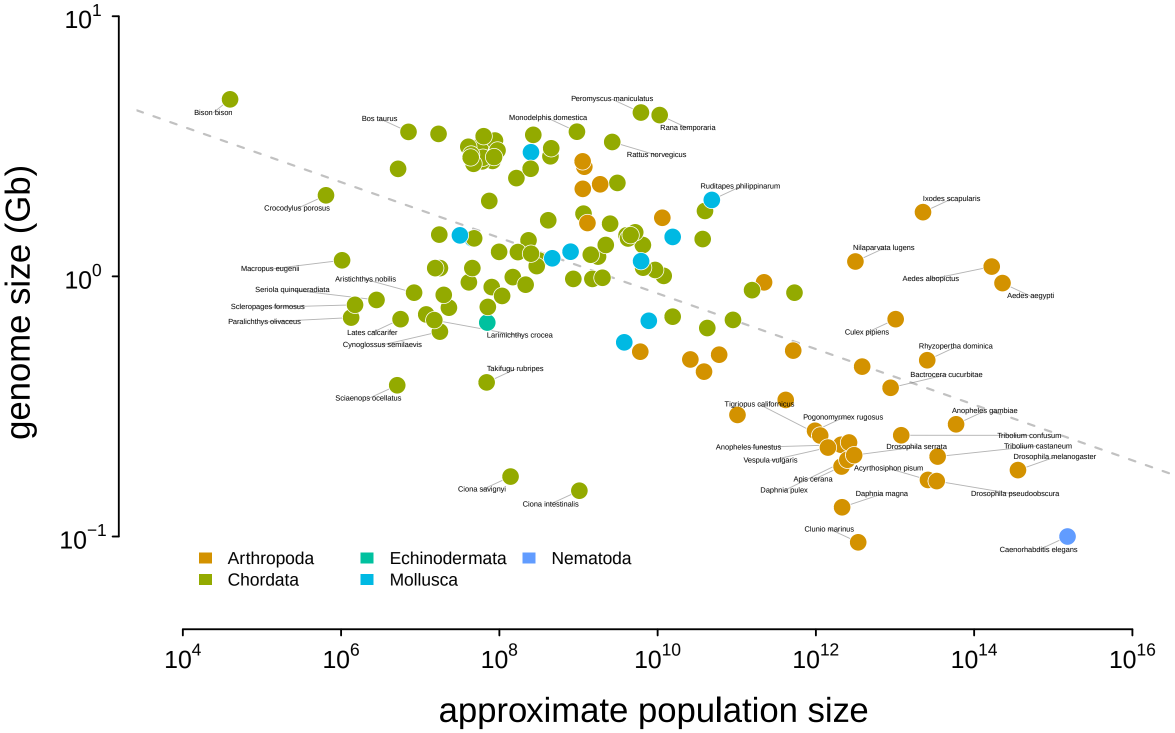


**Figure S7:** The relationship between genome size and approximate census population size. The dashed gray line indicates the OLS fit. Tiger salamander (*Ambystoma tigrinum*) was excluded because of its exceptionally large genome size ( 30Gbp).

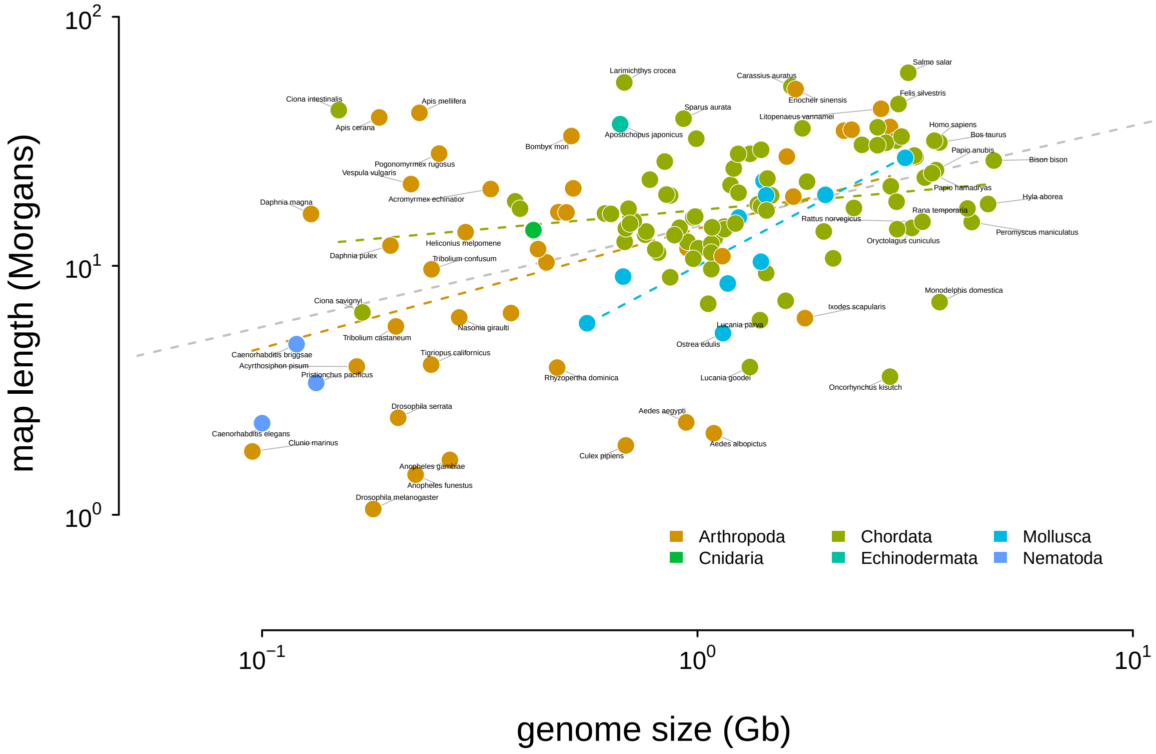


**Figure S8:** The relationship between genome size and recombination map length. The dashed gray line indicates the OLS fit for all taxa, and the dashed colored dashed lines indicate the linear relationship fit by phyla. Tiger salamander (*Ambystoma tigrinum*) was excluded because of its exceptionally large genome size ( 30Gbp).

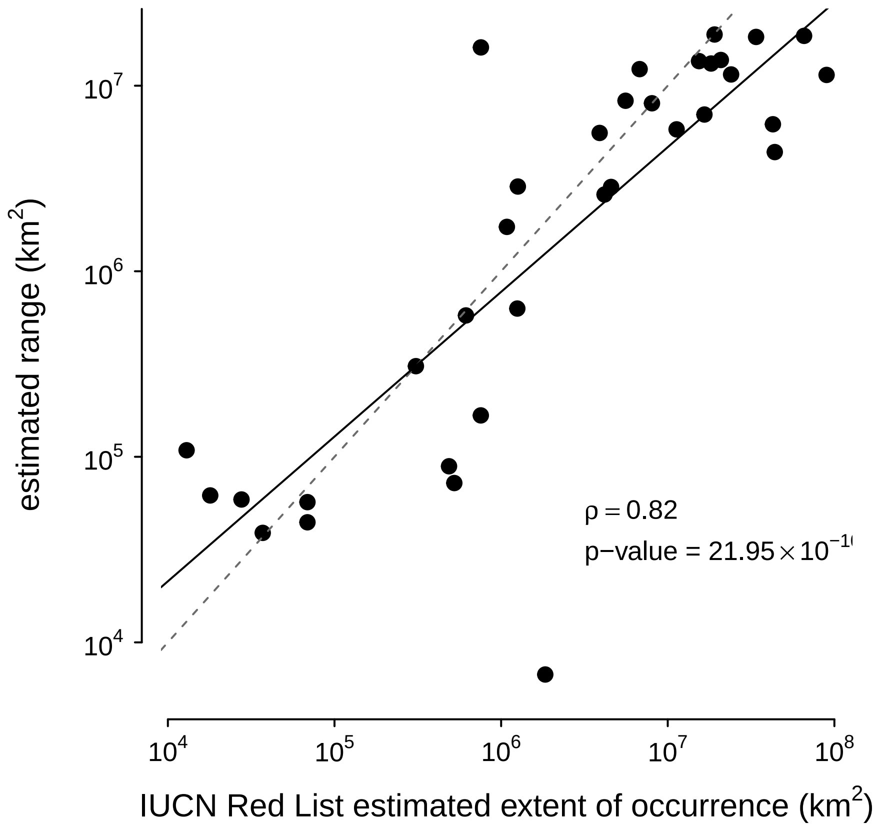


**Figure S9:** The correspondence between the ranges estimated with the alpha hull method applied to GBIF data used in this paper and IUCN Red List's Extent of Occurrence for the subset of species in both datasets. Note that the IUCN Red List contains predominantly endangered species, which leads to ascertainment bias; still, the high correlation between the estimated ranges shows the alpha hull method works well.

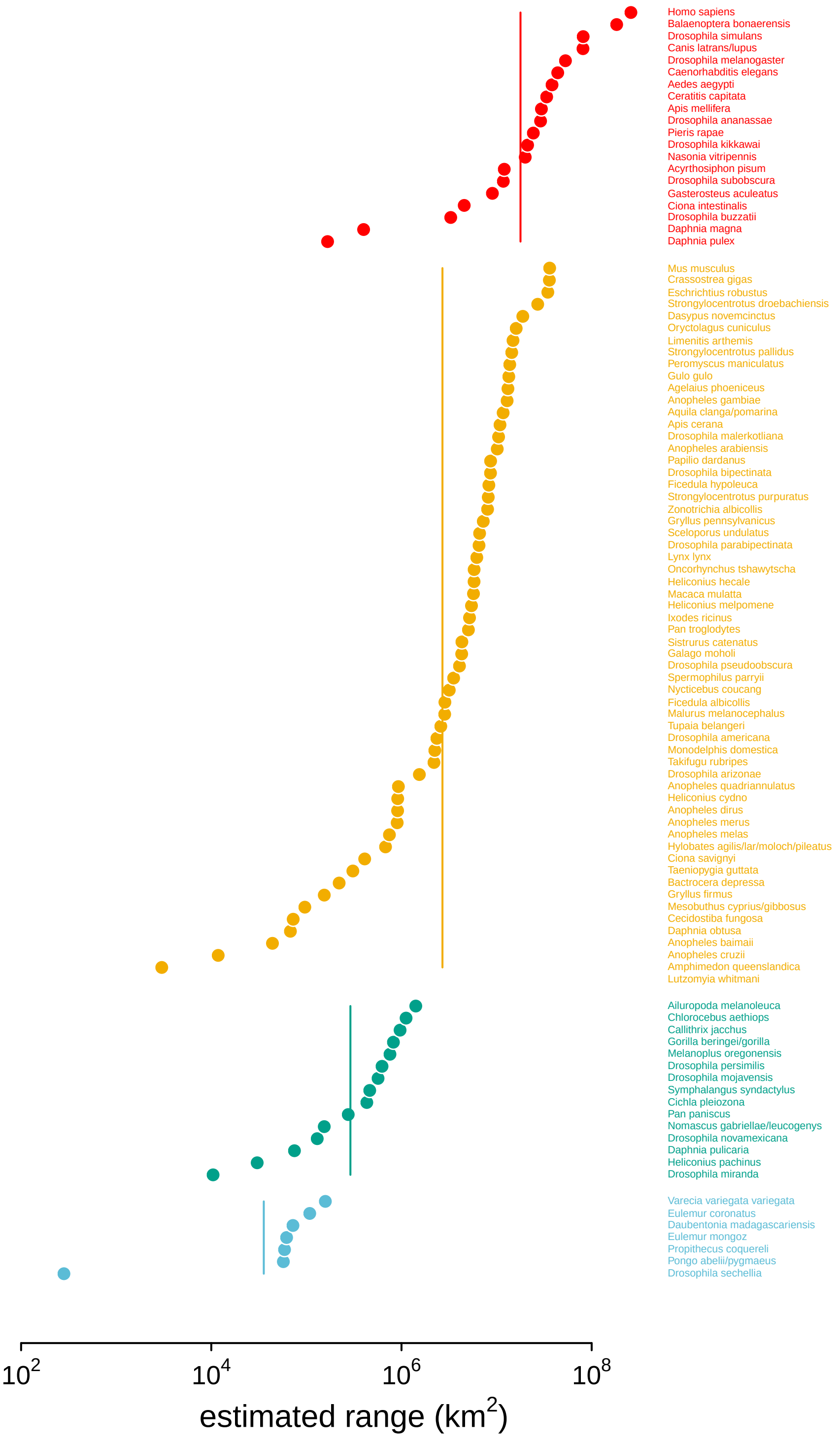


**Figure S10:** The estimated ranges using GBIF occurrence data, ordered within and colored by the original range category labels assigned in Leffler et al. (2012).

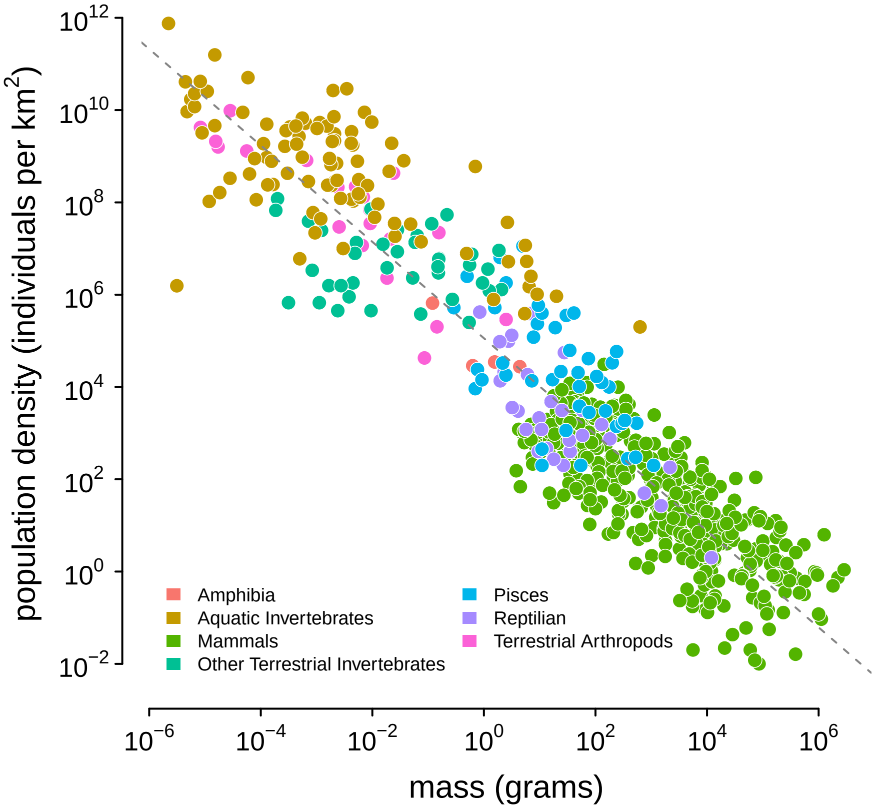


**Figure S11:** The appendix table of Damuth (1987); the color indicates Damuth's original group labels. The dashed line was estimated using a lognormal regression model in Stan. References to each measurement are available in Damuth (1987).

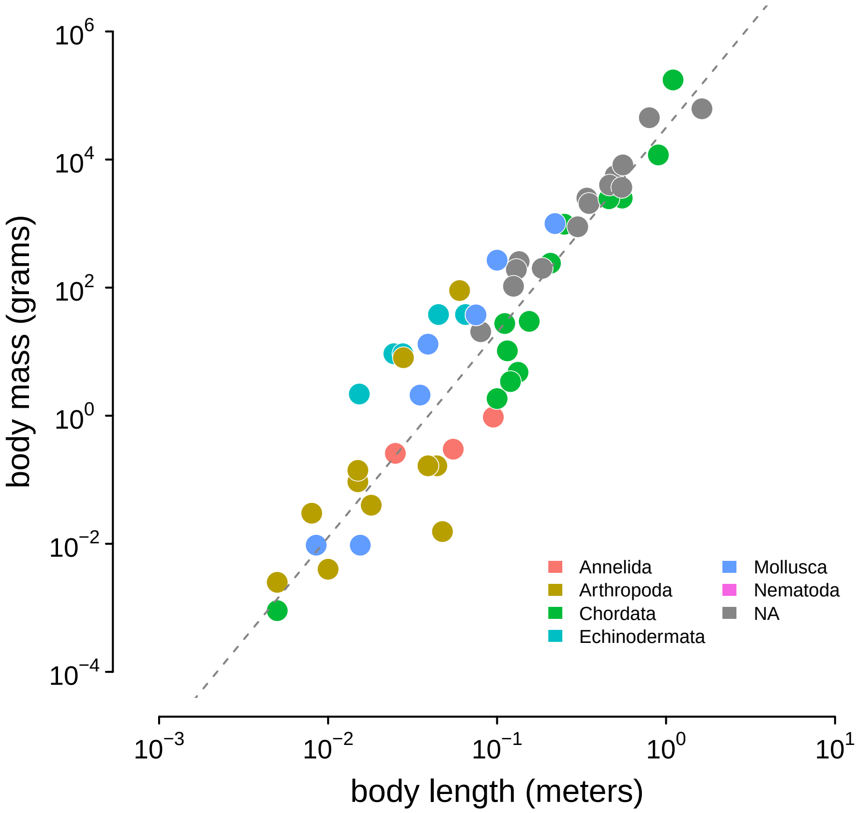


**Figure S12:** The relationship between body length (meters) and body mass (grams) in the Romiguier et al. (2014) data set, used to infer body masses for taxa. The gray dashed line is the line of best fit inferred using Stan.

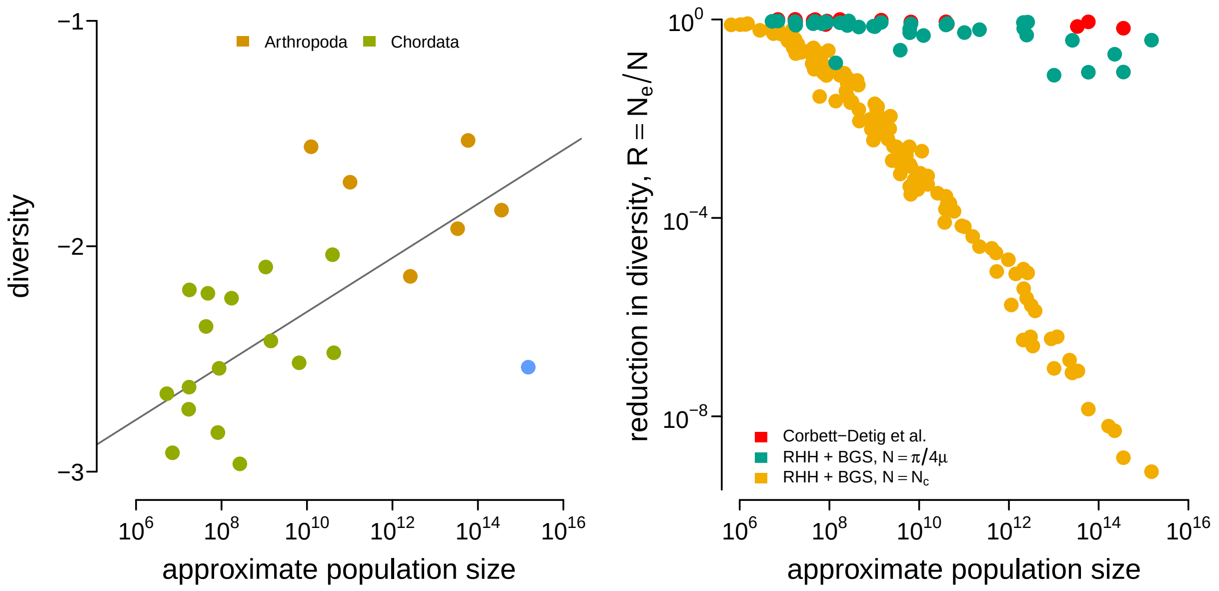


**Figure S13:** (A) The diversity data from Corbett-Detig et al. (2015) and the census population size estimated here for metazoan taxa. (B) The reductions in diversity, $R = {}^{N_e}/_{N}$, plotted against census size across species. The red points are the reductions estimated by Corbett-Detig et al. (2015). This confirms Corbett-Detig et al.'s (2015) finding that the impact of selection ($I = 1 - R$) increases with census population size (though, in the original paper size body size and range were used as separate proxy variables for census population size). The green and red points are the predicted reduction in diversity under the recurrent hitchhiking (RHH) and background selection (BGS) model using the *Drosophila melanogaster* parameters as described in the main text. The reduction in the diversity due to sweeps, from Equation (1), is determined by the term $2NS$. Green points treat $N$ as the implied effective population size from diversity $\widetilde{N}_e = {}^{\widehat{\pi}}/_{4\mu}$, assuming $\mu = 10^{-9}$. Yellow points treat $N$ as the census size, $N = N_c$. Overall, using the census size, e.g. $2N_cS$, leads to reductions in diversity that far exceed the empirical estimates of Corbett-Detig et al. and reasonable model-based predictions from $\widetilde{N}_e$.

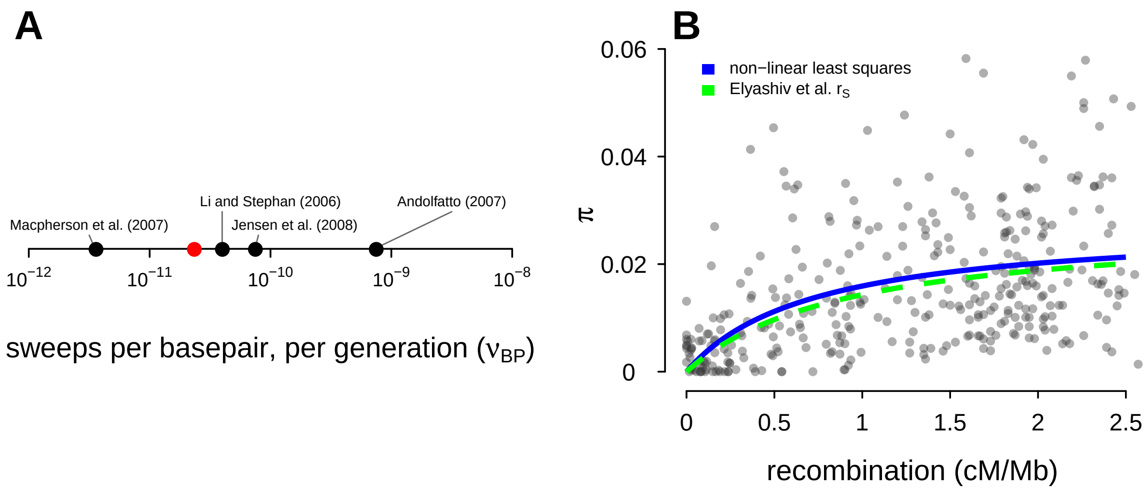


**Figure S14:** (A) The estimate of the number of sweeps per basepair, per genome ($\nu_{\mathrm{BP}}$) from Table 2 of Elyashiv et al. (2016) (the studies included are Andolfatto 2007; Li and Stephan 2006; Macpherson et al. 2007 and Jensen et al. 2008); the red point is my estimate used in this paper. (B) Points are the data from Shapiro et al. (2007). The blue line is the non-linear least squares fit to the data, and the green dashed line is the sweep model parameterized by the genome-wide average sweep coalescent rate $2NS \approx 0.92$ from the classic sweep and background selection model of Elyashiv et al. (2016) ($r_s$ in Supplementary Table S6).

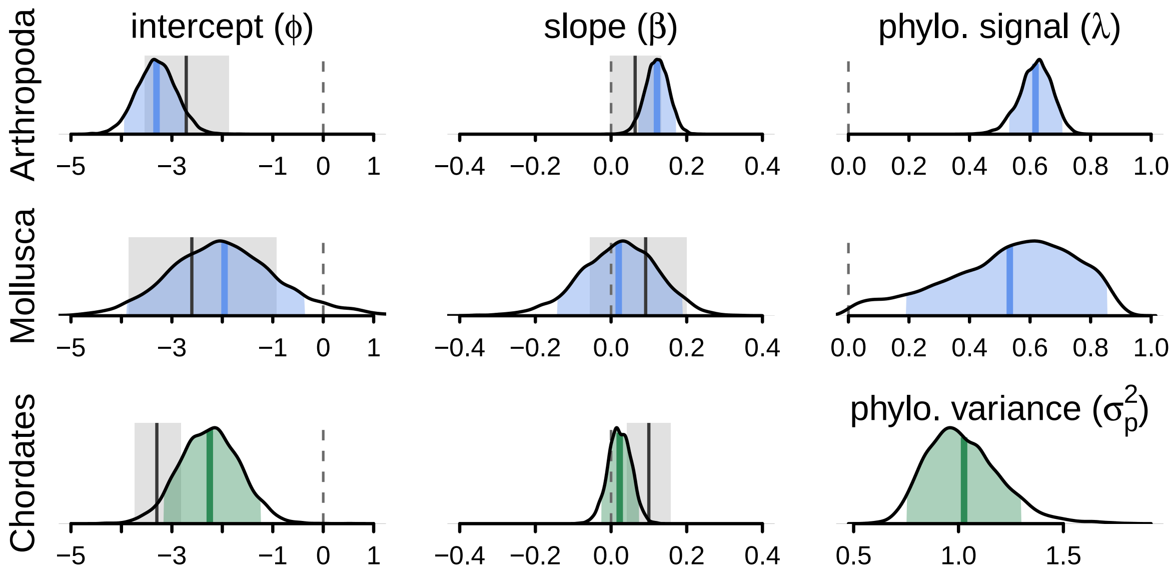


**Figure S15:** The posterior distributions for the parameters of the phylogenetic mixed-effects model of diversity and population size (this is analogous to Figure 3B) fit separately on chordates ($n = 68$), molluscs ($n = 13$), and arthropods ($n = 68$). The phylogenetic mixed-effects model for chordates indicated the best-fitting model had no residual variance ($\sigma_r^2 = 0$), so an alternate model without this variance component was used to ensure proper convergence; this model is shown in green. The light blue (green) shaded regions are the 90% credible intervals, the blue (green) lines the posterior averages, the gray shaded regions the OLS bootstrap 95% confidence intervals, and the gray lines the OLS estimate. Note that unlike Figure 3, the OLS estimate uses all taxa, not just those present in the phylogeny, since splitting the data by phyla reduces sample sizes (OLS with just the subset of taxa in the phylogeny is not significant for either chordates and arthropods). The vertical dashed gray line indicates zero.

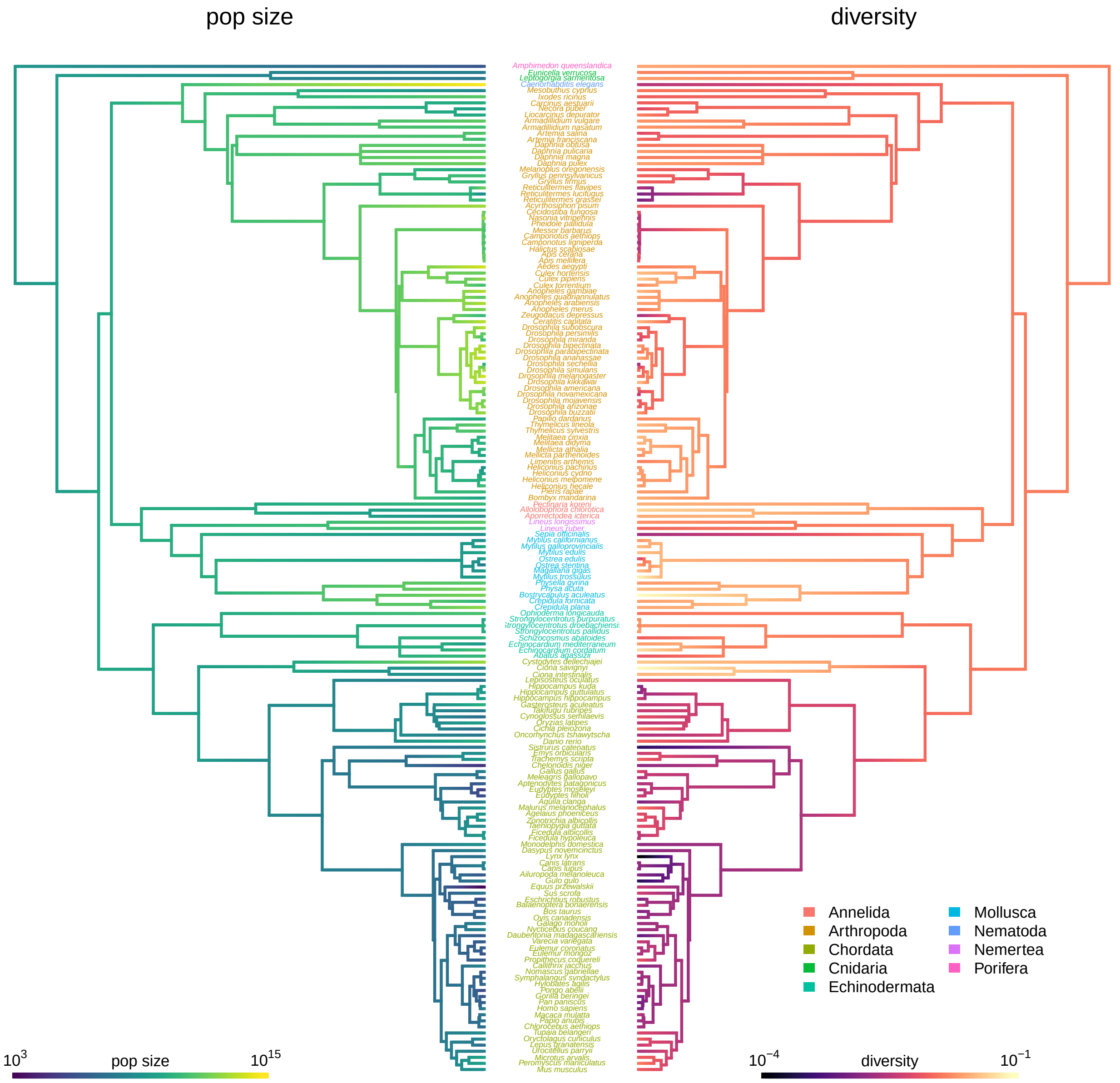
pop size
diversity
Annelida
Arthropoda
Chordata
Cnidaria
Echinodermata
Mollusca
Nematoda
Nemertea
Porifera
$10^{3}$
pop size
$10^{15}$
$10^{-4}$
diversity
$10^{-1}$

**Figure S16**

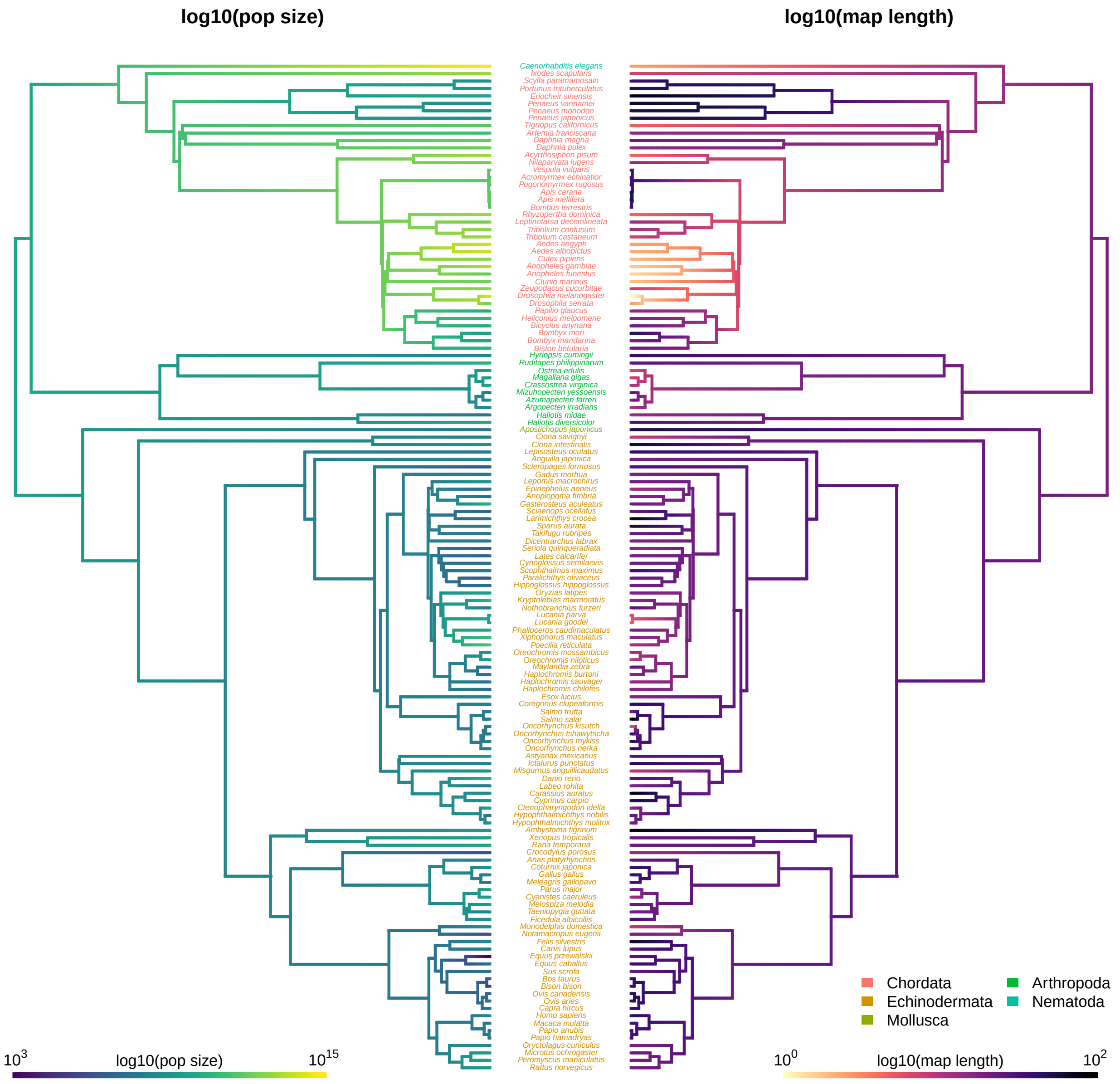
log10(pop size)
log10(map length)
Caenorhabditis elegans
Ixodes scapularis
Scylla paramamosain
Portunus trituberculatus
Eriocheir sinensis
Penaeus vannamei
Penaeus monodon
Penaeus japonicus
Tigriopus californicus
Artemia franciscana
Daphnia magna
Daphnia pulex
Acyrthosiphon pisum
Nilaparvata lugens
Vespula vulgaris
Acromyrmex echinatior
Pogonomyrmex rugosus
Apis cerana
Apis mellifera
Bombus terrestris
Rhyzopertha dominica
Leptinotarsa decemlineata
Tribolium confusum
Tribolium castaneum
Aedes aegypti
Aedes albopictus
Culex pipiens
Anopheles gambiae
Anopheles funestus
Clunio marinus
Zeugodacus cucurbitae
Drosophila melanogaster
Drosophila serrata
Papilio glaucus
Heliconius melpomene
Bicyclus anynana
Bombyx mori
Bombyx mandarina
Biston betularia
Hyriopsis cumingii
Ruditapes philippinarum
Ostrea edulis
Magallana gigas
Crassostrea virginica
Mizuhopecten yessoensis
Azumapecten farreri
Argopecten irradians
Haliotis midae
Haliotis diversicolor
Apostichopus japonicus
Ciona savignyi
Ciona intestinalis
Lepisosteus oculatus
Anguilla japonica
Scleropages formosus
Gadus morhua
Lepomis macrochirus
Epinephelus aeneus
Anoplopoma fimbria
Gasterosteus aculeatus
Sciaenops ocellatus
Larimichthys crocea
Sparus aurata
Takifugu rubripes
Dicentrarchus labrax
Seriola quinqueradiata
Lates calcarifer
Cynoglossus semilaevis
Scophthalmus maximus
Paralichthys olivaceus
Hippoglossus hippoglossus
Oryzias latipes
Kryptolebias marmoratus
Nothobranchius furzeri
Lucania parva
Lucania goodei
Phalloceros caudimaculatus
Xiphophorus maculatus
Poecilia reticulata
Oreochromis mossambicus
Oreochromis niloticus
Maylandia zebra
Haplochromis burtoni
Haplochromis sauvagei
Haplochromis chilotes
Esox lucius
Coregonus clupeaformis
Salmo trutta
Salmo salar
Oncorhynchus kisutch
Oncorhynchus tshawytscha
Oncorhynchus mykiss
Oncorhynchus nerka
Astyanax mexicanus
Ictalurus punctatus
Misgurnus anguillicaudatus
Danio rerio
Labeo rohita
Carassius auratus
Cyprinus carpio
Ctenopharyngodon idella
Hypophthalmichthys nobilis
Hypophthalmichthys molitrix
Ambystoma tigrinum
Xenopus tropicalis
Rana temporaria
Crocodylus porosus
Anas platyrhynchos
Coturnix japonica
Gallus gallus
Meleagris gallopavo
Parus major
Cyanistes caeruleus
Melospiza melodia
Taeniopygia guttata
Ficedula albicollis
Monodelphis domestica
Notamacropus eugenii
Felis silvestris
Canis lupus
Equus przewalskii
Equus caballus
Sus scrofa
Bos taurus
Bison bison
Ovis canadensis
Ovis aries
Capra hircus
Homo sapiens
Macaca mulatta
Papio anubis
Papio hamadryas
Oryctolagus cuniculus
Microtus ochrogaster
Peromyscus maniculatus
Rattus norvegicus
Chordata
Echinodermata
Mollusca
Arthropoda
Nematoda
$10^3$
log10(pop size)
$10^{15}$
$10^0$
log10(map length)
$10^2$

**Figure S17**
